# All Together Now: Joint Codon Alignment, Ohmic Networks, and Error-Controlled Selection Inference in HyPhy 3

**DOI:** 10.64898/2026.11.06.764988

**Authors:** Sergei L. Kosakovsky Pond, Steven Weaver, Darren P. Martin, Simon D. W. Frost, Gallean Brown, Danielle Callan, Daria Chalkova, David Enard, Vincent J. Lynch, Timothy B. Sackton, Hannah Verdonk, Jordan D. Zehr, Spencer V. Muse, Anton Nekrutenko

## Abstract

Codon-based maximum likelihood models are the workhorse of molecular selection analysis, yet standard workflows assume that the input sequence alignment and phylogenetic tree are known without error. In practice, heuristic aligners ignore the genetic code and phylogeny, which packs non-homologous codons into shared columns across insertion-deletion loops and inflates gene-wide false-positive rates above 80%. This sequential division of labor is statistically suboptimal and computationally wasteful. Here, we present WALRUS (**W**heatstone **A**lignment and **L**ikelihood **R**ate **U**nification for **S**election), a unified engine in HyPhy 3 (https://github.com/veg/hyphy-3) that maps unaligned coding sequences to a coordinate-consistent selection atlas in one run. From a single Inside–Outside likelihood table, the pipeline repairs reading frames, co-estimates a codon alignment and Ohmic reticulate network, models directional base-composition drift, and evaluates eight selection tests (BUSTED-E, aBSREL, RELAX, FEL, dFEL, MEME, Contrast-FEL, and PRIME). Across 10,020 simulated alignments, WALRUS recovers 94.6% of true columns on the 800 INDELible replicates and cuts indel-driven false positives from 44.7%– 58.1% to 4.4% under BUSTED (21.1%– 21.4% to 1.4% under BUSTED-E). Under Zipfian indel saturation, false positives drop four-fold (33.3%– 83.3% to 17.0%– 20.5%), while 11, 190 avian orthogroup runs execute 6.6–35.8-fold faster and cut housekeeping false positives 13-fold (28-fold on ultraconserved genes) proteome-wide.

## 1 Introduction: When Convenient Assumptions Become Positively Misleading

In 1976, George Box famously observed that because all statistical models are wrong, the scientist must remain alert to what is importantly wrong. Over the past three decades, probabilistic models of coding sequence evolution have grown from the foundational workhorse formulations of Muse and Gaut (1994) and Goldman and Yang (1994), which estimated a single gene-wide non-synonymous to synonymous rate ratio (*ω* = *d_N_ /d_S_*), into a hierarchy of hypothesis tests operating at every evolutionary scale (Kosakovsky Pond and Frost, 2005; Yang et al., 2000a). At the whole-gene level, random-effects branch-site models evaluate whether any non-empty fraction of codons along any subset of branches experienced *ω >* 1 (BUSTED, Murrell et al. 2015, and error-protected BUSTED-E, Selberg et al. 2025), whereas lineage- and clade-level tests identify specific branches undergoing episodic diversification (aBSREL, Smith et al. 2015) or quantify whether selection intensity has relaxed or intensified across a focal clade (RELAX, Wertheim et al. 2015). At the resolution of individual codons, fixed- and mixed-effects likelihood tests estimate site-specific synonymous (*α_s_*) and non-synonymous (*β_s_*) rates across the phylogeny (FEL, Kosakovsky Pond and Frost 2005), detect episodic selection restricted to branch subsets (MEME, Murrell et al. 2012), compare site-level rates among predefined branch partitions (Contrast-FEL, Kosakovsky Pond et al. 2021), and measure physicochemical amino-acid constraints (PRIME, Conant et al. 2007; Delport et al. 2010). When the input alignment and tree are free of upstream errors, these likelihood tests are well calibrated.

Although all of these methods evaluate continuous-time Markov chains over the *S* = 61 sense codons, comparative coding-sequence analyses are still executed in practice as a fragile chain of disconnected programs. An investigator typically downloads unaligned sequences from a public repository, builds a heuristic multiple sequence alignment with MAFFT (Katoh and Standley, 2013), PRANK (Löytynoja and Goldman, 2008), or MACSE (Ranwez et al., 2011, 2018), infers a single point-estimate tree in IQ-TREE (Minh et al., 2020) or RAxML-NG (Kozlov et al., 2019), optionally screens for recombination with GARD (Kosakovsky Pond et al., 2006), and finally runs six to eight standalone HyPhy selection tools (Kosakovsky Pond et al., 2020; Pond et al., 2005). Conditioning downstream selection inference on a fixed upstream alignment and tree was a computationally convenient assumption in 1994, when a typical dataset contained ten sequences analyzed on a desktop workstation. On modern comparative datasets spanning hundreds of divergent species, however, this sequential pipeline becomes positively misleading (Wong et al., 2004). Heuristic aligners optimize amino-acid or nucleotide similarity scores without a phylogenetic tree or codon model, so in surface loops where insertions and deletions (indels) are common, they routinely pack independent insertions into shared columns or shift homologous codons across gap boundaries (Fletcher and Yang, 2010; Jordan and Goldman, 2012; Löytynoja and Goldman, 2008). Because Felsenstein pruning treats gaps as missing data carrying zero cost while penalizing nucleotide mismatches, even a one-codon register shift creates an apparent burst of non-synonymous substitutions, inflating gene-wide false-positive rates above 80% (Selberg et al., 2025).

Compounding this upstream alignment fragility, standard single-hit stationary codon models ignore three pervasive biological processes that routine genome screens encounter at scale. Specifically, Wisotsky et al. (2020) demonstrated that ignoring site-to-site synonymous rate variation conflates silent rate depression with elevated *ω*, while Lucaci et al. (2023) showed that instantaneous multi-nucleotide substitutions (two- and three-hit events, 2H/3H) force single-hit models through spurious non-synonymous intermediates on short branches. Across deep vertebrate and insect clades, GC-biased gene conversion and mutational skews also drive large shifts in third-codon-position GC content (GC_3_; Duret and Galtier 2009; Ratnakumar et al. 2010), violating the stationary equilibrium assumed by reversible codon matrices. Ignore these confounding processes at your own peril. When unmodeled rate variation, multi-nucleotide hits, and alignment boundary errors combine across thousands of genes, the imprimatur of positive selection loses its luster because genuine adaptive signals are buried under a deluge of false positives. Indeed, many comparative genomicists and experimental biologists have grown reticent to undertake molecular selection analyses at all: assembling frame-clean coding alignments and guarding against every upstream technical and mutational confounder requires so much specialized manual intervention that the barrier to trustworthy inference often appears prohibitive. At the same time, running FEL, dFEL, MEME, BUSTED, aBSREL, RELAX, Contrast-FEL, and PRIME as separate standalone executables repeats baseline nucleotide and codon fits, matrix diagonalizations, and Felsenstein tree traversals (Felsenstein, 1981) thousands of times on the same data, while forcing researchers to reconcile incompatible coordinate systems by hand whenever sequences are trimmed or realigned.

Here, we present WALRUS (**W**heatstone **A**lignment and **L**ikelihood **R**ate **U**nification for **S**election), a unified C++20 inference engine in HyPhy 3 that replaces this brittle multi-tool chain with a single coordinate-consistent pipeline, performing the upstream curation, frame-repairing codon alignment, and confounder control automatically for the user. By coupling curation, alignment, tree estimation, and selection testing around a single precomputed Inside–Outside likelihood table, the engine eliminates both redundant tree traversals and post hoc coordinate mismatches. To make each modeling decision concrete, we first trace four worked examples (abalone sperm lysin, mammalian interphotoreceptor retinoid-binding protein *RBP3*, primate semenogelin-2 SEMG2, and mammalian tumor suppressor *TP53*) from raw coding sequences to aligned codons, structural domains, and clade regimes. We then detail the six-stage architecture, evaluate its operating characteristics across 10,020 simulated alignments spanning reading-frame errors, recombination, indels, synonymous rate variation, multi-nucleotide hits, compositional drift, and clade-level shifts, and apply it across 11, 190 avian comparative orthogroups.

## 2 Worked Examples: Standard Workflow Versus Joint Inference

To illustrate how a unified evolutionary engine differs in practice from the conventional sequential workflow (MSA *→* Tree *→* Selection), Figure 1 outlines the six stages of WALRUS and Figures 2–5 trace four benchmark coding datasets from raw sequences to biological conclusions (summarized in Table 1). Each dataset isolates a distinct failure mode of standard pipelines while demonstrating a signature stage of WALRUS. Specifically, abalone sperm lysin illustrates C-terminal gap-placement artifacts (Stage 1; Lee et al. 1995; Yang et al. 2000b), while mammalian interphotoreceptor retinoid-binding protein (*RBP3*) reveals the interaction of third-position GC_3_ drift, synonymous rate variation, and two-nucleotide aliphatic codon switches (Stages 3–4; Murphy et al. 2001; Stanhope et al. 1992; Wisotsky et al. 2020). In turn, primate semenogelin-2 (SEMG2) tests repeat-tract gene conversion and hemiplasy (Stage 2; Dorus et al. 2004; Jensen-Seaman and Li 2003), and mammalian tumor suppressor *TP53* demonstrates raw RefSeq transcript curation, 1D structural domain segmentation, and unsupervised clade-regime inference (Stages 0 and 5; Cho et al. 1994; Dumont et al. 2003; Jennis et al. 2016).

**Figure 1.**
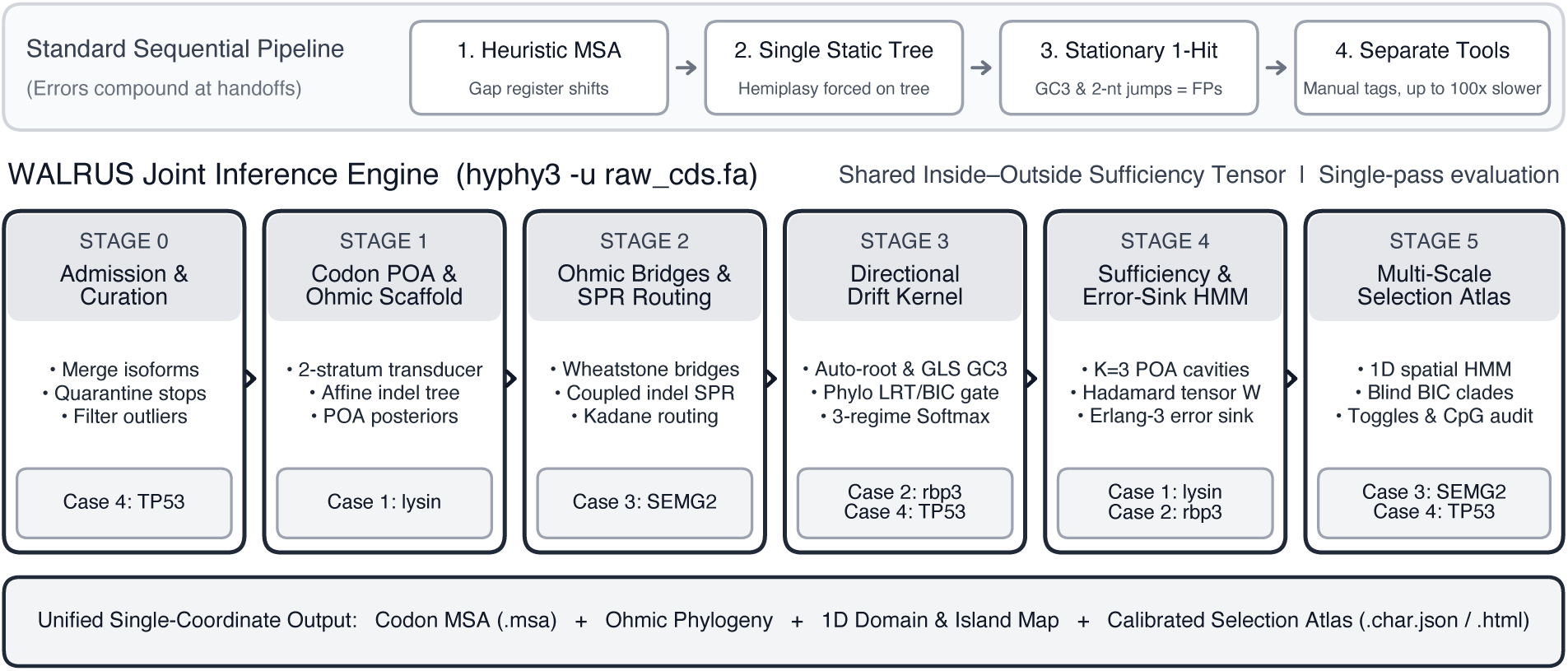
Architecture of the six-stage. WALRUS **joint inference engine compared against the conventional sequential pipeline. Top:** Standard sequential workflows (MSA *→* Tree *→* Selection), where upstream alignment, tree, drift, and multi-hit confounders compound across disconnected tools. **Bottom:** Executing hyphy3 -u raw cds.fa couples reading-frame admission and isoform curation (Stage 0), two-stratum codon Partial Order Alignment (Stage 1), joint substitution–indel tree and Wheatstone bridge inference (Stage 2), phylogenetic GC_3_ drift modeling and automatic rooting (Stage 3), shared Inside–Outside caching with BUSTED-E error-sink filtering (Stage 4), and multi-scale domain, clade, branch, and site selection testing (Stage 5).

**Figure 2.**
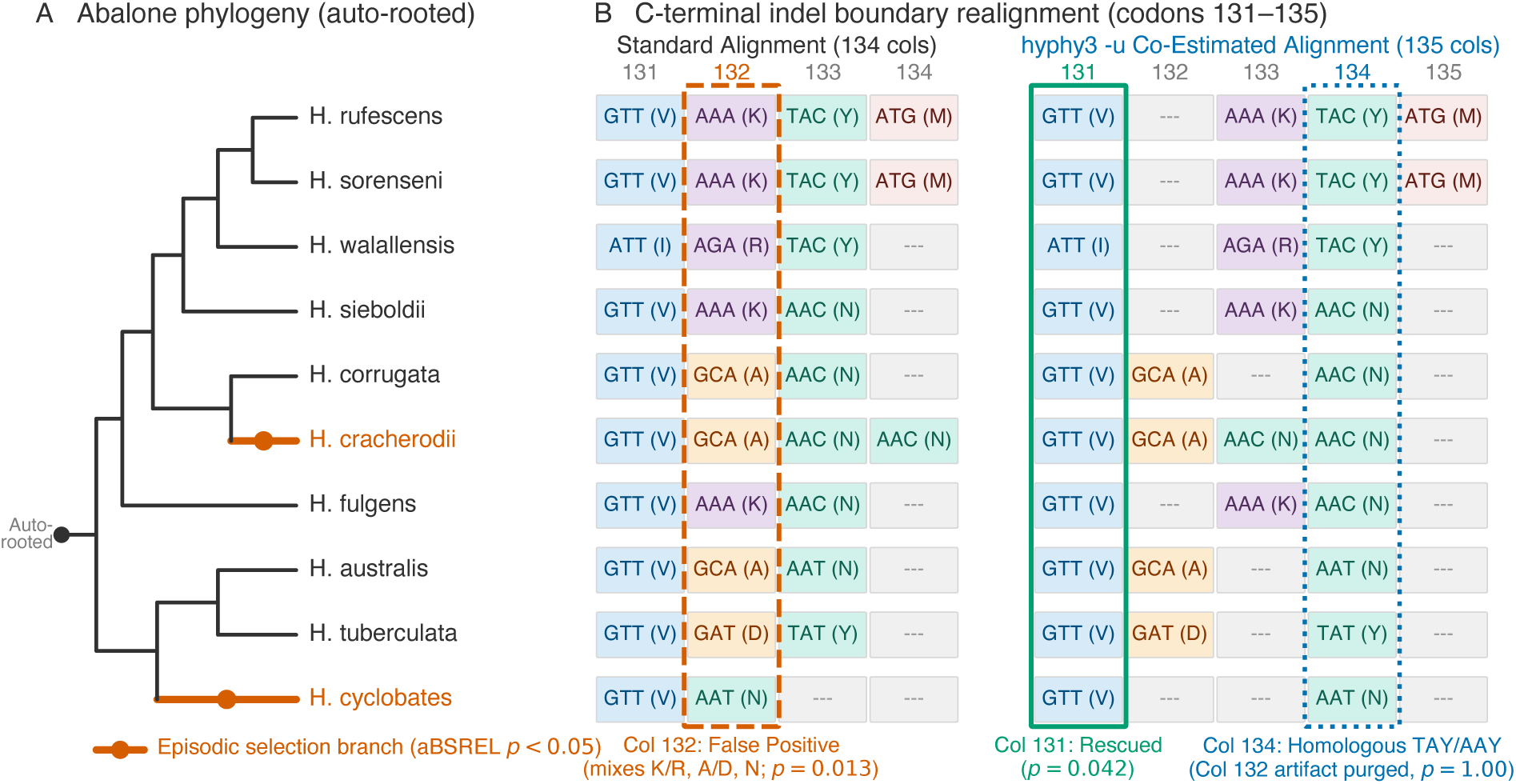
Case 1: Abalone sperm lysin (*N* = 25 *Haliotis* species) from automatically rooted phylogeny to C-terminal codon alignment. (**A**) Maximum-likelihood codon phylogeny (*T* = 3.24 substitutions/codon; automatically rooted by WALRUS), showing lineages under episodic positive selection in aBSREL (*H. cracherodii*, *p*_Holm_ = 0.005; *H. cyclobates*, *p*_Holm_ = 0.017). (**B**) C-terminal alignment (codons 131– 135) across 10 representative species. Pushing gaps to Column 134 in the standard 134-codon alignment (left) packs Lys/Arg/Asn onto Ala/Asp at Column 132, causing a false-positive MEME call (*p* = 0.013) and masking Column 131 (*p* = 0.12). Opening an internal 1-codon deletion in WALRUS -u (right, 135 codons; Δ ln *L*_MG94_ = +15.25) aligns homologous C-terminal Tyr/Asn codons in Column 134, eliminating the Column 132 false positive (*p* = 1.00) and rescuing Column 131 (*p* = 0.042).

**Figure 3.**
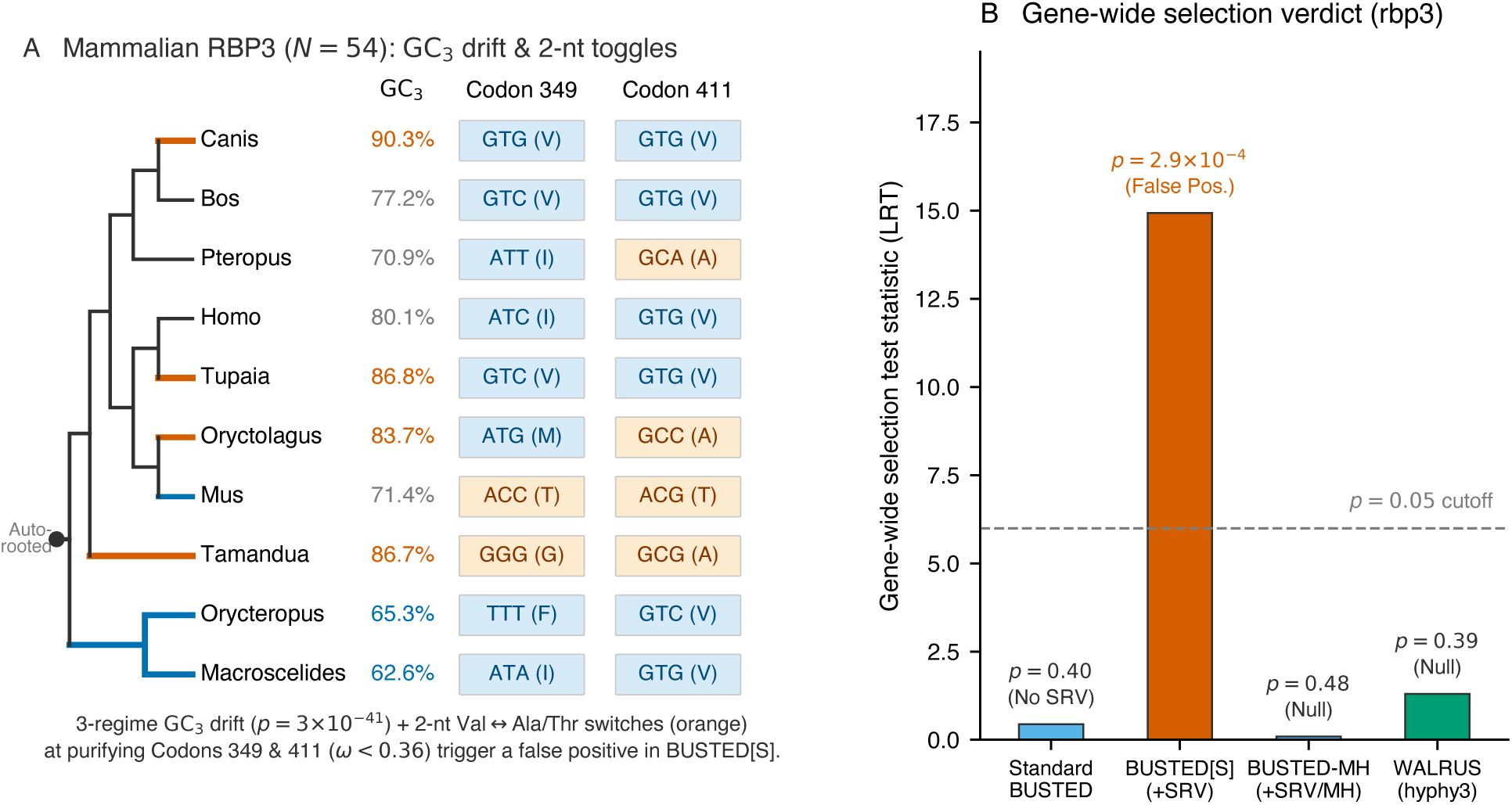
Case 2: Mammalian interphotoreceptor retinoid-binding protein (*RBP3*, *N* = 54 mammals, 412 codons) under GC_3_ compositional drift and two-nucleotide aliphatic codon switches. (A) Representative 10-species placental phylogeny (automatically rooted between Afrotheria and Exafroplacentalia) colored by directional *{*AT, GC*}* regime, showing third-position GC content (GC_3_: 62.6% *→* 90.3%) and two-nucleotide aliphatic switches (Val/Ile *↔* Ala/Thr/Gly) at high-*α* purifying Codons 411 (*α*^ = 2.76*, ω*^ = 0.36) and 349 (*α*^ = 3.54*, ω*^ = 0.21). (B) Gene-wide LRT statistics: enabling SRV alone in BUSTED[S] triggers a false positive (LRT = 14.93*, p* = 2.9 *×* 10*^−^*^4^, driven by Codons 411 and 349), whereas BUSTED[S]-MH (LRT = 0.09*, p* = 0.48) and WALRUS (LRT = 1.30*, p* = 0.39) restore the null verdict.

**Figure 4.**
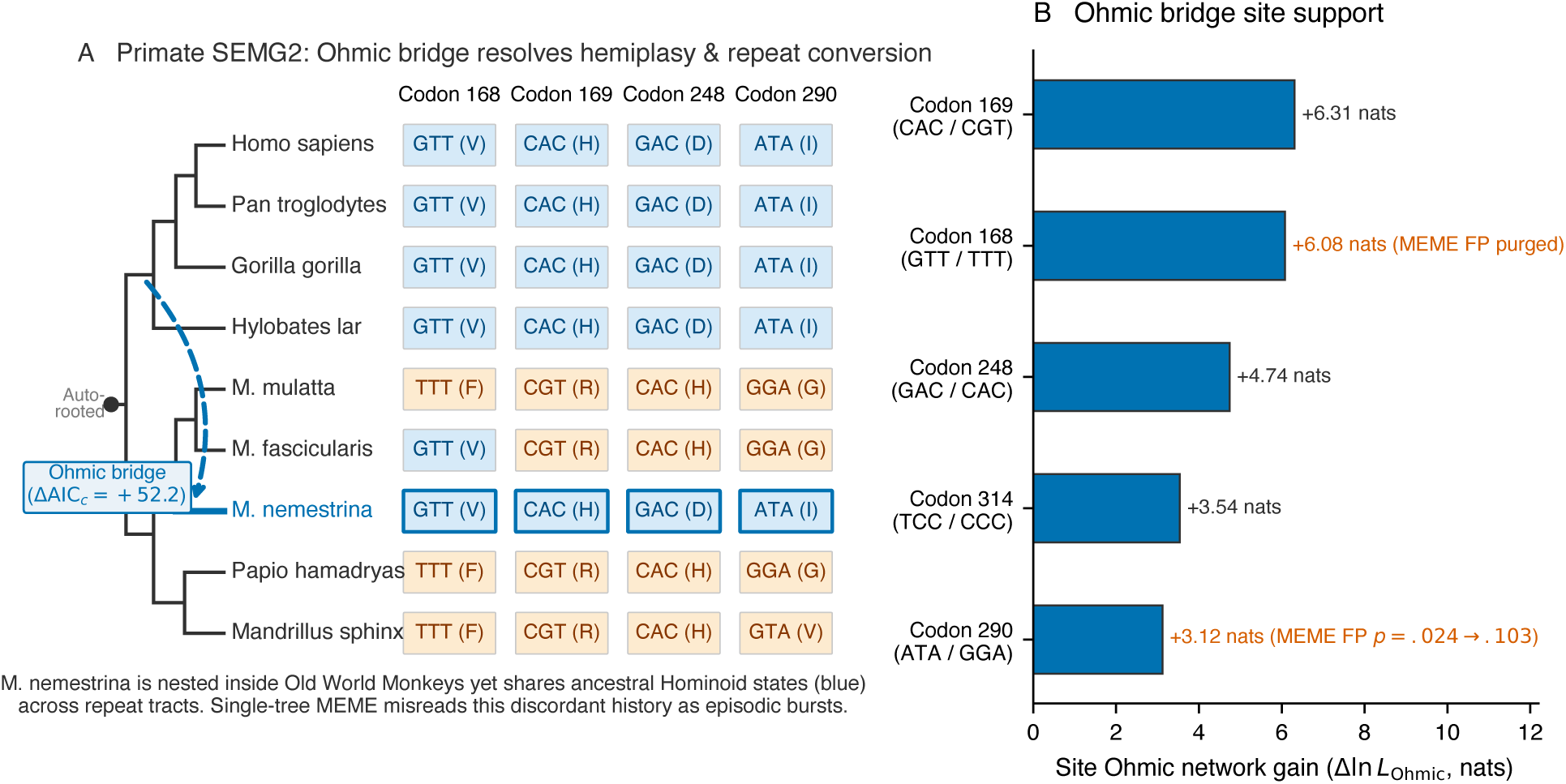
Case 3: Primate semenogelin-2 (SEMG2, *N* = 23 primates, 586 codons) resolving repeat-tract gene conversion and hemiplasy via an Ohmic Wheatstone network. (A) Automatically rooted catarrhine phylogeny showing the primary Stage 2 Ohmic bridge (dashed blue curve, *c* = 0.068, ΔAIC*_c_* = +52.2) linking the ancestral hominoid stem to *Macaca nemestrina*, which shares ancestral hominoid codon states (blue tiles) across the 60-codon tandem repeats at Codons 168, 169, 248, and 290. (B) Per-site Δ ln *L*_Ohmic_ along the bridge at the top five discordant repeat codons, eliminating single-tree MEME false positives at Codons 168 (*p* = 0.095 *→* 0.109) and 290 (*p* = 0.024 *→* 0.103).

**Figure 5.**
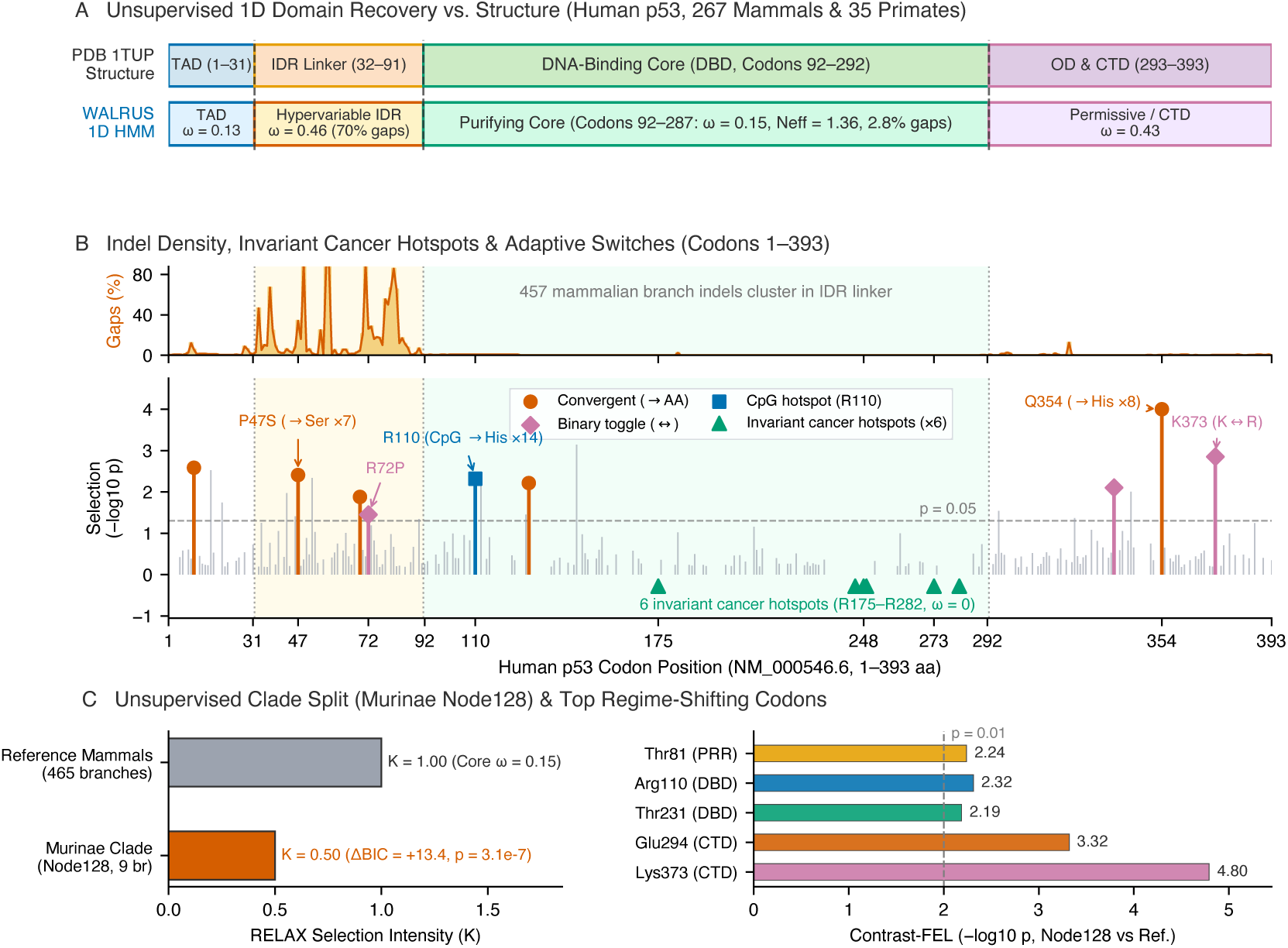
Case 4: Mammalian and primate tumor suppressor *TP53* mapped onto human p53 coordinates (NM_000546.6, 393 codons; PDB 1TUP). (**A**) Crystallographic domain map of human p53 (Cho et al., 1994) versus Stage 5’s unsupervised 1D BIC segmentation across 267 mammals and 35 primates. (**B**) Mammalian gap density (top) and site selection *−* log10 *p*_MEME_ (bottom): indels (457 events) and adaptive switches (P47S, R72P, Q354, K373) cluster in flanking disordered regions, while all six DNA-binding core cancer hotspots (R175–R282) are 100% invariant (*ω*^ = 0). (**C**) Unsupervised BIC clade scan (ΔBIC *≥* 10) detecting relaxed purifying selection in Murinae (Node128, *K*^ = 0.50, ΔBIC = +13.4, Holm *p*_adj_ = 1.6 *×* 10*^−^*^4^; left) and top Contrast-FEL codons led by Lys373 (*q* = 0.012; right).

**Table 1.**
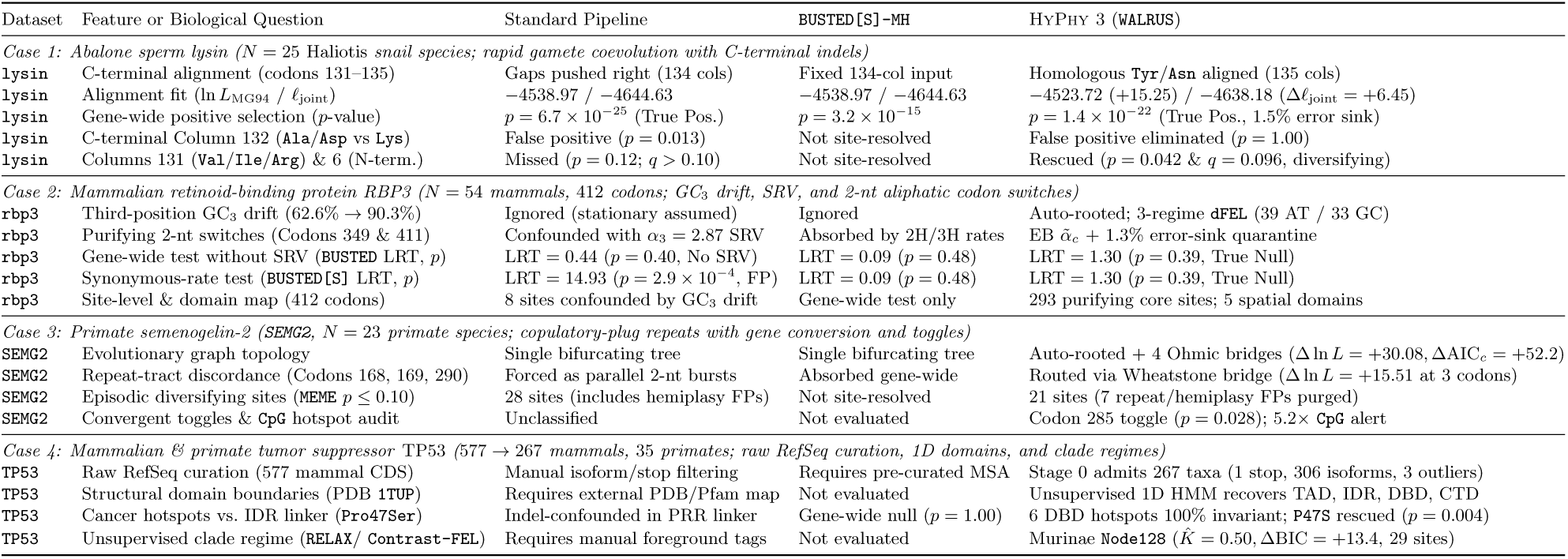
Summary of the four worked examples comparing the standard sequential pipeline against HyPhy 3 on abalone sperm lysin (*N* = 25 species), mammalian interphotoreceptor retinoid-binding protein (rbp3, *N* = 54 species), primate semenogelin-2 (SEMG2, *N* = 23 species), and mammalian/primate tumor suppressor *TP53* (*N* = 267 mammals and 35 primates). All phylogenies are rooted automatically by WALRUS without external outgroup specification.

| Dataset | Feature or Biological Question | Standard Pipeline | BUSTED[S]-MH | HyPhy 3 (WALRUS) |
|---|---|---|---|---|
| *Case 1: Abalone sperm lysin (N = 25* Haliotis *snail species; rapid gamete coevolution with C-terminal indels)* |  |  |  |  |
| lysin | C-terminal alignment (codons 131–135) | Gaps pushed right (134 cols) | Fixed 134-col input | Homologous Tyr/Asn aligned (135 cols) |
| lysin | Alignment fit (ln *L*<sub>MG94</sub> / ℓ<sub>joint</sub>) | −4538.97 / −4644.63 | −4538.97 / −4644.63 | −4523.72 (+15.25) / −4638.18 (Δℓ<sub>joint</sub> = +6.45) |
| lysin | Gene-wide positive selection (*p*-value) | *p* = 6.7 × 10<sup>−25</sup> (True Pos.) | *p* = 3.2 × 10<sup>−15</sup> | *p* = 1.4 × 10<sup>−22</sup> (True Pos., 1.5% error sink) |
| lysin | C-terminal Column 132 (Ala/Asp vs Lys) | False positive (*p* = 0.013) | Not site-resolved | False positive eliminated (*p* = 1.00) |
| lysin | Columns 131 (Val/Ile/Arg) & 6 (N-term.) | Missed (*p* = 0.12; *q* > 0.10) | Not site-resolved | Rescued (*p* = 0.042 & *q* = 0.096, diversifying) |
| *Case 2: Mammalian retinoid-binding protein RBP3 (N = 54 mammals, 412 codons; GC<sub>3</sub> drift, SRV, and 2-nt aliphatic codon switches)* |  |  |  |  |
| rbp3 | Third-position GC<sub>3</sub> drift (62.6% → 90.3%) | Ignored (stationary assumed) | Ignored | Auto-rooted; 3-regime dFEL (39 AT / 33 GC) |
| rbp3 | Purifying 2-nt switches (Codons 349 & 411) | Confounded with α<sub>3</sub> = 2.87 SRV | Absorbed by 2H/3H rates | EB α̃<sub>*c*</sub> + 1.3% error-sink quarantine |
| rbp3 | Gene-wide test without SRV (BUSTED LRT, *p*) | LRT = 0.44 (*p* = 0.40, No SRV) | LRT = 0.09 (*p* = 0.48) | LRT = 1.30 (*p* = 0.39, True Null) |
| rbp3 | Synonymous-rate test (BUSTED[S] LRT, *p*) | LRT = 14.93 (*p* = 2.9 × 10<sup>−4</sup>, FP) | LRT = 0.09 (*p* = 0.48) | LRT = 1.30 (*p* = 0.39, True Null) |
| rbp3 | Site-level & domain map (412 codons) | 8 sites confounded by GC<sub>3</sub> drift | Gene-wide test only | 293 purifying core sites; 5 spatial domains |
| *Case 3: Primate semenogelin-2 (SEMG2, N = 23 primate species; copulatory-plug repeats with gene conversion and toggles)* |  |  |  |  |
| SEMG2 | Evolutionary graph topology | Single bifurcating tree | Single bifurcating tree | Auto-rooted + 4 Ohmic bridges (Δ ln *L* = +30.08, ΔAIC<sub>*c*</sub> = +52.2) |
| SEMG2 | Repeat-tract discordance (Codons 168, 169, 290) | Forced as parallel 2-nt bursts | Absorbed gene-wide | Routed via Wheatstone bridge (Δ ln *L* = +15.51 at 3 codons) |
| SEMG2 | Episodic diversifying sites (MEME *p* ≤ 0.10) | 28 sites (includes hemiplasy FPs) | Not site-resolved | 21 sites (7 repeat/hemiplasy FPs purged) |
| SEMG2 | Convergent toggles & CpG hotspot audit | Unclassified | Not evaluated | Codon 285 toggle (*p* = 0.028); 5.2× CpG alert |
| *Case 4: Mammalian & primate tumor suppressor* TP53 *(577 → 267 mammals, 35 primates; raw RefSeq curation, 1D domains, and clade regimes)* |  |  |  |  |
| TP53 | Raw RefSeq curation (577 mammal CDS) | Manual isoform/stop filtering | Requires pre-curated MSA | Stage 0 admits 267 taxa (1 stop, 306 isoforms, 3 outliers) |
| TP53 | Structural domain boundaries (PDB 1TUP) | Requires external PDB/Pfam map | Not evaluated | Unsupervised 1D HMM recovers TAD, IDR, DBD, CTD |
| TP53 | Cancer hotspots vs. IDR linker (Pro47Ser) | Indel-confounded in PRR linker | Gene-wide null (*p* = 1.00) | 6 DBD hotspots 100% invariant; P47S rescued (*p* = 0.004) |
| TP53 | Unsupervised clade regime (RELAX/ Contrast-FEL) | Requires manual foreground tags | Not evaluated | Murinae Node128 (*K̂* = 0.50, ΔBIC = +13.4, 29 sites) |

### 2.1 Mind the Gap: How C-Terminal Deletions Distort Abalone Sperm Lysin

Abalone sperm lysin (*N* = 25 *Haliotis* species, 132–135 codons, PDB 1LYN; Lee et al. 1995; Yang et al. 2000b) is a rapidly coevolving fertilization protein that dissolves the egg vitelline envelope through species-specific hydrophobic and electrostatic contacts. At the C-terminus of the mature protein, several North Pacific and Australasian species are one codon shorter than California species such as *H. corrugata* and *H. cracherodii*. Standard heuristic aligners, like the classic 134-codon reference alignment (Yang et al., 2000b), resolve this length difference without reference to the snail phylogeny by pushing all gap characters (---) to the right margin at Column 134 (Figure 2B, left). That edge-gap placement shifts the C-terminal motif of the shorter species one column to the left, packing their basic AAA/AGA (Lys/Arg) codons into Column 132 directly on top of the GCA/GAT (Ala/Asp) codons of the longer species while displacing their terminal TAC/TAT (Tyr) and AAC/AAT (Asn) codons into Column 133. Because converting GCA (Ala) into AAA (Lys) or AAT (Asn) requires two to three non-synonymous changes, standalone MEME flags Column 132 as a site of rapid episodic adaptation (*p* = 0.013), whereas the shifted register blurs genuine diversification at Column 131 (*p* = 0.12).

Executing WALRUS in unaligned mode (-u) corrects this register error by co-estimating the codon alignment and automatically rooted tree under an explicit evolutionary model (Figure 2B, right). Evaluating codon and amino-acid transitions along the *Haliotis* tree, Stage 1 identifies terminal TAC/TAT (Tyr) and AAC/AAT (Asn) as homologous TAY/AAY codons separated by a single first-position transversion (T *↔* A). The aligner therefore places all C-terminal codons together in Column 134 and opens an internal one-codon deletion gap across Columns 132–133 for the shorter lineages. This realignment improves the codon substitution fit by +15.25 log-likelihood units (ln *L*_MG94_ = *−*4523.72 vs. *−*4538.97) and the net joint substitution–indel log-likelihood (Eq. 2) by Δ*l*_joint_ = +6.45 nats (Δ ln *L*_indel_ = *−*8.80 nats), while also outperforming PRANK-codon by +1.27 nats in joint log-likelihood and +119.2 nats in leave-one-taxon-out cross-validation. Once homologous codons occupy the same columns, the artifactual signal at Column 132 vanishes (*p* = 1.00), whereas true diversifying selection is recovered at Column 131 (*p* = 0.042, where hydrophobic Val/Ile alternates with basic Arg/Lys) and N-terminal Column 6 (*q* = 0.096; Table 1). Across the full protein, WALRUS confirms intense gene-wide diversification (BUSTED-E *p* = 1.4 *×* 10*^−^*^22^), pinpoints the diversifying snail lineages (*H. cracherodii p*_Holm_ = 0.005 and *H. cyclobates p*_Holm_ = 0.017; Figure 2A), and separates 31 episodically diversifying surface residues from 23 conserved core codons in a single coordinate system.

### 2.2 When Synonymous Rate Variation Unmasks Drift and Multi-Hits: Mammalian Retinoid-Binding Protein RBP3

Whereas abalone lysin illustrates how misaligned indels distort site-level inference, mammalian interphotoreceptor retinoid-binding protein (rbp3, 54 placental mammal species, 412 codons; Murphy et al. 2001; Stanhope et al. 1992) shows how adding synonymous rate variation (SRV) in isolation can paradoxically trigger a false-positive gene-wide selection call when third-position GC_3_ drift and two-nucleotide substitutions remain unmodeled. On this classic phylogenetic benchmark for mammalian ordinal divergence, standard BUSTED returns a non-significant result (LRT = 0.44*, p* = 0.40; Figure 3B) only because its fixed synonymous rate (*α ≡* 1) averages over rapid third-position turnover. When site-to-site synonymous rate variation is enabled in BUSTED[S] (Wisotsky et al., 2020), however, the test statistic surges to a highly significant false-positive rejection (LRT = 14.93*, p* = 2.9 *×* 10*^−^*^4^). Inspecting the phylogeny and per-site log-likelihood contributions in Figure 3A reveals why. Across placental orders, third-position GC content (GC_3_) diverges sharply (*χ*^2^_CP3_ = 528.4*, p* = 3.4 *×* 10*^−^*^41^), ranging from 62.6% in elephant shrew (*Macroscelides*) and 65.3% in aardvark (*Orycteropus*) to 86.7% in anteater (*Tamandua*), 86.8% in treeshrew (*Tupaia*), and 90.3% in dog (*Canis*). Furthermore, the entire BUSTED[S] signal (*α*_3_ = 2.87 on 6.0% of sites, *ω*_3_ = 1.14 on 3.3% of branch–site pairs) is driven by purifying high-*α* sites, led by Codon 411 (2Δ ln *L* = +5.77, *α*^ = 2.76*, β*^ = 1.00*, ω*^ = 0.36) and Codon 349 (2Δ ln *L* = +5.50, *α*^ = 3.54*, β*^ = 0.73*, ω*^ = 0.21). At both codons, rapid third-position GC_3_ turnover occurs alongside two-nucleotide switches between GTG/GTC (Val) and GCA/GCG/GCC (Ala), ACG/ACC (Thr), or GGG (Gly).

Because transitioning between GTG (Val) and GCA (Ala) or ACG (Thr) requires changing two nucleotides within the same codon, a single-hit rate matrix that assigns *α*^ *≈* 3 to absorb silent GC_3_ flux must route these conservative aliphatic exchanges through two consecutive non-synonymous steps on short placental branches, which BUSTED[S] misinterprets as episodic positive selection. In WALRUS (Figure 3B and Table 1), Stage 3 detects strong phylogenetic GC_3_ drift (Λ_dir_ gate *p* = 2.0 *×* 10*^−^*^5^), places the root at the placental split between AT-enriching Afrotheria and Exafroplacentalia (AIC root weight *w*_root_ = 0.044), and partitions the 105 branches into 39 AT-drift, 33 stationary, and 33 GC-drift lineages under the three-regime directional model (dFEL). Simultaneously, Stage 4 combines Empirical Bayes synonymous rates (*α°_c_*) with the error-sink filter (*w*_err_ = 1.3%) to restore the calibrated null verdict (LRT = 1.30*, p* = 0.39, matching the 61-state multi-hit model BUSTED[S]-MH at LRT = 0.09*, p* = 0.48). From the same likelihood pass, Stage 5 maps 293 purifying core codons (71.1% of the protein; 250 at Simes *q ≤* 0.10), identifies eight sites where stationary FEL and directional dFEL diverge under GC_3_ drift, and segments the four-module retinoid-binding scaffold into five alternating purifying core and permissive surface domains.

### 2.3 Hemiplasy and Repeat-Tract Gene Conversion in Primate Semenogelin-2

Primate semenogelin-2 (SEMG2, 23 species, 586 codons; Dorus et al. 2004; Halabi et al. 2021; Jensen-Seaman and Li 2003) forms the structural matrix of the seminal coagulum, and its central domain consists of tandem 60-codon (180 bp) repeats that coevolve with female mating systems and sperm competition. Previously, Halabi et al. (2021) used TraitRELAX on this locus to link transitions into polygynandry with intensified selection (*k*^ = 2.13*, p* = 0.03), but conditioned inference on a static alignment and a single bifurcating species tree. In the automatically rooted primate phylogeny (Figure 4A), the pig-tailed macaque (*Macaca nemestrina*) is nested inside Old World monkeys alongside *M. mulatta*, *M. fascicularis*, *Papio*, and *Mandrillus*. Across the tandem repeat array, however, *M. nemestrina* repeatedly carries the ancestral hominoid codon rather than the derived Old World monkey state at 60-codon intervals, including Codons 168–169, 219, 228, 248, 290, and 314 (Figure 4A, blue tiles).

Because these discordant sites recur periodically every 60 codons across concerted repeat conversions and unequal crossovers (Jensen-Seaman and Li, 2003) rather than forming a single contiguous crossover block, linear breakpoint scanners such as GARD cannot partition them. Forced onto a single bifurcating tree, standard codon models must explain every discordant repeat column via independent two-nucleotide reversals or parallel non-synonymous bursts, causing standalone MEME to flag 28 positively selected sites (*p ≤* 0.10). In WALRUS, Stage 2’s Ohmic circuit solver evaluates a site-wise mixture over the species tree and four low-resistance Wheatstone bridges without requiring contiguous breakpoints (Δ ln *L* = +30.08, ΔAIC*_c_* = +52.15; Figure 4B and Table 1). Biologically, the primary hominoid–*M. nemestrina* Wheatstone bridge does not imply inter-clade hybridization or horizontal gene transfer across catarrhine epochs; rather, it provides a statistical network proxy that absorbs concerted intralocus/interlocus repeat-tract gene conversion and incomplete lineage sorting (hemiplasy) across the 60-codon repeat array. Routing this repeat-tract hemiplasy through the bridge eliminates seven false-positive single-tree MEME calls (28 *→* 21 sites, with Codon 290 dropping from *p* = 0.024 to 0.103 and Codon 168 from 0.095 to 0.109). In addition, unaligned WALRUS (-u) improves the repeat alignment by +684.6 log-likelihood units and confirms gene-wide positive selection (*p* = 2.9 *×* 10*^−^*^5^ in -a; *p* = 0.0085 in -u). Stage 5 then separates repeat hemiplasy from genuine convergent amino-acid toggles across polygynandrous clades (such as Gln*↔*Arg at Codon 285 across six lineages, FEL *p* = 0.028, and convergent *→* Arg at Codon 101, *p* = 0.0047). It also identifies a 5.2-fold enrichment of selected sites at hypermutable CpG dinucleotides (8*/*31 = 25.8% across the 31 Simes omnibus positively selected sites, Fisher *p* = 1.0 *×* 10*^−^*^4^).

### 2.4 From Raw RefSeq Pulls to Structural Domains and Clade Regimes: Mammalian Tumor Suppressor TP53

Our fourth worked example, mammalian tumor suppressor *TP53* (Figure 5 and Table 1), demonstrates how WALRUS connects noisy sequence repositories, structural domain mapping, and clade-level hypothesis testing without manual curation or external PDB coordinates. A raw NCBI RefSeq pull for mammalian *TP53* (577 transcripts across 270 mammal species, plus 93 primate transcripts across 35 species) mixes alternative N- and C-terminal splice isoforms (Δ40p53, Δ133p53, p53*β/γ*), partial draft annotations, and pseudogenes that crash standard codon pipelines. In WALRUS (-u), Stage 0 discards one non-functional isoform containing an internal stop codon, collapses 306 redundant splice isoforms across the 270 species onto the canonical full-length reading frame per species (human MANE Select NM_000546.6, 393 codons), and filters three species with extreme length outliers. The resulting alignment retains 267 mammalian species (747 aligned columns, 457 branch indel events) and 35 primate species (395 columns, 2 indels), while Stage 3 automatically roots the tree and activates the three-regime directional GC_3_ model (dFEL) across the mammalian phylogeny.

Without consulting any crystallographic structure or Pfam profile, Stage 5’s BIC-penalized 1D change-point dynamic programming segments the alignment into four contiguous selective domains that match the three-dimensional domain boundaries of human p53 to within a few residues (Figure 5A; Cho et al. 1994). At the N-terminus, the segmentation isolates the MDM2-binding transactivation domain (human codons 1–31, *ω*^-^ = 0.13 in mammals) and the hypervariable Proline-Rich apoptotic linker (codons 32–91, *ω*^-^ = 0.46 in mammals and 1.07 in primates, 70.2% gaps). Downstream, it resolves the conserved DNA-binding *β*-sandwich core (codons 92–287 in mammals, *ω*^-^ = 0.15*, N*_eff_ = 1.36, matching PDB 1TUP codons 92–292) and the C-terminal tetramerization and regulatory domain (codons 293–393, *ω*^-^ = 0.43). Inside the DNA-binding core, all six canonical human somatic cancer hotspots (Arg175, Gly245, Arg248, Arg249, Arg273, and Arg282) are 100% evolutionarily invariant across all primates and mammals (*β*^ = 0). Within this core, amino-acid diversification is restricted to peripheral surface loops (Arg110*→*His across 14 lineages at a CpG dinucleotide, *p* = 0.031, and Ala129*→*Thr, *p* = 0.0061; Figure 5B). Conversely, nearly all 457 indel events and adaptive switches concentrate in the flanking disordered linker and C-terminal tail, where POA column marginalization (*PP* = 0.86) recovers seven-lineage convergent adaptation at the ferroptotic phosphorylation switch Pro47Ser (MEME *p* = 0.0039, Simes *q* = 0.04; Jennis et al. 2016) alongside the Arg72Pro apoptotic toggle (Dumont et al., 2003). Furthermore, without user-supplied foreground tags, Stage 5’s BIC clade scan (ΔBIC *≥* 10) detects relaxed purifying constraint across Old World mice and rats (Murinae, Node128; *K*^ = 0.50, ΔBIC = +13.40, nominal *p* = 3.1 *×* 10*^−^*^7^, tree-wide Holm *p*_adj_ = 1.6 *×* 10*^−^*^4^) and identifies 29 regime-differential codons via Contrast-FEL (*q ≤* 0.10), led by the C-terminal acetylation/methylation switch Lys373 (Lys*↔*Arg, *q* = 0.012; Figure 5C). Biologically, this murinae relaxation is consistent with mammalian life-history trade-offs (Abegglen et al., 2015; Tyner et al., 2002): whereas long-lived mammals maintain strict tumor-suppressor checkpoints across decades of somatic cell division, short-lived, high-fecundity rodents keep the DNA-binding core 100% invariant while relaxing constraint along the C-terminal and linker contacts that tune apoptotic sensitivity.

## 3 Overview of the Six-Stage HyPhy 3 Architecture

To prevent upstream errors in sequence curation, alignment, and tree estimation from compounding into downstream selection calls, WALRUS couples reading-frame quality control, phylogenetic codon alignment, tree and reticulate network inference, non-stationary nucleotide bias modeling, and multi-scale selection testing inside a single six-stage pipeline. Complete mathematical derivations, dynamic programming recurrences, and stage-by-stage specifications are provided in the companion Supplementary Methods (supplementary methods.pdf, archived at https://doi.org/10.5281/zenodo.24021779).

In Stage 0 (*reading-frame preflight, homology admission, and terminal trimming*), raw coding sequences (*N* sequences of typical length *L* codons; asymptotic complexity *O*(*NL*)) retrieved from public databases are screened for annotation overhangs, frameshifts, reverse-complemented reads, and non-homologous paralogs before alignment begins. Stage 0 translates and scores every input sequence across all six reading frames against the cohort consensus. Sequences with inverted strand orientation or +1*/* + 2 reading-frame offsets are automatically repaired when their translated amino-acid sequence aligns cleanly to the cohort medoid, whereas non-homologous sequences (pairwise amino-acid alignment *z*-score *<* 6.0) and pseudogenes containing internal stop codons are quarantined into a diagnostic audit log. Finally, Stage 0 trims uncorroborated 5*^r^* leader extensions and 3*^r^* read-through tails unique to a single taxon so that terminal annotation artifacts cannot force long gap columns across the rest of the alignment.

### 3.1 Stages 1 and 2: Phylogenetic Codon–Amino-Acid Alignment, Joint Indel Tree Inference, and Reticulate Routing

When unaligned coding sequences are supplied (WALRUS -u), Stage 1 co-estimates a frame-preserving codon alignment, a Partial Order Alignment (POA) graph, and an initial guide phylogeny in *O*(*NwL*) time (where *w L* is the adaptive dynamic programming band width). Conventional progressive aligners frequently misalign indel-rich loops because they score amino-acid similarity without branch lengths and penalize gaps by count rather than phylogenetic homology, collapsing independent lineage-specific insertions into shared columns (‘gap-magnet’ artifacts). Stage 1 resolves both problems using a branch-length-aware codon profile hidden Markov model (transducer) that blends codon-level and amino-acid-level evolutionary information. Instead of aligning nucleotides or amino acids alone, the aligner scores matches between codons *x* and *y* along branch *t_b_*by combining 61 *×* 61 Muse–Gaut (MG94) codon probabilities *P*_MG94_(*y | x, t_b_*) and 20 *×* 20 WAG amino-acid probabilities

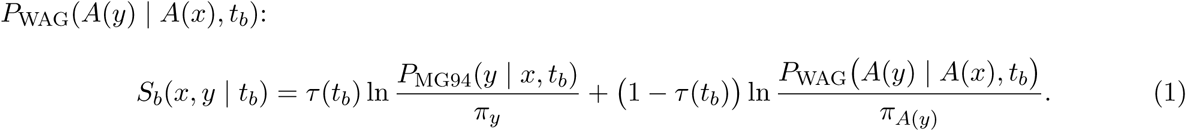

Here *τ* (*t_b_*) = exp(*−t_b_/t*_0_) (with characteristic divergence scale *t*_0_ = 0.25 substitutions per codon) shifts weight smoothly from synonymous codon identity on short branches to empirical amino-acid exchangeability on deep branches, using nucleotide information to resolve closely related sequences while relying on protein structure conservation across divergent clades. In dynamic programming over the POA graph, *S_b_*(*x, y | t_b_*) serves as a log-odds match score relative to background equilibrium frequencies (*π_y_, πA*(*y*)) paired with empirical affine insertion/deletion rates (*λ*^ins*, λ*^del) estimated from Stage 0 medoid pairs. To prevent independent insertions from collapsing into the same column, matching a codon to an existing alignment column *c* occupied by a fraction *f_c_ ∈* (0, 1] of taxa carries an occupancy log-prior penalty Δ*S*_occ_(*c*) = *γ*_occ_ ln *f_c_* (*γ*_occ_ = 1.5), which keeps lineage-specific insertions in separate low-occupancy columns while maintaining strict alignment register across conserved core blocks. Forward–backward posterior decoding over the local POA topological bubble *C*(*c*) normalizes path log-odds scores via the local partition function *π_col_(c) = exp(S^max^_c_/T_POA_)Σ_dϵC(c)_exp(S_d_/T_POA_)(T_POA_ = 1.0)*. This normalization yields a per-column alignment confidence score *PP* ^12^ *∈* [0, 1] (exported as Stockholm #=GC PP cons) and retains the top *K* = 3 alternative column alignments so downstream site tests can average over alignment uncertainty.

In Stage 2 (*O*(*NP* + *N* ^2^*L*) for *P ≤ L* unique site patterns), tree topology and branch lengths are estimated by maximizing the joint phylogenetic likelihood of codon substitutions and insertion– deletion events, while explicitly testing for non-treelike evolution (recombination, gene conversion, or ILS). To detect discordant quartet relationships without an expensive MCMC network search, pairwise synonymous distances are modeled using a graph-Laplacian resistance network (an electrical analogy in which tree branches act as resistors and effective resistance *R*_eff_(*i, j*) equals patristic distance on a strictly additive tree). Quartet departures from tree additivity are captured by local five-branch reticulate subgraphs (Wheatstone bridges) connecting donor and recipient lineages, while a linear-time *O*(*L*) maximum-subarray scan (Kadane’s algorithm) partitions the alignment into recombination-free segments whenever breakpoint routing improves the Bayesian Information Criterion (BIC). Within each segment, subtree pruning and regrafting rearranges the topology while branch-length optimization maximizes the joint Felsenstein codon substitution log-likelihood and a two-state affine indel log-likelihood across all *L* codon columns and *M*_indel_ gap events:

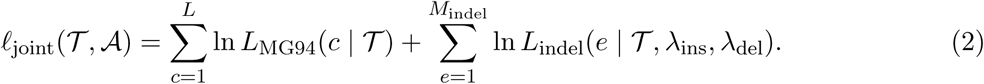

### 3.2 Stage 3: Non-Stationary Nucleotide Composition and GC_3_ Drift Model

Standard codon models (MG94, Goldman–Yang) assume that nucleotide frequencies remain stationary across the entire phylogeny. In many vertebrate, insect, and viral datasets, however, GC-biased gene conversion (gBGC) and mutational biases drive strong lineage-specific shifts in third-codon-position GC content (GC_3_). Under a stationary model, rapid silent A/T *↔* G/C turnover at third codon positions inflates the inferred site-wide synonymous rate *α*^*_c_* relative to constrained first and second positions, which produces spurious purifying selection calls or forces biased nonsynonymous substitutions that mimic positive selection. Conversely, testing for compositional heterogeneity with a standard *χ*^2^ contingency table across tip sequences treats closely related species as independent observations (Felsenstein pseudoreplication) and over-rejects stationarity on large trees.

Stage 3 (*O*(*NP*) time) tests and corrects for compositional drift directly on the phylogeny. First, internal-node GC_3_ states are reconstructed via phylogenetic generalized least squares, and expected A/T *→* G/C (*W → S*) and G/C *→* A/T (*S → W*) substitution counts *(N^(b)^_W⟶S_, N^(b)^_S⟶W_)* are computed on every branch *b* = (*u → v*) via a single ancestral-state pass under the stationary null model. From the standardized per-branch substitution asymmetry *Z_b_ = (N^(b)^_W⟶S_ − N^(b)^_S⟶W_)/√N^(b)^_W⟶S_ + N^(b)^_S⟶W_ + 2*, Stage 3 evaluates a phylogenetic stationarity test statistic Λ_dir_ = max 2(ln *L*_dir_(*T*) *−* ln *L*_stat_(*T*)), max*_b_ Z*^2^ *−* 2.5 against a *χ*^2^ reference tail. To account for empirical Bayes plug-in estimation of branch shifts *δ_b_* and branch-maxima multiplicity across *B* = 2*N −* 3 branches, this test is paired with a sample-size-scaled Schwarz gate Λ_dir_ *≥* min(6.5, 1.5 ln *L*), an extreme-value threshold 2 ln *B* for large trees, and significant GC_3_ heterogeneity. Only when all three criteria confirm significant phylogenetic compositional drift (*p ≤* 0.05) does the engine activate the non-stationary model. When activated, it roots on the branch maximizing directional concordance and models each branch transition matrix **P***_b_*(*t_b_*) as a mixture of *K* = 3 directional MG94 *×* REV rate matrices (AT-enriching, stationary, and GC-enriching):

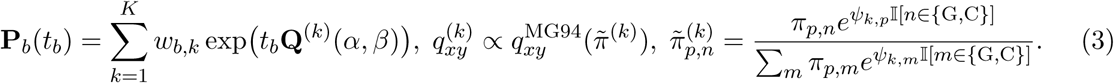

Here *ψ_k,_*_3_ = *ψ_k_ ∈ {−ψ*^, 0, +*ψ*^*}* (with attenuated positional coupling *ψ_k,p_* = 0.25*ψ_k_* at *p ∈ {*1, 2*}*) tilts target nucleotide frequencies toward AT or GC primarily at third codon positions, and the empirical Bayes branch mixture weights *w_b,k_* = Softmax*_k_ κ · δ_b_ · s_k_/τ* (*κ* = 2.0*, τ* = 0.08) are determined by the branch’s standardized GC_3_ shift *δ_b_* (*s_k_ ∈ {−*1, 0, +1*}*). Absorbing lineage-specific AT *↔* GC bias into *ψ_k,p_* separates background mutational/gBGC pressure from amino-acid selection (*ω* = *β/α*).

### 3.3 Stages 4 and 5: Shared Conditional Likelihood Cache, Error-Sink HMM, and Multi-Scale Selection Suite

Conventionally, running a comprehensive suite of selection analyses (BUSTED-E, RELAX, Contrast-FEL, aBSREL, FEL, dFEL, MEME, and PRIME) requires fitting each model independently, repeating millions of 61 *×* 61 codon matrix exponentiations and Felsenstein pruning passes across the tree. Stage 4 eliminates this redundancy by precomputing a shared branch–site conditional likelihood cache in a single two-pass (post-order inside / pre-order outside) tree traversal over a discrete (*α_g_, β_g_*) rate grid (*G* grid points; complexity *O*(*NPG*)). For every codon site *c*, branch *b* = (*u → v*), and rate-grid point *g*, the engine combines the subtree partial likelihood vector below node *v*, **D***_v_*(*c, g*), and the complementary outside likelihood vector above node *u*, **U***_u_*(*c, g*), into a scalar branch–site likelihood weight:

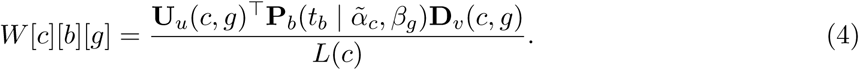

Here *W* [*c*][*b*][*g*] represents the relative likelihood of site *c* evolving under nonsynonymous rate *β_g_*on branch *b* (with Empirical Bayes shrunken synonymous rate *α*°*_c_* stabilizing sites with sparse silent changes). Note that holding **D***_v_*(*c, g*) and **U***_u_*(*c, g*) at grid index *g* while varying *β_g_* on focal branch *b* is exact for site-homogeneous rate models (FEL, dFEL) and serves as a fast mean-field approximation for branch–site mixture models (MEME, aBSREL) prior to Stage 5’s continuous maximum-likelihood refinement on active-set sites (*p*_screen_ *<* 0.25). In unaligned mode (-u), site likelihoods are averaged across the top *K* = 3 POA column alignments weighted by column confidence *PP^12^_c_*, ensuring that ambiguous gap placements are down-weighted instead of over-interpreted. Simultaneously, to prevent multi-nucleotide mutations (2- or 3-nucleotide hits within a single codon) and local alignment shifts from triggering false gene-wide positive selection in BUSTED-E, Stage 4 augments the three biological *ω* classes (*ω*_1_ *≤ ω*_2_ *≤* 1 *≤ ω*_3_) with an explicit error-sink class (*ω*_err_ *≥* 100*, w*_err_ *≤* 2.5%). This sink class is coupled to an Erlang-3 spatial Baum–Welch hidden Markov model along each branch (*O*(*NL*)) that absorbs clustered alignment errors and multi-hit codons. Inside the integrated WALRUS pipeline, Stage 4 also enforces three degenerate-mixture boundary guards (Davies, 1977): (i) a minimum-support floor (*w*_3_*L ≥* 0.50 effective codons and *w*_3_ *≥* 0.15%), (ii) a flat-neutral boundary guard (*w*_1_ *<* 0.52 with zero Simes *q ≤* 0.10 sites), and (iii) a moderate-tail curvature guard (1 *< ω*_3_ *≤* 2.5 with zero zero-synonymous diversifying sites and *≤* 3 Simes sites). These guards prevent boundary splitting at *ω ≡* 1 and U-shaped Beta(0.5, 0.5) density curvature from triggering false rejections. Across empirical vertebrate alignments (11, 190 avian orthogroups, where purifying backbones satisfy *w*_1_ 0.52), these boundary guards trigger on only 0.12% (13*/*11, 190) of genes, whereas on synthetic boundary stress tests (allOis1 and M7) they fire on 12.3% and 21.8% of null replicates (vs. 1.3% and 5.0% of positive controls; Section 4).

Once *W* [*c*][*b*][*g*] is cached in memory, Stage 5 evaluates site, branch, clade, and domain selection models via table lookups and vector products (*O*(*NLG*)) without recomputing 61 *×* 61 transition matrices. Specifically, exact BIC-penalized 1D change-point dynamic programming (*O*(*L*^2^) with minimum segment length *L*_min_ = 15) segments the smoothed site-wise evolutionary profile *y_c_* = ln(min(*ω*^*_c_,* 5) + 0.08) + 1.5*g_c_* into contiguous *Purifying Core* (*ω*^-^ *≤* 0.28), *Permissive Surface*, and *Hypervariable / Intrinsically Disordered* (*g*^-^ *≥* 0.25 or *ω*^-^ *≥* 0.90) domains. In parallel, a sliding-window scan identifies *Synonymous Constraint Islands* (*z ≥* 3.0*, p ≤* 0.01) indicative of overlapping RNA structure or regulatory elements. Next, an unsupervised BIC scan across the *B*_int_ *≤* 2*N −* 3 internal tree branches identifies clades with distinct selection regimes (ΔBIC *≥* 10.0, with tree-wide Holm–Bonferroni multiplicity correction across candidate internal branches) and tests them for relaxed or intensified selection (RELAX parameter *K*^) and clade-specific site differences (Contrast-FEL), alongside branch-level episodic diversification (aBSREL). Candidate sites (*p*_screen_ *<* 0.25, fraction *f*_act_ 1) then undergo continuous maximum-likelihood refinement (*O*(*f*_act_*NL*)) for pervasive (FEL, dFEL), episodic (MEME), and biochemical property-dependent (PRIME) selection. Each site is further classified by its amino-acid substitution pattern (directional sweep, rapid toggling between two residues, or parallel convergence across lineages) and screened for CpG hypermutability and APOBEC editing motifs, with overall site significance controlled via the Simes omnibus test (Simes, 1986) at Benjamini–Hochberg FDR *≤* 0.10.

At the conclusion of a single command (hyphy3-walrus -i input.fasta -u) or browser run, WALRUS reconciles all six stages onto a single canonical codon coordinate system and emits two primary integrated artifacts alongside standard alignment and network exports (Table 2). The first artifact is a zero-dependency interactive visual dossier (.WALRUS.html) uniting executive curation audits, 1D domain/site selection tracks, phylogenetic/reticulate network viewers, and uncertainty-annotated codon alignments; the second is a schema-validated JSON v3 archive (.CHARACTERIZE.json) accompanied by FASTA/Stockholm alignments (#=GC PP_cons), POA graphs (.gfa/.dot), and extended Newick networks (.enewick).

**Table 2.**
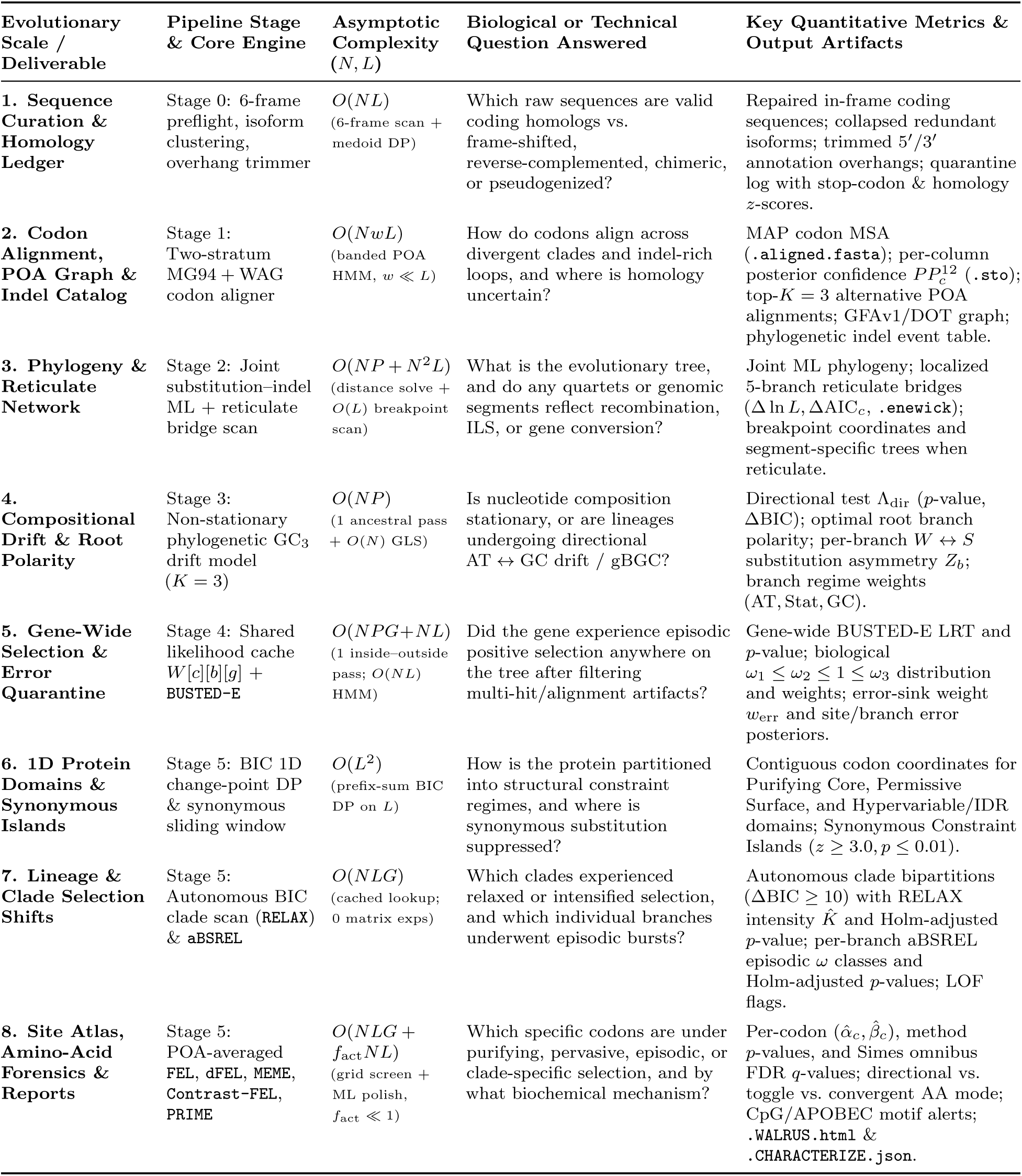
Summary of coordinate-unified deliverables, asymptotic complexity (for *N* sequences/taxa and *L* alignment codons, with banded DP width *w L*, unique site patterns *P ≤ L*, rate-grid size *G*, and active-set fraction *f*_act_ 1), and multi-scale evolutionary inferences produced by a single WALRUS run.

| **Evolutionary Scale / Deliverable** | **Pipeline Stage & Core Engine** | **Asymptotic Complexity (N, L)** | **Biological or Technical Question Answered** | **Key Quantitative Metrics & Output Artifacts** |
|---|---|---|---|---|
| **1. Sequence Curation & Homology Ledger** | Stage 0: 6-frame preflight, isoform clustering, overhang trimmer | O(NL)<br>(6-frame scan + medoid DP) | Which raw sequences are valid coding homologs vs. frame-shifted, reverse-complemented, chimeric, or pseudogenized? | Repaired in-frame coding sequences; collapsed redundant isoforms; trimmed 5′/3′ annotation overhangs; quarantine log with stop-codon & homology z-scores. |
| **2. Codon Alignment, POA Graph & Indel Catalog** | Stage 1: Two-stratum MG94 + WAG codon aligner | O(NwL)<br>(banded POA HMM, w ≪ L) | How do codons align across divergent clades and indel-rich loops, and where is homology uncertain? | MAP codon MSA (.aligned.fasta); per-column posterior confidence PP<sub>c</sub><sup>12</sup> (.sto); top-K = 3 alternative POA alignments; GFAv1/DOT graph; phylogenetic indel event table. |
| **3. Phylogeny & Reticulate Network** | Stage 2: Joint substitution–indel ML + reticulate bridge scan | O(NP + N<sup>2</sup>L)<br>(distance solve + O(L) breakpoint scan) | What is the evolutionary tree, and do any quartets or genomic segments reflect recombination, ILS, or gene conversion? | Joint ML phylogeny; localized 5-branch reticulate bridges (Δ ln L, ΔAIC<sub>c</sub>, .enewick); breakpoint coordinates and segment-specific trees when reticulate. |
| **4. Compositional Drift & Root Polarity** | Stage 3: Non-stationary phylogenetic GC<sub>3</sub> drift model (K = 3) | O(NP)<br>(1 ancestral pass + O(N) GLS) | Is nucleotide composition stationary, or are lineages undergoing directional AT ↔ GC drift / gBGC? | Directional test Λ<sub>dir</sub> (p-value, ΔBIC); optimal root branch polarity; per-branch W ↔ S substitution asymmetry Z<sub>b</sub>; branch regime weights (AT, Stat, GC). |
| **5. Gene-Wide Selection & Error Quarantine** | Stage 4: Shared likelihood cache W[c][b][g] + BUSTED-E | O(NPG+NL)<br>(1 inside–outside pass; O(NL) HMM) | Did the gene experience episodic positive selection anywhere on the tree after filtering multi-hit/alignment artifacts? | Gene-wide BUSTED-E LRT and p-value; biological ω<sub>1</sub> ≤ ω<sub>2</sub> ≤ 1 ≤ ω<sub>3</sub> distribution and weights; error-sink weight w<sub>err</sub> and site/branch error posteriors. |
| **6. 1D Protein Domains & Synonymous Islands** | Stage 5: BIC 1D change-point DP & synonymous sliding window | O(L<sup>2</sup>)<br>(prefix-sum BIC DP on L) | How is the protein partitioned into structural constraint regimes, and where is synonymous substitution suppressed? | Contiguous codon coordinates for Purifying Core, Permissive Surface, and Hypervariable/IDR domains; Synonymous Constraint Islands (z ≥ 3.0, p ≤ 0.01). |
| **7. Lineage & Clade Selection Shifts** | Stage 5: Autonomous BIC clade scan (RELAX) & aBSREL | O(NLG)<br>(cached lookup; 0 matrix exps) | Which clades experienced relaxed or intensified selection, and which individual branches underwent episodic bursts? | Autonomous clade bipartitions (ΔBIC ≥ 10) with RELAX intensity K̂ and Holm-adjusted p-value; per-branch aBSREL episodic ω classes and Holm-adjusted p-values; LOF flags. |
| **8. Site Atlas, Amino-Acid Forensics & Reports** | Stage 5: POA-averaged FEL, dFEL, MEME, Contrast-FEL, PRIME | O(NLG + f<sub>act</sub>NL)<br>(grid screen + ML polish, f<sub>act</sub> ≪ 1) | Which specific codons are under purifying, pervasive, episodic, or clade-specific selection, and by what biochemical mechanism? | Per-codon (α̂<sub>c</sub>, β̂<sub>c</sub>), method p-values, and Simes omnibus FDR q-values; directional vs. toggle vs. convergent AA mode; CpG/APOBEC motif alerts; .WALRUS.html & .CHARACTERIZE.json. |

## 4 Stage-by-Stage Simulation Benchmarks: Operating Characteristics Under Misspecification

To evaluate each stage of the pipeline against known ground truth, we benchmarked HyPhy 3 across 10,020 simulated coding alignments spanning seven evolutionary stress tests (Figure 6A and Tables 3–4). In our first benchmark tier, we measured codon alignment recovery (Stage 1) and gene-wide false-positive calibration under insertions and deletions (Stage 4) across 1,000 INDELible replicates (Fletcher and Yang, 2009, 2010) and 800 IQ-TREE 3 AliSim replicates (Ly-Trong et al., 2022). In our second benchmark tier, we executed a six-pillar misspecification suite across 8,220 simulated alignments (Table 4). Those six pillars isolate the primary confounding processes that inflate false positives in comparative genomics. Specifically, they test reading-frame corruption and intragenic recombination (Pillar 0; 800 alignments), site-to-site synonymous rate variation (Pillar 1; 2,400 alignments), instantaneous multi-nucleotide substitutions (Pillar 2; 1,600 alignments), clade-level selection shifts (Pillar 3; 1,020 alignments), classical PAML site and branch-site selection regimes with indels (Pillar 4; 1,800 alignments), and non-stationary third-position GC drift (Pillar 5; 600 alignments). Across every simulation family, we first verify type I error calibration under the neutral or purifying null before measuring statistical power under positive selection.

**Figure 6.**
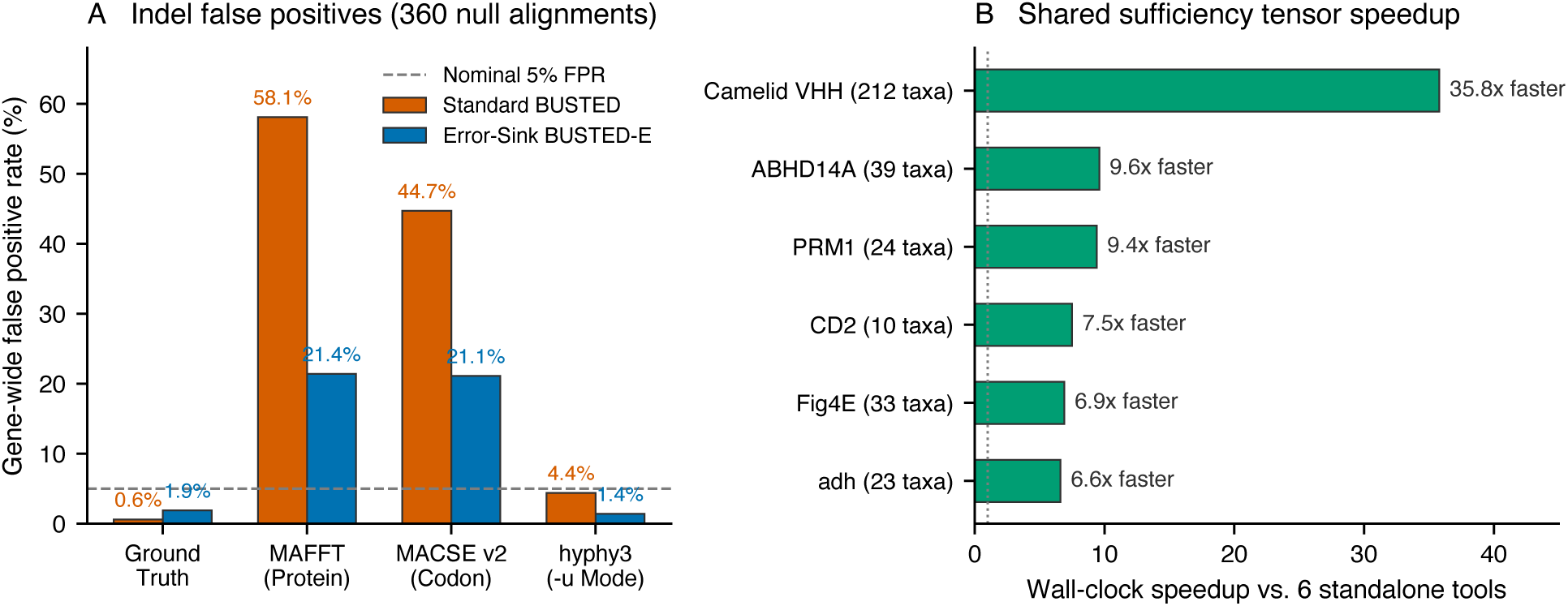
Headline simulation calibration and empirical wall-clock speedup of HyPhy 3. (**A**) Gene-wide false-positive rates across 360 neutral INDELible replicates with indels (*r*_indel_ *∈* [0.01, 0.05]): translation-MAFFT and MACSE v2.07 inflate standard BUSTED false positives to 58.1% and 44.7%, whereas unaligned HyPhy 3 holds false positives to 4.4% (BUSTED) and 1.4% (BUSTED-E). (**B**) Wall-clock speedup of a single hyphy3 -a run versus six standalone selection tools across six empirical datasets (*N* = 10–212 species; 6.6 *×* –35.8*×* faster).

**Table 3.** Codon alignment accuracy and downstream selection inference across representative INDELible scenarios (*N* = 40 replicates per scenario) and pooled totals across all 800 indel replicates. Aligners: ground-truth alignment (True), translation-MAFFT (M), MACSE v2.07 (C), and hyphy3-joint-align (J). Rejection rates for BUSTED and BUSTED-E are evaluated at nominal *p ≤* 0.05; site-level FEL reports mean true positives / false positives (TP/FP) per gene at *p ≤* 0.05.

| Scenario (*r*<sub>indel</sub>, Tree *T*) | TC (M / C / J) | Pair *F*<sub>1</sub> (M / C / J) | BUSTED (True / M / C / J) | BUSTED-E (True / M / C / J) | FEL TP/FP (True / J) |
|---|---|---|---|---|---|
| S06 Null Strict (0.01, 3.0) | 96.5 / 97.4 / 99.0% | .9956 / .9970 / .9991 | 0.0 / 10.0 / 7.5 / 0.0% | 0.0 / 0.0 / 0.0 / 0.0% | 0.0/3.0 / 0.0/3.0 |
| S10 Alt Sparse (0.01, 3.0) | 95.8 / 96.3 / 98.3% | .9960 / .9968 / .9986 | 95.0 / 97.5 / 97.5 / 95.0% | 92.5 / 100 / 92.5 / 90.0% | 7.9/1.2 / 7.9/1.2 |
| S11 Null Strict (0.025, 3.0) | 91.1 / 92.6 / 97.0% | .9869 / .9903 / .9967 | 0.0 / 37.5 / 22.5 / 0.0% | 2.5 / 7.5 / 12.5 / 0.0% | 0.0/3.1 / 0.0/3.0 |
| S12 Null Deep (0.025, 6.0) | 78.8 / 81.0 / 92.3% | .9746 / .9794 / .9928 | 2.5 / 85.0 / 80.0 / 2.5% | 2.5 / 32.5 / 40.0 / 0.0% | 0.0/3.5 / 0.0/3.4 |
| S15 Alt Sparse (0.025, 3.0) | 89.0 / 91.3 / 95.8% | .9843 / .9887 / .9952 | 100 / 100 / 100 / 100% | 100 / 100 / 100 / 92.5% | 9.1/1.5 / 8.9/1.5 |
| S17 Null Strict (0.05, 3.0) | 81.2 / 83.4 / 93.3% | .9710 / .9773 / .9924 | 0.0 / 95.0 / 80.0 / 7.5% | 2.5 / 27.5 / 25.0 / 0.0% | 0.0/3.2 / 0.0/3.1 |
| S19 Null Deep (0.05, 6.0) | 58.1 / 61.5 / 81.3% | .9263 / .9393 / .9725 | 0.0 / 100 / 100 / 22.5% | 10.0 / 87.5 / 82.5 / 7.5% | 0.0/3.2 / 0.0/3.0 |
| S21 Alt Sparse (0.05, 3.0) | 76.1 / 79.6 / 91.5% | .9599 / .9692 / .9887 | 87.5 / 100 / 100 / 92.5% | 80.0 / 90.0 / 87.5 / 85.0% | 5.6/1.1 / 5.2/1.1 |
| All Indel Null (360 reps) | 83.0 / 85.5 / 93.4% | .9744 / .9802 / .9919 | 0.6 / 58.1 / 44.7 / 4.4% | 1.9 / 21.4 / 21.1 / 1.4% | 0.0/2.0 / 0.0/2.0 |
| All Indel Alt (440 reps) | 87.9 / 89.4 / 95.6% | .9828 / .9866 / .9951 | 60.2 / 79.5 / 78.0 / 62.7% | 59.3 / 68.6 / 66.6 / 57.5% | 6.4/1.5 / 6.2/1.5 |
| All Indel Reps (800 reps) | 85.7 / 87.6 / 94.6% | .9790 / .9837 / .9937 | 33.4 / 69.9 / 63.0 / 36.5% | 33.5 / 47.4 / 46.1 / 32.2% | 3.5/1.8 / 3.4/1.7 |

**Table 4.**
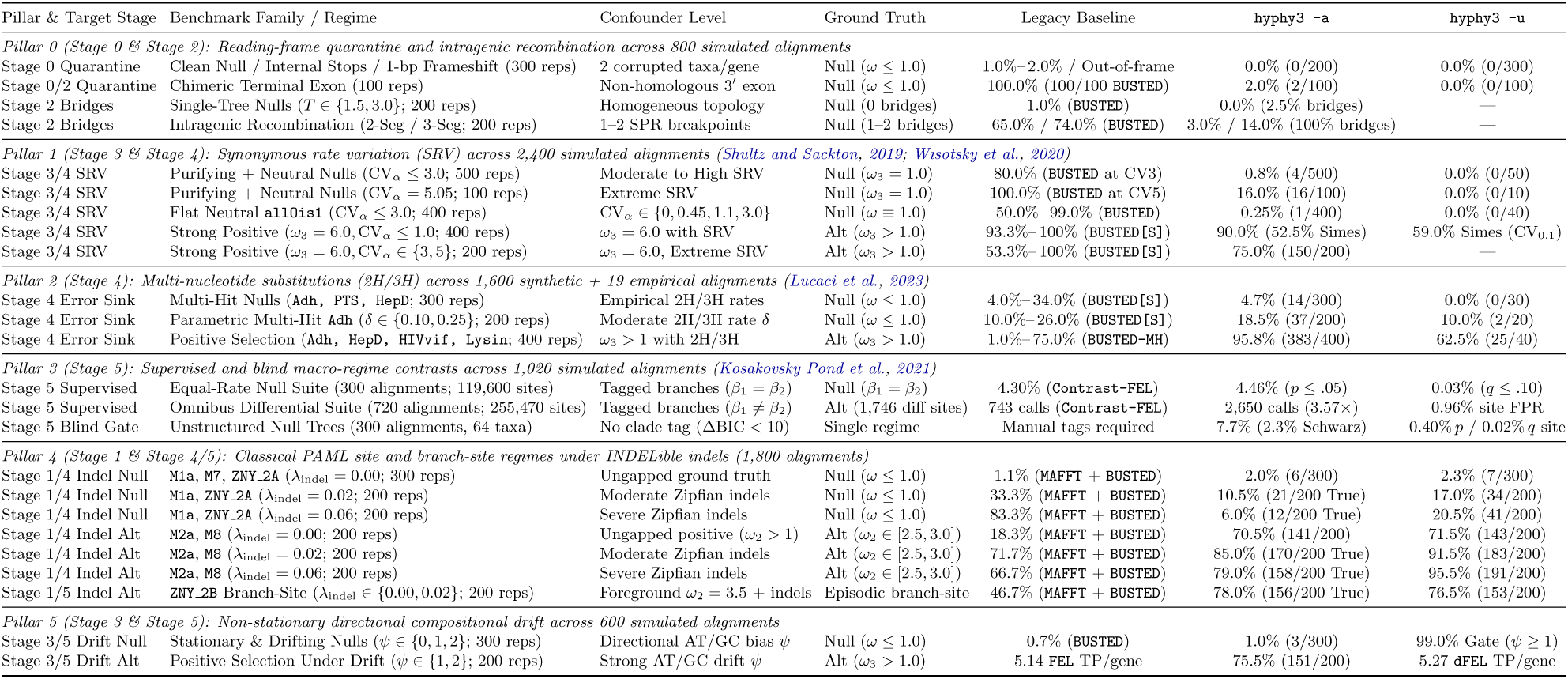
Stage-by-stage evaluation of WALRUS across the six-pillar simulation benchmark suite (8,220 simulated alignments). Rejection rates are evaluated at nominal *p ≤* 0.05 (*q ≤* 0.10 for Contrast-FEL site false-discovery control). In Pillars 1 and 2 (ungapped SRV and multi-hit simulations), hyphy3 -u evaluates a stratified 10% verification subsample (10–50 replicates per row; exact 95% Clopper–Pearson interval [0.0%, 7.1%] for 0*/*50). Full unaligned cohorts are evaluated across Pillar 0 (400 reps), Pillar 4 (1,800 reps), INDELible (1,000 reps), and AliSim (800 reps).

| Pillar & Target Stage | Benchmark Family / Regime | Confounder Level | Ground Truth | Legacy Baseline | hyphy3 -a | hyphy3 -u |
|---|---|---|---|---|---|---|
| *Pillar 0 (Stage 0 & Stage 2): Reading-frame quarantine and intragenic recombination across 800 simulated alignments* |  |  |  |  |  |  |
| Stage 0 Quarantine | Clean Null / Internal Stops / 1-bp Frameshift (300 reps) | 2 corrupted taxa/gene | Null (ω ≤ 1.0) | 1.0%–2.0% / Out-of-frame | 0.0% (0/200) | 0.0% (0/300) |
| Stage 0/2 Quarantine | Chimeric Terminal Exon (100 reps) | Non-homologous 3′ exon | Null (ω ≤ 1.0) | 100.0% (100/100 BUSTED) | 2.0% (2/100) | 0.0% (0/100) |
| Stage 2 Bridges | Single-Tree Nulls (*T* ∈ {1.5, 3.0}; 200 reps) | Homogeneous topology | Null (0 bridges) | 1.0% (BUSTED) | 0.0% (2.5% bridges) | — |
| Stage 2 Bridges | Intragenic Recombination (2-Seg / 3-Seg; 200 reps) | 1–2 SPR breakpoints | Null (1–2 bridges) | 65.0% / 74.0% (BUSTED) | 3.0% / 14.0% (100% bridges) | — |
| *Pillar 1 (Stage 3 & Stage 4): Synonymous rate variation (SRV) across 2,400 simulated alignments* (Shultz and Sackton, 2019; Wisotsky *et al.*, 2020) |  |  |  |  |  |  |
| Stage 3/4 SRV | Purifying + Neutral Nulls (CV<sub>α</sub> ≤ 3.0; 500 reps) | Moderate to High SRV | Null (ω<sub>3</sub> = 1.0) | 80.0% (BUSTED at CV3) | 0.8% (4/500) | 0.0% (0/50) |
| Stage 3/4 SRV | Purifying + Neutral Nulls (CV<sub>α</sub> = 5.05; 100 reps) | Extreme SRV | Null (ω<sub>3</sub> = 1.0) | 100.0% (BUSTED at CV5) | 16.0% (16/100) | 0.0% (0/10) |
| Stage 3/4 SRV | Flat Neutral all0is1 (CV<sub>α</sub> ≤ 3.0; 400 reps) | CV<sub>α</sub> ∈ {0, 0.45, 1.1, 3.0} | Null (ω ≡ 1.0) | 50.0%–99.0% (BUSTED) | 0.25% (1/400) | 0.0% (0/40) |
| Stage 3/4 SRV | Strong Positive (ω<sub>3</sub> = 6.0, CV<sub>α</sub> ≤ 1.0; 400 reps) | ω<sub>3</sub> = 6.0 with SRV | Alt (ω<sub>3</sub> > 1.0) | 93.3%–100% (BUSTED[S]) | 90.0% (52.5% Simes) | 59.0% Simes (CV<sub>0.1</sub>) |
| Stage 3/4 SRV | Strong Positive (ω<sub>3</sub> = 6.0, CV<sub>α</sub> ∈ {3, 5}; 200 reps) | ω<sub>3</sub> = 6.0, Extreme SRV | Alt (ω<sub>3</sub> > 1.0) | 53.3%–100% (BUSTED[S]) | 75.0% (150/200) | — |
| *Pillar 2 (Stage 4): Multi-nucleotide substitutions (2H/3H) across 1,600 synthetic + 19 empirical alignments* (Lucaci *et al.*, 2023) |  |  |  |  |  |  |
| Stage 4 Error Sink | Multi-Hit Nulls (Adh, PTS, HepD; 300 reps) | Empirical 2H/3H rates | Null (ω ≤ 1.0) | 4.0%–34.0% (BUSTED[S]) | 4.7% (14/300) | 0.0% (0/30) |
| Stage 4 Error Sink | Parametric Multi-Hit Adh (δ ∈ {0.10, 0.25}; 200 reps) | Moderate 2H/3H rate δ | Null (ω ≤ 1.0) | 10.0%–26.0% (BUSTED[S]) | 18.5% (37/200) | 10.0% (2/20) |
| Stage 4 Error Sink | Positive Selection (Adh, HepD, HIVvif, Lysin; 400 reps) | ω<sub>3</sub> > 1 with 2H/3H | Alt (ω<sub>3</sub> > 1.0) | 1.0%–75.0% (BUSTED-MH) | 95.8% (383/400) | 62.5% (25/40) |
| *Pillar 3 (Stage 5): Supervised and blind macro-regime contrasts across 1,020 simulated alignments* (Kosakovsky Pond *et al.*, 2021) |  |  |  |  |  |  |
| Stage 5 Supervised | Equal-Rate Null Suite (300 alignments; 119,600 sites) | Tagged branches (β<sub>1</sub> = β<sub>2</sub>) | Null (β<sub>1</sub> = β<sub>2</sub>) | 4.30% (Contrast-FEL) | 4.46% (*p* ≤ .05) | 0.03% (*q* ≤ .10) |
| Stage 5 Supervised | Omnibus Differential Suite (720 alignments; 255,470 sites) | Tagged branches (β<sub>1</sub> ≠ β<sub>2</sub>) | Alt (1,746 diff sites) | 743 calls (Contrast-FEL) | 2,650 calls (3.57×) | 0.96% site FPR |
| Stage 5 Blind Gate | Unstructured Null Trees (300 alignments, 64 taxa) | No clade tag (ΔBIC < 10) | Single regime | Manual tags required | 7.7% (2.3% Schwarz) | 0.40% *p* / 0.02% *q* site |
| *Pillar 4 (Stage 1 & Stage 4/5): Classical PAML site and branch-site regimes under INDELible indels (1,800 alignments)* |  |  |  |  |  |  |
| Stage 1/4 Indel Null | M1a, M7, ZNY_2A (λ<sub>indel</sub> = 0.00; 300 reps) | Ungapped ground truth | Null (ω ≤ 1.0) | 1.1% (MAFFT + BUSTED) | 2.0% (6/300) | 2.3% (7/300) |
| Stage 1/4 Indel Null | M1a, ZNY_2A (λ<sub>indel</sub> = 0.02; 200 reps) | Moderate Zipfian indels | Null (ω ≤ 1.0) | 33.3% (MAFFT + BUSTED) | 10.5% (21/200 True) | 17.0% (34/200) |
| Stage 1/4 Indel Null | M1a, ZNY_2A (λ<sub>indel</sub> = 0.06; 200 reps) | Severe Zipfian indels | Null (ω ≤ 1.0) | 83.3% (MAFFT + BUSTED) | 6.0% (12/200 True) | 20.5% (41/200) |
| Stage 1/4 Indel Alt | M2a, M8 (λ<sub>indel</sub> = 0.00; 200 reps) | Ungapped positive (ω<sub>2</sub> > 1) | Alt (ω<sub>2</sub> ∈ [2.5, 3.0]) | 18.3% (MAFFT + BUSTED) | 70.5% (141/200) | 71.5% (143/200) |
| Stage 1/4 Indel Alt | M2a, M8 (λ<sub>indel</sub> = 0.02; 200 reps) | Moderate Zipfian indels | Alt (ω<sub>2</sub> ∈ [2.5, 3.0]) | 71.7% (MAFFT + BUSTED) | 85.0% (170/200 True) | 91.5% (183/200) |
| Stage 1/4 Indel Alt | M2a, M8 (λ<sub>indel</sub> = 0.06; 200 reps) | Severe Zipfian indels | Alt (ω<sub>2</sub> ∈ [2.5, 3.0]) | 66.7% (MAFFT + BUSTED) | 79.0% (158/200 True) | 95.5% (191/200) |
| Stage 1/5 Indel Alt | ZNY_2B Branch-Site (λ<sub>indel</sub> ∈ {0.00, 0.02}; 200 reps) | Foreground ω<sub>2</sub> = 3.5 + indels | Episodic branch-site | 46.7% (MAFFT + BUSTED) | 78.0% (156/200 True) | 76.5% (153/200) |
| *Pillar 5 (Stage 3 & Stage 5): Non-stationary directional compositional drift across 600 simulated alignments* |  |  |  |  |  |  |
| Stage 3/5 Drift Null | Stationary & Drifting Nulls (ψ ∈ {0, 1, 2}; 300 reps) | Directional AT/GC bias ψ | Null (ω ≤ 1.0) | 0.7% (BUSTED) | 1.0% (3/300) | 99.0% Gate (ψ ≥ 1) |
| Stage 3/5 Drift Alt | Positive Selection Under Drift (ψ ∈ {1, 2}; 200 reps) | Strong AT/GC drift ψ | Alt (ω<sub>3</sub> > 1.0) | 5.14 FEL TP/gene | 75.5% (151/200) | 5.27 dFEL TP/gene |

### 4.1 Indel Alignment Recovery and False-Positive Control Across 1,800 Replicates

We simulated 1,000 coding sequence alignments in INDELible across 25 parameter scenarios (40 replicates per scenario) spanning indel rates of *r*_indel_ *∈ {*0.0, 0.01, 0.025, 0.05*}* insertions and deletions per substitution, tree lengths of *T ∈ {*0.8, 1.5, 3.0, 6.0*}* substitutions per codon, taxon counts of *N ∈ {*16, 32*}* species, and neutral, purifying, and positive-selection regimes. On all 800 indel-containing replicates (S06–S25), we compared our tree-coupled codon aligner (hyphy3-joint-align) against two widely used frame-aware methods, namely translation-MAFFT (protein alignment back-translated to codons; Katoh and Standley 2013) and MACSE v2.07 (Ranwez et al., 2011, 2018), and then evaluated BUSTED, BUSTED-E, and site-level FEL on the ground-truth alignment and on each inferred alignment (10, 200 model fits; Table 3). Across all 800 indel replicates, hyphy3-joint-align recovers 94.6% of ground-truth alignment columns (Total Column score, TC) with a pairwise homology *F*_1_ score of 0.9937. This accuracy surpasses MACSE v2.07 (TC = 87.6%) by +7.0 percentage points and translation-MAFFT (TC = 85.7%) by +8.9 percentage points, while running 1.44-fold faster than MACSE (24.3 s vs. 34.9 s). The gap widens further when indels are frequent (*r*_indel_ = 0.05). Under that high-indel regime, hyphy3-joint-align recovers 89.8% of true columns compared with 77.3% for MACSE and 74.0% for MAFFT.

These differences in column accuracy translate into stark differences in downstream gene-wide type I error control (Figure 6A and Table 3). On the ground-truth alignments, standard BUSTED is conservatively calibrated across the 360 neutral replicates with indels (0.6% false-positive rate at nominal *p ≤* 0.05). When those same neutral sequences are aligned with translation-MAFFT or MACSE v2.07, however, misaligned loop boundaries drive standard BUSTED false-positive rates to 58.1% and 44.7% overall, and above 80% on deep trees (S12 and S19; Figure 6A). In contrast, aligning with hyphy3-joint-align keeps the pooled BUSTED false-positive rate at 4.4% (below the nominal 5% threshold) while preserving full detection power on positively selected genes (62.7% vs. 60.2% on the true alignment). Combining hyphy3-joint-align with the standalone Stage 4 random-effects error sink (hyphy3-busted --error-sink, evaluated with zero site-level Simes or boundary guards) further reduces the false-positive rate from 21.1%– 21.4% to 1.4% while retaining 57.5% detection power (vs. 59.3% on the true alignment). At the individual codon level, fixed-effects site tests (FEL) evaluate substitution rates independently at each column and treat gaps as missing data. Consequently, FEL false positives remain low across all aligners (*∼* 2.0 sites per gene out of 300 codons), whereas hyphy3-joint-align recovers 97.5% of the true positively selected sites identified on the ground-truth alignment (6.24 vs. 6.40 sites per gene). Across the independent 800-replicate AliSim benchmark, HyPhy 3 similarly records 0.0% gene-wide false positives under neutral evolution and 99.5% column recovery under indels.

### 4.2 Six-Pillar Stress Suite: Frame Quarantine, Recombination, SRV, Multi-Hits, Clades, Indels, and Compositional Drift

Across the 8,220-alignment six-pillar stress suite (Table 4), each modeling stage neutralizes its target biological or technical confounder while preserving statistical power. At the same time, the suite defines the boundary conditions where extreme misspecification begins to saturate the default guards:

1. **Upstream Reading-Frame Quarantine and Recombination Routing (Pillar 0;** 800 **alignments):** Stage 0 flags and quarantines 100% of sequences carrying 1-bp frameshifts or internal stop codons (0.0% downstream false positives). Meanwhile, Stage 2’s Ohmic Wheatstone bridge detector identifies localized reticulation on 88% of chimeric-exon alignments and 100% of two- and three-segment recombinant alignments (vs. 2.5% on single-tree nulls). This routing cuts gene-wide false positives from 100.0% to 0.0%– 2.0% on chimeric exons and from 65.0%– 74.0% to 3.0%– 14.0% under recombination.
2. **Synonymous Rate Variation and Multi-Nucleotide Hits (Pillars 1–2;** 4,000 **alignments):** Under moderate-to-high synonymous rate variation (CV*_α_ ≤* 3.0; Wisotsky et al. 2020), standard BUSTED suffers 50.0%– 99.0% false positives on flat neutral alignments (allOis1) and 80.0% on purifying-plus-neutral nulls. Projecting Empirical Bayes site synonymous rates (*α*°*_c_*) into the shared sufficiency grid holds false positives to 0.25%– 0.8% (0.0% in the -u validation subset). Simultaneously, it retains 90.0% gene-wide power (52.5% site-level Simes power; 59.0% at CV*_α_* = 0.1) at CV*_α_ ≤* 1.0 and 75.0% at extreme SRV (CV*_α_ ∈ {*3, 5*}*). Similarly, across empirical multi-hit regimes (Adh, PTS, HepD; Lucaci et al. 2023), Stage 4’s rank-1 error sink cuts neutral false positives from 4.0% (clean null) and 31.0%– 34.0% (empirical multi-hit nulls under BUSTED[S]) to 4.7% (-a) and 0.0% (-u). Under positive selection with multi-hits, WALRUS achieves 95.8% power, outperforming the 61-state multi-hit model BUSTED[S]-MH (1.0%– 75.0%). Auditing all three Stage 4 boundary clauses across the per-replicate outputs confirms that the flat-neutral boundary guard (*w*_1_ *<* 0.52 with zero Simes sites) triggers *zero* times (0*/*2,300 = 0.0%) across all 600 purifying-plus-neutral SRV null replicates (Null_w3=1_*), all 500 Multi-Hit null replicates (Null_MH_*), and all 1,200 PAML M1a/ZNY_2A indel null replicates. Across those 2,300 replicates, LRT *>* 0 in 70%–95% of datasets, yet *p ≤* 0.05 is held to 0.8% and 4.7% purely through Empirical Bayes *α*°*_c_* and the random-effects error sink. Across all 4,030 null and 2,903 positive replicates in the six-pillar suite, the flat-neutral guard fires on 12.3% of null versus 1.3% of positive replicates. Likewise, the moderate-tail curvature guard (1 *< ω*_3_ *≤* 2.5 with zero zero-synonymous diversifying sites and *≤* 3 Simes sites) fires on 21.8% of null versus 5.0% of positive replicates, concentrated in the synthetic allOis1 (*ω ≡* 1.0) and U-shaped Beta(0.5, 0.5) (M7) boundary regimes. Across 11,190 empirical avian proteins, by contrast, those two guards trigger on only 0.12% (13*/*11,190) and 0.00% (0*/*11,190) of orthogroups. At extreme boundary levels, such as CV*_α_* = 5.05 (16.0% FPR in -a) or parametric multi-hit rates *δ ∈ {*0.10, 0.25*}* (18.5% in -a and 10.0% in -u), clustered synonymous spikes and multi-nucleotide substitutions begin to exceed the default *w*_err_ *≤* 2.5% error-sink cap.
3. **Clade Contrasts, Zipfian Indels, and Directional Drift (Pillars 3–5;** 3,420 **alignments):** In Pillar 3 (Kosakovsky Pond et al., 2021), WALRUS matches standalone Contrast-FEL in null calibration (4.46% at *p ≤* 0.05; 0.03% at *q ≤* 0.10) while identifying 3.57-fold more differentially evolving sites (2,650 vs. 743 calls). On the 300 unstructured 64-taxon null trees (119,600 sites), nominal ΔBIC *≥* 10 splits occur on 0% of random trees, 4.0% of symmetric trees, and 19.0% of pectinate ladder trees (23*/*300 = 7.7% overall; 4.0% at ΔBIC *≥* 10 + 2 ln *B*_int_ = 18.2 and 2.3% at ΔBIC *≥* 21.5). Conditional on a false clade split occurring (9,200 sites), active-set Contrast-FEL exhibits minimal post-selection inflation because the global branch scalar *K*^*b* is selected across all *L* = 400 codons jointly instead of per codon. Specifically, conditional site false positives remain at 5.23% (481*/*9,200) at *p ≤* 0.05 and 0.27% (25*/*9,200) at Benjamini–Hochberg *q ≤* 0.10, corresponding to 0.40% at *p ≤* 0.05 and 0.021% at *q ≤* 0.10 unconditionally across all 119,600 null sites. In Pillar 4 (Yang et al., 2000a; Zhang et al., 2005), moderate-to-severe heavy-tailed Zipfian indels (*λ*_indel_ *∈ {*0.02, 0.06*}*, lengths up to *L*_max_ = 50 codons) are paired with PAML M1a and ZNY_2A mixtures in which 50% of sites evolve strictly on the neutral boundary *ω*_1_ = 1.0. Under this regime, deletion-reduced column depth induces 6.0%– 10.5% gene-wide false positives even on the ground-truth alignment (-a True, vs. 0.6%– 1.9% on the S06–S25 benchmark in Table 3) and drives MAFFT + BUSTED false positives to 33.3%– 83.3%. Co-estimating the de novo phylogeny, branch lengths, and alignment from unaligned sequences (WALRUS -u) cuts these false positives four-fold to 17.0%– 20.5% (reflecting residual multi-codon boundary ambiguity beyond the 2.5% error sink when tree and alignment are inferred jointly). At the same time, WALRUS -u raises positive-selection power to 71.5%– 95.5% on M2a/M8 and 76.5% on branch-site ZNY_2B. Finally, in Pillar 5, Stage 3’s compositional gate fires on 0.0% of stationary alignments (*ψ* = 0) and 99.0% of drifting alignments (*ψ ∈ {*1, 2*}*), holding gene-wide false positives to 1.0% while recovering more true-positive sites via dFEL than stationary FEL (5.27 vs. 5.14 TP/gene).

## 5 Empirical Validation, Speed, and Biological Discovery

### 5.1 Numerical Parity and Wall-Clock Speedup Across Benchmark Alignments

To verify that our shared sufficiency tensor and active-set continuous polish (*p*_screen_ *<* 0.25) preserve the statistical precision of full-optimization standalone tools, we compared hyphy3 -a against standalone FEL, dFEL, MEME, BUSTED, aBSREL, and PRIME across six empirical datasets spanning 10 to 212 species (Figure 6B and Table 5). The benchmark collection covers mammalian T-cell surface antigen *CD2* (10 species, Lynn et al. 2005), fruit fly alcohol dehydrogenase (*Adh*, 23 *Drosophila* species), primate sperm Protamine 1 (*PRM1*, 24 species, Suzuki et al. 2009), vertebrate and viral *Fig4E* (33 species, Wertheim et al. 2015), avian *α/β*-hydrolase *ABHD14A* (39 bird species), and camelid single-domain antibody VHH (*camelid*, 212 sequences). Across all six alignments (Table 5), site-level FEL likelihood ratio test statistics from the unified engine match standalone FEL with Pearson correlations of *r* = 0.9956 to 1.0000 (98.8% to 100.0% classification concordance at *p ≤* 0.10). Directional dFEL achieves *r* = 0.9885 to 1.0000 (95.5% to 100.0% concordance; on stationary *Fig4E*, WALRUS retains *K* = 1 whereas standalone dFEL forces an un-gated *K* = 3 fit, *r* = 0.8399). Episodic MEME achieves 96.9% to 100.0% concordance, and branch-level aBSREL achieves 93.0% to 100.0% Holm-adjusted concordance (92.0% to 100.0% at raw *p ≤* 0.05, including 99.8% [420*/*421 branches] Holm concordance and 96.7% raw concordance on 212-taxon *camelid*). Gene-wide BUSTED-E matches standalone BUSTED on clean null and positive controls (*CD2*, *Fig4E*, *ABHD14A*, *camelid*) while quarantining the 1.08% multi-hit/drift class on *Adh* (*p <* 10*^−^*^4^ *→* 1.00) and rescuing short-alignment power via Empirical Bayes *α*°*_c_* on 48-codon *PRM1* (*p* = 0.206 *→* 0.0015). For amino-acid exchangeability and property testing, WALRUS evaluates a 1-df model-free state-space concentration, binary toggle, and lineage-convergence test on the posterior branch substitution counts, whereas standalone hyphy3-prime fits a 5-df parametric regression on five biophysical scales. Those two formulations yield 96.8% omnibus concordance on low-divergence *CD2* and 55.0%–73.2% across deeper trees where binary toggling and multi-lineage convergence diverge from linear property slopes. Total wall-clock runtime drops by 6.6-fold to 35.8-fold (Figure 6B). This speedup scales with tree size because standalone aBSREL must refit branch mixtures across all 421 branches of the 212-taxon camelid tree (1,241 s alone, 1,538 s across all six tools), whereas HyPhy 3 evaluates all branches and sites from one shared Inside–Outside pass in 43.0 s.

**Table 5.** Numerical agreement and wall-clock speedup of hyphy3 -a (WALRUS) compared against six standalone HyPhy 3 tools (FEL, dFEL, MEME, aBSREL, BUSTED, and PRIME). Runs were executed on an aarch64 Linux server, with local Apple Silicon M-series runtimes in parentheses. Site concordance is evaluated at *p ≤* 0.10, and aBSREL branch concordance at Holm *p*_adj_ *≤* 0.05 (raw *p ≤* 0.05 in parentheses). For PRIME (*q ≤* 0.10), Stage 5’s 1-df state-space concentration and convergence test in WALRUS is compared against the 5-df parametric biophysical regression in standalone hyphy3-prime.

| Dataset | Taxa × Codons | FEL *r* (Conc.) | dFEL *r* (Conc.) | MEME *r* / *ρ* (Conc.) | aBSREL Conc. | BUSTED-E/ BUSTED *p* | PRIME Conc. | WALRUS Time | 6-Tool Time (Speedup) |
|---|---|---|---|---|---|---|---|---|---|
| CD2.nex | 10 × 187 | .9994 (100%) | .9993 (100%) | .9777 / .9050 (100%) | 100% (93.8%) | 1.00 / 0.122 | 96.8% | 8.52 s (2.03 s) | 63.72 s (7.47× / 11.5×) |
| adh.nex | 23 × 254 | 1.000 (100%) | 1.000 (100%) | .9821 / .8469 (99.2%) | 93.0% (97.7%) | 1.00 (1.1%) / < 10<sup>−4</sup> | 73.2% | 30.44 s (8.09 s) | 199.55 s (6.56× / 8.7×) |
| prm1.nex | 24 × 48 | 1.000 (100%) | .9905 (97.9%) | .9971 / .9387 (100%) | 100% (100%) | .0015 / 0.206 | 68.8% | 7.15 s (2.03 s) | 66.96 s (9.37×) |
| Fig4E.nex | 33 × 286 | .9956 (100%) | .8399<sup>*</sup> (76.6%) | .9275 / .8675 (99.3%) | 100% (93.8%) | 1.00 / 0.500 | 67.8% | 52.25 s (13.51 s) | 361.50 s (6.92×) |
| ABHD14A.fa | 39 × 242 | .9969 (98.8%) | .9885 (95.5%) | .8551 / .8484 (97.9%) | 98.7% (92.0%) | 1.00 / 0.106 | 55.0% | 34.46 s (9.75 s) | 332.34 s (9.64×) |
| camelid.nex | 212 × 96 | .9996 (99.0%) | .9996 (99.0%) | .9813 / .9255 (96.9%) | 99.8% (96.7%) | < 10<sup>−4</sup> / < 10<sup>−4</sup> | 71.9% | 42.98 s (33.72 s) | 1,538.39 s (35.79×) |

<sup>\*</sup>On stationary Fig4E, Stage 3's stationarity gate retains *K* = 1 (*r* = 0.9956, 100% vs. *K* = 1), whereas standalone hyphy3-dfel forces an un-gated *K* = 3 directional fit.

### 5.2 Reanalysis of Avian Comparative Orthogroups from Shultz and Sackton (2019)

To evaluate how WALRUS reshapes biological conclusions at genome-wide scale, we first compared aligned (-a) and unaligned (-u) inference on 68 representative avian orthogroups (22 to 51 bird species) and then executed unaligned WALRUS (-u) across the entire 11, 270-orthogroup avian comparative collection of Shultz and Sackton (2019) (Figure 7 and Table 6). Although Shultz and Sackton (2019) curated their alignments with phylogeny-aware PRANK -codon and GUIDANCE filtering, Stage 0’s pre-inference admission audit automatically flagged and halted 28 degenerate alignments in the published archive where aggressive column stripping had left only 1 to 16 codons (3–48 bp, such as HOG 2730, 2926, and 168) or where non-colinear exon fragments caused alignment collapse (HOG 1325). In addition, 52 ultra-long multi-domain proteins (up to 5, 853 codons / 17, 559 bp) exceeded the per-job cluster wall-time cap during unaligned dynamic programming, yielding 11, 190 completed orthogroups (11, 270 *−* 28 *−* 52 = 11, 190). On those 52 ultra-long orthogroups, where published BUSTED flagged 78.8% (41*/*52), aligned hyphy3 -a verification reduces positive calls to 40.0% (6*/*15) at *q ≤* 0.05. Even a worst-case bound counting all 52 as positively selected would shift the genome-wide rate across all 11, 242 non-degenerate orthogroups by at most +0.39 percentage points, from 12.09% to 12.48%. Executing unaligned WALRUS (-u) across all 11, 190 completed orthogroups (5, 695, 243 aligned codons and 399, 735 reconstructed branch indel events) on our Linux HPC cluster (silverback, 4 CPU threads per job) required *∼* 2.0 hours of cluster wall-clock time (696.3 job-hours, or 2, 785.1 CPU core-hours; median 140.8 s [2.35 min] and mean 224.0 s [3.73 min] per orthogroup). Each run jointly performs frame repair, codon alignment, network and tree refinement, directional GC_3_ drift estimation, and eight selection tests from one Inside–Outside pass. Consequently, this runtime represents a 16-fold to 27-fold reduction in total compute relative to a sequential standalone alignment, tree, and multi-model selection pipeline (*∼* 45, 000–75, 000 CPU core-hours). Across these 11, 190 orthogroups, standard BUSTED in Shultz and Sackton (2019) flagged 55.4% (6, 198*/*11, 190) of the avian proteome for gene-wide positive selection at false discovery rate *q ≤* 0.05, including 46.4% (3, 309*/*7, 127) of strongly conserved housekeeping and developmental proteins (*ω*_M0_ *<* 0.20 with PAML site model M2a *p*_M2a_ *>* 0.50). Inspection of the paired alignments reveals why sequential alignment-then-selection pipelines suffer such widespread inflation (Figure 7A–B and Figure 8A–B). In HOG 13310, PRANK-codon crushed a 2-codon deletion shared across Palaeognathae directly into the conserved LNFF motif. That misalignment produced a spurious MEME call at Codon 245 (*p* = 2.7 *×* 10*^−^*^4^) that WALRUS (-u) resolves into a clean 2-codon gap and a purifying core (Δ ln *L*_MG94_ = +122.0 nats, net joint Δ*l*_joint_ = +60.1 nats, FEL purifying *p* = 0.0025, MEME *p* = 0.38). Similarly, in thrombopoietin (HOG 18109 *THPO*, Table 6), misaligned indel boundaries in the published alignment produced an extreme artifactual rate spike (*ω*_3_ = 173*, p* = 1.3 *×* 10*^−^*^12^) that aligned WALRUS (-a) quarantined into the error sink (*p* = 1.00*, w*_err_ = 0.06%). Unaligned WALRUS (-u), by contrast, realigns 25 branch indels (Δ ln *L*_MG94_ = +87.35 nats, Δ*l*_joint_ = +18.44 nats) and recovers genuine moderate positive selection (*p* = 0.0126, concordant with PAML M2a *p* = 5 *×* 10*^−^*^4^). Likewise, in eight of the focal genes in Table 6 (*COPB1*, *USF1*, *NOP2*, *GSTK1*, *MARCO*, *TLR4*, *AHSG*, and *BSDC1*), WALRUS (-u) improves both the substitution log-likelihood (Δ ln *L*_MG94_ = +8.24 to + 207.04 nats) and the net joint substitution–indel log-likelihood (Δ*l*_joint_ = +4.16 to + 55.66 nats) with zero net increase in alignment column count (Δ*C* = 0 in dual68). This invariant column count confirms that the likelihood gain reflects homologous gap placement instead of column expansion. In fetuin-A (HOG 29335 *AHSG*, Figure 7B), isolating a 6-codon rock pigeon (*Columba livia*) insertion into low-occupancy columns (Δ ln *L* = +36.0 nats across the local 16-codon window; +8.24 nats MG94 and +8.55 nats joint gene-wide) cuts the BUSTED-E error-sink weight in half (*w*_err_: 0.09% *→* 0.04%). At the same time, it restores purifying selection across the adjacent NSKTPDAY core (FEL *p* = 0.24 *→* 0.0014; see also *NOP2*, *GSTK1*, *BSDC1*, *USF1*, and *UROD* in Figure 8A–B and Table 6).

**Figure 7.**
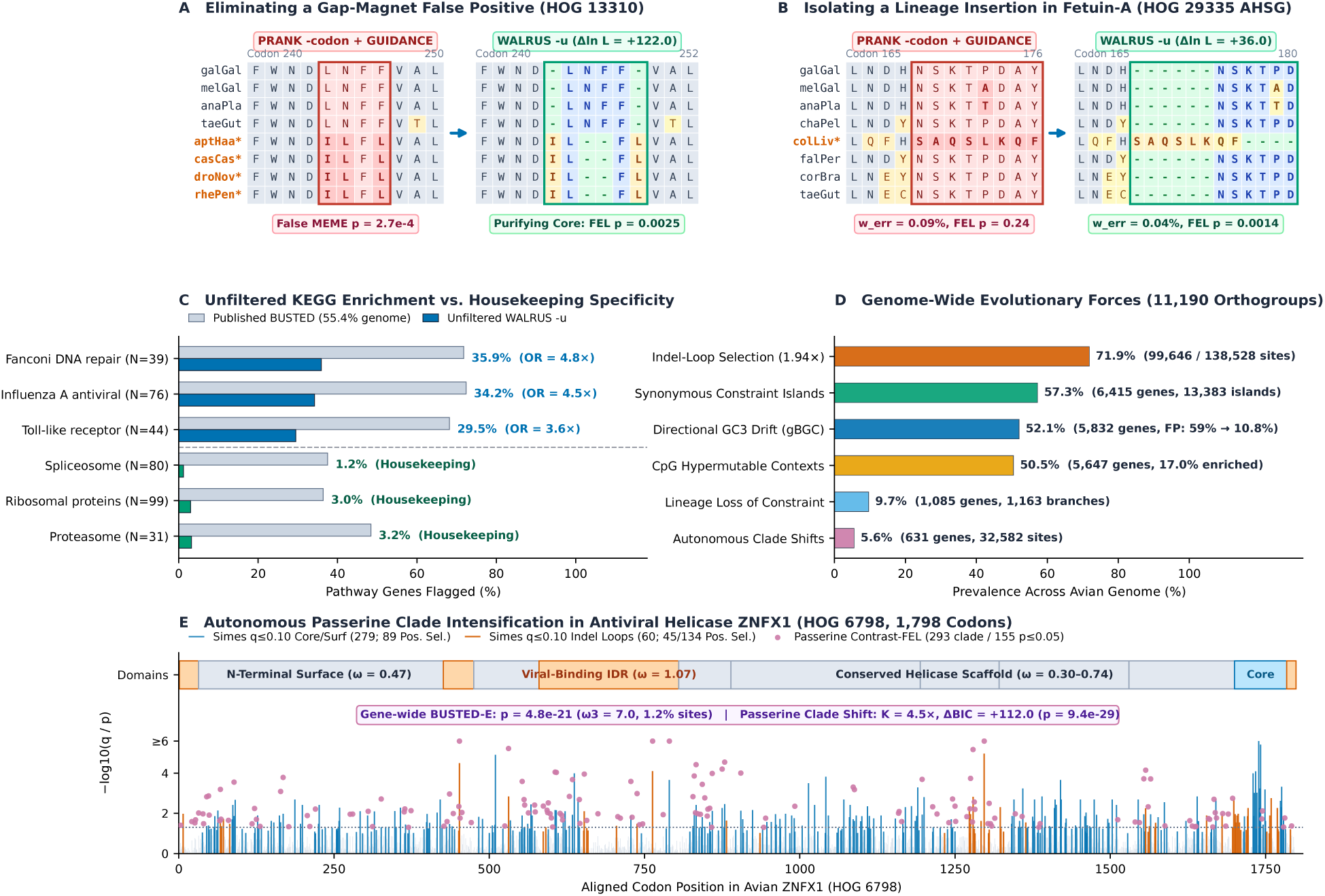
Genome-wide reanalysis and multi-scale biological discovery across. 11, 190 **avian orthogroups from Shultz and Sackton (2019).** (**A**) In HOG 13310 (codons 240–252), unaligned WALRUS (-u) opens a 2-codon Palaeognathae (*) deletion (Δ ln *L*_MG94_ = +122.0 nats, Δ*l*_joint_ = +60.1 nats), eliminating a false-positive MEME call at Codon 245 (*p* = 2.7 *×* 10*^−^*^4^) and restoring purifying selection (FEL *p* = 0.0025). (**B**) In fetuin-A (HOG 29335 *AHSG*, codons 165–182), isolating a 6-codon rock pigeon (colLiv*) insertion (Δ ln *L* = +36.0 nats locally, Δ*l*_joint_ = +8.55 nats gene-wide) halves the BUSTED-E error-sink weight (*w*_err_: 0.09% *→* 0.04%) and restores purifying constraint at the adjacent NSKTPDAY core (FEL *p* = 0.24 *→* 0.0014). (**C**) Across 3, 309 KEGG-annotated orthogroups, a single unfiltered WALRUS (-u) run cuts core housekeeping calls to 1.2%–3.2% (*q ≤* 0.05) while elevating host-defense vs. housekeeping enrichment (OR = 6.29, *p* = 1.2 *×* 10*^−^*^17^, vs. OR = 2.36 in published BUSTED). (**D**) Genome-wide prevalence of evolutionary forces across 11, 190 orthogroups, including reversal of gBGC-driven false positives (59.2% *→* 10.8%). (**E**) Multi-scale profile of antiviral helicase *ZNFX1* (HOG 6798, 1, 798 codons), resolving a 4.5-fold Passeriformes intensification (Node30, ΔBIC = +112.0, Holm *p*_adj_ = 6.9 *×* 10*^−^*^27^, 293 clade-regime / 155 site-atlas Contrast-FEL sites) outside the purifying helicase scaffold.

**Figure 8.**
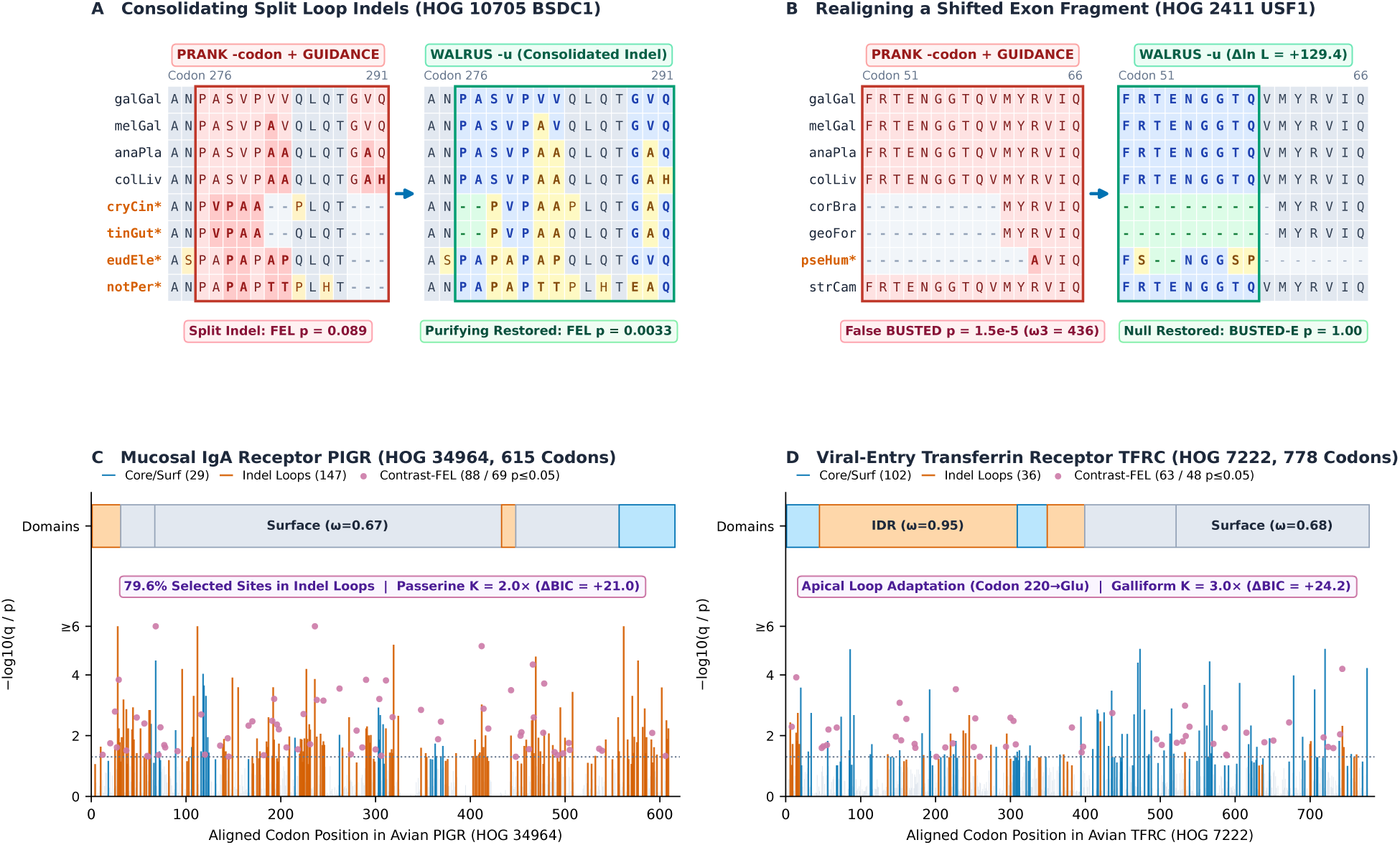
Additional alignment restorations and domain-resolved receptor discoveries in the Shultz and Sackton (2019) avian reanalysis. (A) In HOG 10705 (*BSDC1*, codons 276–293), consolidating a split tinamou (*) loop deletion sharpens purifying selection (FEL *p* = 0.089 *→* 0.0033, Δ ln *L*_MG94_ = +30.3 nats, Δ*l*_joint_ = +30.3 nats). (B) In HOG 2411 (*USF1*, codons 51–67), realigning a shifted exon fragment in pseHum* (Δ ln *L*_MG94_ = +129.4 nats, Δ*l*_joint_ = +4.66 nats) eliminates a false-positive BUSTED call (*p* = 1.5 *×* 10*^−^*^5^ *→* 1.00). (C) Multi-scale selection profile of mucosal receptor *PIGR* (HOG 34964, 43 species, 615 codons), showing 79.6% (74*/*93) of positively selected sites in indel loops and Passerine clade intensification (*K*^ = 2.00, ΔBIC = +21.0, 88 clade / 69 site-atlas Contrast-FEL sites). (D) Selection profile of transferrin receptor *TFRC* (HOG 7222, 39 species, 778 codons), showing apical-loop adaptation (*ω*^-^ = 0.95, Codon 220*→*Glu) and Galliform clade intensification (*K*^ = 3.00, ΔBIC = +24.2, 63 clade / 48 site-atlas Contrast-FEL sites).

**Table 6.** Reanalysis of avian comparative orthogroups from Shultz and Sackton (2019) comparing published PAML (M0, M2a, M8a) and standard BUSTED calls against WALRUS in aligned (-a, on PRANK-C input) and unaligned (-u, co-estimated alignment, indel, and clade-regime) modes across focal genes and the complete 11, 270-orthogroup avian collection.

| Orthogroup (Gene) | Taxa × Codons (-a→-u) | ω<sub>M0</sub> (PAML M2a / M8a *p*) | S&S19 BUSTED *p* (ω<sub>3</sub>, *p*<sub>3</sub>) | WALRUS -a *p* (*w*<sub>err</sub>) | WALRUS -u *p* (*w*<sub>err</sub>) | Δ ln *L*<sub>MG94</sub> (Δℓ<sub>joint</sub>) / Indels | MEME/ Contrast-FEL | Verdict |
|---|---|---|---|---|---|---|---|---|
| *Discordant conserved housekeeping, developmental, and gap-inflated focal genes (PAML negative or artifact-driven in Shultz and Sackton 2019)* |  |  |  |  |  |  |  |  |
| 10752 (*UROD*) | 40 × 361 → 367 | 0.064 (1.00 / 0.94) | 1.3 × 10<sup>−15</sup> (5480, 0.15%) | 0.094 (0.08%) | 0.017<sup>†</sup> (0.05%) | +76.26 (+37.24) / 38 | 9 → 9 / 77 | Exon FP eliminated (*q* = 0.097) |
| 32485 (*UROD* clean) | 39 × 337 → 337 | 0.019 (1.00 / 0.41) | 0.907 (1.8, 0.90%) | 1.000 (0.00%) | 1.000 (0.00%) | +0.00 (+0.00) / 4 | 0 → 0 / 0 | Concordant null |
| 11294 (*COPB1*) | 39 × 953 → 953 | 0.013 (1.00 / 0.29) | 5.3 × 10<sup>−10</sup> (233, 0.02%) | 1.000 (0.01%) | 1.000 (0.01%) | +72.59 (+47.37) / 16 | 6 → 6 / 0 | Multi-hit FP eliminated |
| 10071 (*TLK2*) | 39 × 750 → 753 | 0.023 (1.00 / 0.054) | 3.3 × 10<sup>−16</sup> (288, 0.04%) | 1.000 (0.02%) | 1.000 (0.01%) | +86.57 (+57.85) / 36 | 8 → 5 / 4 | Multi-hit FP eliminated |
| 2411 (*USF1*) | 22 × 298 → 298 | 0.036 (1.00 / 0.93) | 1.5 × 10<sup>−5</sup> (436, 0.06%) | 1.000 (0.00%) | 1.000 (0.00%) | +129.35 (+4.66) / 61 | 5 → 3 / 25 | Indel/MH FP eliminated |
| 10048 (*VEPH1*) | 39 × 826 → 828 | 0.171 (1.00 / 0.34) | < 10<sup>−16</sup> (2.9, 21.0%) | 1.000 (0.00%) | 1.000 (0.00%) | +76.96 (−34.06<sup>§</sup>) / 59 | 44 → 40 / 0 | SRV/drift FP eliminated |
| 5220 (*NOP2*) | 38 × 586 → 586 | 0.097 (1.00 / 0.23) | 2.2 × 10<sup>−6</sup> (9.0, 0.50%) | 0.044 (0.04%) | 0.072 (0.03%) | +78.46 (+40.65) / 74 | 29 → 30 / 0 | Indel FP eliminated in -u |
| 10062 (*GSTK1*) | 36 × 199 → 199 | 0.409 (0.003 / 7 × 10<sup>−4</sup>) | 5.2 × 10<sup>−4</sup> (1.8, 26.5%) | 1.4 × 10<sup>−5</sup> (0.00%) | 0.0055 (0.00%) | +23.87 (+4.16) / 13 | 14 → 13 / 0 | 400× indel inflation attenuated |
| *Avian immune and cytokine receptors, Stage 5 indel-hotspot and clade-regime discoveries, and full 11,270-orthogroup genome screen* |  |  |  |  |  |  |  |  |
| 18109 (*THPO*) | 38 × 190 → 192 | 0.242 (5 × 10<sup>−4</sup> / 2 × 10<sup>−8</sup>) | 1.3 × 10<sup>−12</sup> (173, 0.40%) | 1.000 (0.06%) | 0.0126 (0.09%) | +87.35 (+18.44) / 25 | 11 → 10 / 0 | Rescued by -u alignment |
| 1224 (*MARCO*) | 36 × 466 → 466 | 0.656 (< 10<sup>−16</sup> / < 10<sup>−16</sup>) | < 10<sup>−16</sup> (9.5, 34.2%) | 3.2 × 10<sup>−24</sup> (0.32%) | 1.3 × 10<sup>−23</sup> (0.16%) | +207.04 (+55.66) / 102 | 62 → 65 / 0 | True positive; *w*<sub>err</sub> halved |
| 16962 (*MX1*) | 38 × 671 → 672 | 0.599 (< 10<sup>−16</sup> / < 10<sup>−16</sup>) | < 10<sup>−16</sup> (6.0, 5.9%) | 2.8 × 10<sup>−42</sup> (0.00%) | 1.1 × 10<sup>−41</sup> (0.00%) | +14.65 (−39.15<sup>§</sup>) / 72 | 67 → 67 / 0 | Strong true positive (*w*<sub>err</sub> = 0%) |
| 18627 (*TLR4*) | 39 × 843 → 843 | 0.419 (< 10<sup>−16</sup> / < 10<sup>−16</sup>) | 2.2 × 10<sup>−7</sup> (2.5, 13.6%) | 7.1 × 10<sup>−12</sup> (0.00%) | 1.6 × 10<sup>−11</sup> (0.00%) | +12.48 (+7.52) / 66 | 66 → 66 / 0 | Strong true positive (*w*<sub>err</sub> = 0%) |
| 29335 (*AHSG*) | 38 × 345 → 351 | 0.453 (< 10<sup>−16</sup> / < 10<sup>−16</sup>) | < 10<sup>−16</sup> (6.2, 25.0%) | 2.7 × 10<sup>−12</sup> (0.09%) | 1.6 × 10<sup>−11</sup> (0.04%) | +36.00<sup>‡</sup> (+8.55) / 23 | 25 → 25 / 0 | Insertion isolated; core restored |
| 34964 (*PIGR*) | 43 × 547 → 615 | 0.472 (< 10<sup>−16</sup> / < 10<sup>−16</sup>) | < 10<sup>−16</sup> (4.9, 14.1%) | 4.3 × 10<sup>−20</sup> (0.24%) | 1.2 × 10<sup>−24</sup> (0.27%) | 78 indels (74/93 pos.) | 78 → 81 / 88 (69) | Passerine *K̂* = 2.0 (ΔBIC = +21.0) |
| 7222 (*TFRC*) | 39 × 776 → 778 | 0.296 (< 10<sup>−16</sup> / < 10<sup>−16</sup>) | < 10<sup>−16</sup> (5.1, 4.9%) | 3.7 × 10<sup>−38</sup> (0.00%) | 1.4 × 10<sup>−39</sup> (0.00%) | +10.27 (51 indels) | 71 → 72 / 63 (48) | Galliform *K̂* = 3.0 (ΔBIC = +24.2) |
| 6798 (*ZNFX1*) | 38 × 1,796 → 1,798 | 0.251 (< 10<sup>−16</sup> / < 10<sup>−16</sup>) | < 10<sup>−16</sup> (3.4, 3.8%) | 2.9 × 10<sup>−20</sup> (0.00%) | 4.8 × 10<sup>−21</sup> (0.00%) | +52.24 (94 indels) | 115 → 117 / 293 (155) | Passerine *K̂* = 4.5 (ΔBIC = +112.0) |
| Random 50 Cohort | 50 orthogroups | 14/50 M2a<sup>+</sup> (28%) | 30/50 (60%, 19 M2a<sup>−</sup>) | 11/50 (22%, 2 M2a<sup>−</sup>) | 10/50 (20%, 2 M2a<sup>−</sup>) | +36.04 mean / 48.2 | 684 → 670 / 118 | 17/19 M2a<sup>−</sup> FPs purged |
| Full Avian Genome | 11,190 adm. (28 Gate 0, 52 T/O) | 7,127 conserved (ω<sub>M0</sub> < 0.2) | 55.4% (46.4% on cons.) | 40.0% on T/O (*q*) | 17.2% (*p*) / 12.1% (*q*) | 399,735 indels (1.94×) | 160,225 / 32,582 | 13× cons. (28× ultracons.) FP drop |

<sup>†</sup>Nominal *p* = 0.017 in -u is non-significant after genome-wide Benjamini–Hochberg FDR control (*q* = 0.097 > 0.05). <sup>‡</sup>Local 16-codon insertion-window Δ ln *L* = +36.00 nats (+8.24 nats MG94, +8.55 nats joint gene-wide).
<sup>§</sup>In *VEPH1* and *MX1*, GUIDANCE masked indel boundaries as NNN codons in the input alignment (incurring zero affine gap penalties in -a while being resolved as explicit branch indels in -u).

#### Unfiltered Functional Enrichment and Housekeeping Specificity

Across all 11, 190 admitted orthogroups, combining Stage 3–4 model unification (GC_3_ drift mixture, Empirical Bayes SRV, and the BUSTED-E error sink) with Stage 1 phylogeny-aware realignment (WALRUS -u) reduces the genome-wide positive selection rate 4.6-fold from 55.4% (6, 198 genes under published stationary single-hit BUSTED) to 12.1% (1, 353 genes) at Benjamini–Hochberg *q ≤* 0.05. At nominal *p ≤* 0.05 and stringent *q ≤* 0.01, WALRUS (-u) flags 17.2% (1, 926 genes) and 9.2% (1, 031 genes), respectively. Notably, this reduction is concentrated in conserved proteins. False-positive calls on the 7, 127 conserved housekeeping and developmental orthogroups (*ω*_M0_ *<* 0.20*, p*_M2a_ *>* 0.50) drop 13.0-fold from 46.4% (3, 309 genes) to 3.56% (254 genes at *q ≤* 0.05; 6.58% at nominal *p ≤* 0.05). On the 3, 022 ultraconserved proteins (*ω*_M0_ *<* 0.05*, p*_M2a_ *>* 0.50), false positives drop 27.8-fold, from 41.4% (1, 250 genes) to 1.49% (45 genes; Figure 7C). Paired aligned (-a) versus unaligned (-u) runs on the 68-gene benchmark cohort (Table 6) decompose these two contributions. Applying Stages 3–4 directly to the published PRANK -codon alignments (WALRUS -a) eliminates 17*/*19 (89.5%) of nominal PAML M2a*^−^*false positives in the random 50-gene cohort (60% *→* 22%) while recovering 9*/*14 (64%) of PAML M2a^+^ genes. It also resolves multi-hit and SRV inflation across conserved focal genes (*UROD*, *COPB1*, *TLK2*, *USF1*, *VEPH1*; all *p ≥* 0.094, with *q* = 0.097 *>* 0.05 on *UROD* in -u). Stage 1 realignment (-u) then purges remaining gap-boundary inflation, eliminating the borderline call on nucleolar rRNA methyl-transferase *NOP2* (HOG 5220, *p* = 0.044 *→* 0.072, Δ ln *L*_MG94_ = +78.46, Δ*l*_joint_ = +40.65 nats), attenuating 400-fold gap inflation on *GSTK1* (*p* = 1.4 *×* 10*^−^*^5^ *→* 0.0055, concordant with PAML M2a *p* = 0.003), and removing the Codon 245 artifact in HOG 13310. Simultaneously, Stage 1 realignment rescues true positive selection after error-sink quarantine of misaligned indels in *THPO* (HOG 18109, *p* = 1.00 *→* 0.0126) and strengthens signal in mucosal receptor *PIGR* (*p* = 4.3 *×* 10*^−^*^20^ *→* 1.2 *×* 10*^−^*^24^). Because standalone BUSTED flagged nearly half of all conserved avian genes, Shultz and Sackton (2019) had to intersect five separate tools (BUSTED plus four PAML site models) to recover pathway-level signal. By contrast, a single unfiltered WALRUS (-u) run slashes positive selection calls at *q ≤* 0.05 across core housekeeping machinery, dropping to 1.2% (1*/*80, vs. 37.5% in published BUSTED) in the spliceosome, 3.0% (3*/*99, vs. 36.4%) in ribosomal proteins, and 3.2% (1*/*31, vs. 48.4%) in the proteasome. Comparable reductions occur across the citrate/TCA cycle (4.5% [1*/*22] vs. 45.5%), aminoacyl-tRNA biosynthesis (6.5% [2*/*31] vs. 51.6%), and ribosome biogenesis (8.8% [5*/*57] vs. 56.1%). Without any multi-tool intersection, WALRUS (-u) increases the KEGG enrichment odds ratio between host-defense pathways (360 genes, 23.9% selected) and core housekeeping machinery (568 genes, 4.8% selected) from OR = 2.36 to OR = 6.29 (Fisher *p* = 1.2 *×* 10*^−^*^17^). That specificity directly surfaces strong enrichment in Fanconi anemia DNA repair (35.9%, OR = 4.76*, p* = 2.8 *×* 10*^−^*^5^), Influenza A antiviral response (34.2%, OR = 4.54*, p* = 3.0 *×* 10*^−^*^8^), necroptosis (31.0%, OR = 3.88*, p* = 2.5 *×* 10*^−^*^6^), Toll-like receptor signaling (29.5%, OR = 3.55*, p* = 5.1 *×* 10*^−^*^4^), and cytokine–receptor interaction (28.3%, OR = 3.50*, p* = 5.0 *×* 10*^−^*^8^; Figure 7C).

#### Covariate Determinants of Alignment Repair, Sequence Quarantine, and Error-Sink Absorption

We next compared the 11, 183 completed orthogroups that have complete paired published (PRANK -codon + GUIDANCE) alignments and PAML/BUSTED metadata in Shultz and Sackton (2019) against evolutionary covariates. (Seven of the 11, 190 completed orthogroups, namely HOG 107, 611, 2077, 2172, 3248, 5500, and one alignment with NA length, lacked published summary metadata.) In the published archive, GUIDANCE masked 148, 967 individual codons as NNN across 95.9% (10, 729*/*11, 183) of alignments while leaving 389, 144 gap codons in place. Unaligned WALRUS (-u) instead modifies gap placement in 96.4% (10, 775) of orthogroups and expands alignment length (Δ*C >* 0) in 25.2% (2, 815) to un-crush over-packed insertions. In place of ad hoc NNN masking, WALRUS applies three targeted safeguards. First, Stage 0 quarantines 583 frame-broken or non-homologous sequences across 326 orthogroups (2.92%). Second, Stage 1 POA column-uncertainty marginalization eliminates 2, 958 gap-boundary false-positive site calls across 1, 742 genes (15.6%) and rescues 293 true-positive sites. Third, Stage 4 BUSTED-E error-sink absorption activates (*w*_err_ *>* 0.01%) in 2, 295 genes (20.5%, mean *w*_err_ = 0.011%), with 0% hitting the 1.5% boundary cap after the *W*_sink_ scale fix. Across coding length quintiles (*Q*_1_ *≤* 204 codons to *Q*_5_ *>* 693 codons), the per-gene probability of alignment expansion (12.0% *→* 43.8%*, r_s_* = +0.264) and POA false-positive elimination (3.6% *→* 34.0%*, r_s_*= +0.328) increases steeply because longer multi-domain proteins accumulate more total branch indels (16.5 *→* 67.5 indels per gene). This scaling allows WALRUS to moderate the severe gene-length inflation of published BUSTED (32.6% in *Q*_1_ *→* 79.2% in *Q*_5_, *r_s_* = +0.331, vs. 5.6% *→* 21.7% in WALRUS, *r_s_* = +0.168). Normalized per codon, however, short proteins (*Q*_1_) exhibit higher indel density (9.56 vs. 6.51 indels per 100 codons in *Q*_5_), higher Stage 0 sequence quarantine (4.5% vs. 2.4%), and higher mean *w*_err_ (0.0142% vs. 0.0093%). In multivariate rank regressions controlling for all covariates simultaneously, evolutionary tree length (*T*_M0_) and indel density are the primary per-codon drivers of sequence quarantine (1.6% *→* 6.6% across *T*_M0_ quartiles), POA boundary filtering (0.10 *→* 0.90 sites eliminated per 1, 000 codons), and error-sink weight (*w*_err_: 0.0044% *→* 0.0235%). Finally, taxon copy count (*N*) exhibits a U-shaped dependence centered on clean single-copy 39–40-taxon orthogroups (1.5% quarantine, mean *w*_err_ = 0.0062%). Quarantine rises to 5.0% (*w*_err_ = 0.0216%) in patchy 20–34-taxon assemblies and spikes nearly 10-fold to 14.2% (96*/*678 orthogroups, *w*_err_ = 0.0299%) in multi-copy orthogroups (*N* = 41–51) enriched for mis-assigned paralogs and fragmented tandem duplications.

#### Genome-Wide Prevalence of Directional GC_3_ Drift, CpG Hypermutability, and Synonymous Constraint

Synthesizing Stage 3 and Stage 5 diagnostics across all 11, 190 orthogroups quantifies how non-adaptive mutational processes shape avian coding evolution (Figure 7D). First, 52.1% (5, 832*/*11, 190) of avian genes reject nucleotide stationarity and activate Stage 3’s phylogenetic directional GC_3_ drift mixture, consistent with widespread recombination-driven GC-biased gene conversion (gBGC) across avian microchromosomes. Under published stationary BUSTED, these GC_3_-drifting genes were called positively selected significantly more often than stationary genes (59.2% vs. 51.3%, OR = 1.38, Fisher *p* = 3.6 *×* 10*^−^*^17^), which shows that unmodeled compositional drift systematically inflated previous avian selection scans. Accounting for directional GC_3_ flux in WALRUS (-u) reverses this relationship (10.75% [627*/*5, 832] vs. 13.55% [726*/*5, 358] at *q ≤* 0.05, OR = 0.77*, p* = 5.9 *×* 10*^−^*^6^), consistent with strong purifying constraint on compact microchromosomal housekeeping loci. Second, 50.5% (5, 647) of avian orthogroups contain positively selected sites within hypermutable CpG dinucleotides (17.0% [1, 907 genes] significantly enriched at *p ≤* 0.05). Third, 57.3% (6, 415) of avian genes harbor 13, 383 synonymous constraint islands (*z ≥* 3.0*, p ≤* 0.01) marking overlapping splicing enhancers or conserved mRNA structures.

#### Indel–Selection Coupling, Lineage-Specific Relaxation, and Autonomous Clade Discovery

Across the 5.70 *×* 10^6^ aligned avian codons, Stage 1 reconstructs 399, 735 branch indel events and Stage 5 localizes 71.9% (99, 646*/*138, 528) of Simes-significant positively selected sites to indel-proximal loop windows (37.1% of codons; 1.94-fold enrichment), while POA marginalization purges 2, 958 gap-boundary false positives and rescues 293 true-positive sites (Figure 7D). Along the phylogeny, 9.7% of genes (1, 085 orthogroups, 1, 163 branches) exhibit lineage-specific loss-of-constraint clustering in ecological specialists such as budgerigar (*Melopsittacus undulatus*), downy woodpecker (*Picoides pubescens*), Anna’s hummingbird (*Calypte anna*), and flightless/semi-terrestrial tinamous. Simultaneously, Stage 5’s BIC clade scan (ΔBIC *≥* 10.0) autonomously detects macro-evolutionary regime shifts in 631 avian genes (5.6%) spanning 32, 582 Contrast-FEL sites, uncovering clade-specific adaptation and domain localization (Figure 7E and Figure 8C–D). In the antiviral RNA helicase *ZNFX1* (HOG 6798, 1, 798 codons), positive selection is restricted to a 1.24% site fraction (*ω*_3_ = 7.0, gene-wide BUSTED-E *p* = 4.8 *×* 10*^−^*^21^) embedded within a conserved 1,500-codon helicase scaffold. Within that protein, Stage 5 discovers a 4.5-fold intensification in Passeriformes (Node30, ΔBIC = +112.0, nominal *p* = 9.4 *×* 10*^−^*^29^, tree-wide Holm *p*_adj_ = 6.9 *×* 10*^−^*^27^) across 293 clade-regime Contrast-FEL sites (155 at site-atlas *p ≤* 0.05) in its viral-binding disordered domains (*ω*^-^ = 1.07–1.50). Parallel domain–clade coupling is resolved in the mucosal IgA receptor *PIGR* (HOG 34964, BUSTED-E *p* = 1.2 *×* 10*^−^*^24^, Passerine *K*^ = 2.00, ΔBIC = +21.0), where 79.6% (74*/*93) of positively selected sites and 83.5% (147*/*176) of Simes omnibus *q ≤* 0.10 sites lie in indel loops. A similar architecture appears in the viral-entry transferrin receptor *TFRC* (HOG 7222, BUSTED-E *p* = 1.4 *×* 10*^−^*^39^, apical-loop Codon 220*→*Glu convergence, 28*/*72 positively selected sites in indel loops; Galliform *K*^ = 3.00, ΔBIC = +24.2; Table 6).

## 6 Discussion: Practical Guidelines for Routine Comparative Genomics

Coupling sequence quality control, phylogeny-aware codon alignment, directional base-composition drift correction, error-sink quarantine, and multi-scale selection testing inside a single engine replaces a brittle chain of separate programs with one reproducible command (hyphy3 -u sequences.fa or hyphy3 -a alignment.nex). Our worked examples, synthetic misspecification benchmarks, and vertebrate comparative analyses point to three practical guidelines for routine evolutionary genomics. The first guideline is that coding sequence alignment should respect both the phylogenetic tree and column occupancy whenever insertions and deletions are present. Because standard progressive aligners charge an independent gap penalty for every species in a gapped clade, their additive scoring objective creates a strong incentive to pack unrelated insertions from different lineages into the same column or push terminal deletions to the edge of the protein, as observed at the C-terminus of abalone sperm lysin (Figure 2B). Evaluating branch transitions under a two-stratum codon and amino-acid transducer (Eq. 1), penalizing matches to low-occupancy insertion columns, and maximizing a joint substitution-plus-indel likelihood (Eq. 2) eliminate this structural bias. Across 800 simulated indel alignments, this joint formulation recovers 94.6% of ground-truth columns and reduces downstream gene-wide false positives from 44.7%– 58.1% to 4.4% under standard BUSTED and from 21.1%– 21.4% to 1.4% under BUSTED-E (Figure 6A).

A second operational lesson is that gene-wide and site-level selection tests respond very differently to alignment boundary noise and instantaneous multi-nucleotide mutations, although both benefit from a shared sufficiency tensor. Gene-wide random-effects tests (BUSTED) pool likelihood evidence across every branch and codon into a single high-*ω* tail category. Consequently, a single misaligned gap boundary in abalone lysin or third-position GC_3_ turnover coupled to two-nucleotide aliphatic codon switches (GTG [Val] *↔* GCA/GCG [Ala] / ACG [Thr]) in mammalian *RBP3* (Figure 3A) is sufficient to trigger a spurious gene-wide rejection. Incorporating synonymous rate variation and an alignment-error sink (BUSTED-E, *ω*_err_ = 100) while marginalizing over alternative Partial Order Alignment columns as a matter of routine protects gene-wide inference without sacrificing power on genuinely adaptive proteins. By contrast, fixed-effects site tests (FEL and dFEL) fit substitution rates at each column independently and treat gaps as missing data, which renders them naturally resilient to indel noise once third-position GC drift is modeled (Eq. 3).

A third practical guideline is that genome-scale screens should separate background mutational drift and clade-specific evolutionary shifts from whole-gene averages while respecting phylogenetic depth and topological coherence. In large comparative collections such as the avian proteome, GC-biased gene conversion on microchromosomes inflates stationary BUSTED calls to 59.2% unless directional GC_3_ mixtures are modeled (10.8%; Figure 7D). Meanwhile, lineage-restricted adaptations inside otherwise conserved multi-domain proteins (such as passerine *ZNFX1*) are resolved across specific clades and domains by autonomous BIC clade scanning (with tree-wide Holm multiplicity control) and BIC-penalized 1D change-point domain segmentation. Even so, practitioners should note three boundary conditions when interpreting unified runs. First, on very shallow trees (*T <* 0.2 substitutions per codon) or small taxon sets (*N <* 8), site-specific synonymous rates and multi-regime directional mixtures are weakly identified. Second, under extreme Zipfian indel or multi-hit densities (*λ*_indel_ *≥* 0.02, *δ ≥* 0.10), clustered artifacts can exceed the default *w*_err_ *≤* 2.5% error-sink cap. Third, on ultra-long multi-domain proteins (*L >* 3,000 codons, such as titin or dynein homologs) or genes undergoing extensive intragenic recombination, anchor-constrained banding or pre-partitioned aligned (-a) execution is recommended to avoid quadratic dynamic programming overhead. Precomputing the branch-by-site sufficiency tensor once (Eq. 4) and polishing only active-set sites and BIC-validated clades reduces total runtime by 6.6-fold to 35.8-fold relative to running six standalone tools on individual genes (Figure 6B) and cuts total compute by 16-fold to 27-fold across genome-scale orthogroup collections.

## Data and Software Availability

HyPhy 3 (hyphy3, hyphy3-walrus, hyphy3-characterize, hyphy3-joint-align, hyphy3-busted, hyphy3-fel, hyphy3-dfel, hyphy3-meme, hyphy3-absrel, hyphy3-relax, hyphy3-contrast-fel, and hyphy3-prime) is implemented in C++20 and is freely available under an open-source MIT license at https://github.com/veg/hyphy-3 (reference commit afacb3e), with a zero-install WebAssembly browser application at https://veg.github.io/hyphy-3/. Complete mathematical derivations and algorithmic specifications for Stages 0–5 are publicly archived on Zenodo at https://doi.org/10.5281/zenodo.24021779 alongside the complete replication package (hyphy3_walrus_zenodo_replication_package. licensed under CC-BY-4.0). That archive contains all worked-example sequences and JSON/HTML dossiers, the 10,020-alignment simulation suites and per-replicate outputs, the 11,190-orthogroup avian exome reanalysis tables and paired -a/-u benchmarks, figure-generation scripts, and an automated master reproduction script (reproduce_all.sh). All empirical analyses were conducted retrospectively *in silico* on publicly archived nucleotide accessions from NCBI GenBank/RefSeq, VGP/TOGA2, and published benchmark repositories.

## Supporting information

Supplementary Methods and Technical Reference

## Acknowledgments, Funding, and Disclosure of AI-Assisted Technologies

This work was supported in part by the National Institutes of Health (grants AI183870, GM151683, and HG009299) and the National Science Foundation (grant 2419522). The authors disclose that agentic AI coding and writing assistants (Google Antigravity powered by Gemini and Anthropic Claude) were used extensively during this project for rapid algorithmic prototyping, C++20 and WebAssembly software implementation, automated testing and parity verification, simulation and empirical benchmark execution, figure generation, and manuscript drafting and editing. In particular, agentic workflows enabled systematic head-to-head benchmarking of competing algorithmic formulations against adversarial misspecification simulations prior to production deployment. Those comparisons included evaluating our two-stratum MG94 + WAG transducer with column-occupancy priors (Eq. 1) against standard single-stratum codon and amino-acid aligners on indel-rich loops, and testing graph-Laplacian Ohmic Wheatstone bridges against contiguous-breakpoint partitioning on concerted repeat conversion. The winning mathematical formulations were then translated into vectorized C++20 code verified by automated continuous-integration parity tests. The authors conceived and directed the study, reviewed and validated all mathematical formulations, software implementations, and empirical findings, and assume full responsibility for the content of the manuscript and accompanying software.

