## Supplementary Methods and Technical Reference for "All Together Now: Joint Codon Alignment, Ohmic Networks, and Error-Controlled Selection Inference in HyPhy 3"

### Supplementary Methods and Technical Reference: Mathematical and Algorithmic Architecture of WALRUS in HyPhy 3

---

Overview. This companion Supplementary Methods and Technical Reference presents the complete, self-contained mathematical formulation, dynamic programming recurrences, circuit-theoretic topology solvers, hidden Markov model smoothing recurrences, and Inside–Outside sufficiency tensor derivations for Stages 0 through 5 of `WALRUS` (*Whole-gene Alignment, Lineage, Recombination, and Unified Selection*, invoked as `hyphy3-walrus` or `hyphy3 characterize`), alongside supplementary covariate stratification and codon-level alignment forensics for the 11,190-orthogroup avian exome reanalysis. It accompanies the main manuscript, *All Together Now: Joint Codon Alignment, Ohmic Networks, and Error-Controlled Selection Inference in HyPhy 3*. All algorithms are implemented in C++20 at `https://github.com/veg/hyphy-3` (commit `afacb3e`), and the complete replication package is archived on Zenodo at `https://doi.org/10.5281/zenodo.24021779`.

---

### 1 Complete Mathematical and Algorithmic Specification of WALRUS

Conventional evolutionary analyses of protein-coding DNA treat sequence curation, multiple sequence alignment, tree estimation, recombination screening, and selection testing as independent sequential steps. Standard pipelines pass a single point-estimate alignment and tree down a chain of disconnected programs, even though judging whether two codons are homologous across an indel boundary—or whether a cluster of nonsynonymous changes reflects positive selection rather than recombination or compositional drift—depends on the underlying phylogeny and codon substitution process. `WALRUS` (*Whole-gene Alignment, Lineage, Recombination, and Unified Selection*, invoked as `hyphy3-walrus` or `hyphy3 characterize`) replaces this multi-tool chain with a unified six-stage inference workflow (Stages 0–5) that accepts either raw unaligned coding sequences (`-u`) or an existing alignment (`-a`).

When starting from unaligned sequences (`-u`), Stage 0 screens inputs for reading-frame integrity, strand orientation, internal stop codons, terminal non-homologous overhangs, and sequence homology, shifting frame-shifted or reverse-complemented entries into the canonical coding frame while excluding pseudogenes and non-homologous outliers. Stage 1 co-estimates an in-frame codon alignment and initial phylogeny by combining an alignment-free screened Ohmic conductance guide scaffold with a continuous-time two-stratum Markov codon transducer (MG94 $\times$ REV augmented with empirical core/loop Grantham–Miyata and multi-hit exchangeabilities) whose match and gap scores depend on branch lengths and fractional clade occupancy. Local insertion–deletion windows are then refined via leave-one-taxon-out Felsenstein Inside–Outside cavities and a two-state Pair-HMM, retaining both the maximum a posteriori (MAP) alignment and a Partial Order Alignment (POA) graph that records column-level homology uncertainty ($PP_c$). Stage 2 breaks guide-tree indel lock-in via coupled Ohmic Subtree Prune and Regraft (SPR) search with local indel realignment, and screens for intragenic recombination via Kadane's $O(C)$ maximum-subarray breakpoint routing gated by 2,000 site-order permutations and BIC. Stage 3 audits third-codon-position ($GC_3$) compositional drift via phylogenetic generalized least squares (GLS) ancestral reconstruction and Inside–Outside $W \leftrightarrow S$ substitution flux, automatically rooting the tree along the branch of maximal directional divergence and activating a three-regime Softmax directional kernel (`dFEL`) when stationarity is rejected. Stage 4 pre-computes a shared bank of Inside (subtree) and Outside (complement-tree) conditional likelihood vectors across a 16-point $\omega$ spectral grid, averaging over up to $K = 3$ POA column realizations where $PP_c < 0.99$, projecting into the $(k, g)$ eigenbasis via Hadamard products for exact site-rate ($\tilde{\alpha}_c$) and segment-routed branch-length ($t_{b,c}$) scaling, and attenuating multi-codon frameshift/overhang tracts via an Erlang-3 minimum-duration Baum–Welch error-sink HMM. Finally, Stage 5 executes a two-stage screen-and-polish across all biological scales from this single cached sufficiency tensor $W[c][b][g]$: gene-wide episodic diversification with an error sink and boundary-consistency guards (`BUSTED-E`), BIC-penalized 1D change-point protein domain segmentation and Synonymous Constraint Island (SCI) detection, autonomous clade-level selection intensity (RELAX) and regime contrasts (Contrast-FEL) with tree-wide Holm multiplicity control, branch-level adaptive bursts (aBSREL) versus core loss-of-constraint (LoC), and codon-level pervasive (FEL/dFEL), episodic (MEME), and biochemical property (PRIME) selection synthesized via Simes omnibus testing and Benjamini–Hochberg false-discovery control.

#### 1.1 Stage 0: Four-Gate Input Filtering, Phase Repair, and Homology Admission Control

Public sequence repositories and automated orthology pipelines frequently contain annotation defects that violate the assumptions of codon substitution models: amino-acid FASTA files mislabeled as nucleotide coding sequences (CDS), antisense transcripts deposited on the reverse-complement strand, 5′-truncated exons out of reading-frame phase, unspliced introns or pseudogenes harboring premature stop codons, chimeric terminal extensions, and non-homologous paralogs. Admitting even a single out-of-frame or non-homologous sequence into progressive alignment forces dozens of spurious nonsynonymous substitutions and frame-shifting gaps across the phylogeny. WALRUS applies a four-gate admission filter before alignment begins (when invoked with -u) and an alignment-guided terminal overhang trimmer when invoked on pre-aligned sequences (-a).

**Gate 1 (Lexical and coverage verification).** Gate 1 verifies that the input alphabet is nucleotide rather than amino-acid text (rejecting files in which amino-acid-specific letters E, F, I, L, P, Q exceed 8% of alphabetic characters) and excludes empty sequences, fragments shorter than 15 sense codons, or heavily masked entries containing more than 20% ambiguous (N or ?) bases.

**Gate 2 (Six-frame phase and strand resolution).** Gate 2 resolves reading-frame phase and strand orientation across all six reading frames ($+0, +1, +2$ on the forward and reverse-complement strands) under the active genetic code. In Pass 1, sequences that have zero internal stop codons in frame $+0$ and begin with an ATG start codon (or all stop-free frame-$+0$ sequences if none begin with ATG) populate a reference hash table of observed amino-acid 3-mers. In Pass 2, any sequence that contains internal stop codons in frame $+0$, or that lacks an ATG start and shares fewer than 15 amino-acid 3-mers with the cohort reference table, is translated across all five alternative frames (catching GC-rich antisense sequences that happen to lack stop codons on the reverse strand). A candidate is shifted or reverse-complemented into an alternative frame if and only if that frame is free of internal stop codons and strictly increases shared cohort 3-mer hits; sequences carrying internal stop codons across all six frames are excluded as pseudogenes or intronic contaminants. If more than 25% of input sequences fail reading-frame validation, execution terminates with a diagnostic report.

**Gate 3 (Piecewise-affine capped semi-global homology DP and decoy $Z$-score calibration).** Gate 3 removes extreme length outliers ($< 0.30\times$ or $> 2.80\times$ the cohort median codon length) and tests pairwise sequence homology across the $M$ surviving candidates. For each pair $(S_i, S_j)$, we order the sequences by length ($L_{\text{eff}} = \min(L_i, L_j)$, $L_{\max} = \max(L_i, L_j)$, $\Delta L = L_{\max} - L_{\text{eff}}$) and align them via semi-global overlap banded codon dynamic programming with band slack $B_0 = 24$ codons. Leading $5'$ overhangs on the longer sequence up to $\Delta L + B_0$ codons and trailing $3'$ overhangs within $[\max(1, L_{\text{eff}} - B_0), L_{\max}]$ incur zero terminal gap penalty, requiring $O((B_0 + \Delta L)L_{\text{eff}})$ time and $O(L_{\max})$ memory. Codon pairs $(c_1, c_2)$ and internal gaps of length $g \geq 1$ are scored via piecewise-affine capped recurrences:

$$s(c_1, c_2) = \begin{cases} 1.00, & c_1 = c_2, \\ 0.90, & c_1 \neq c_2,\ A(c_1) = A(c_2), \\ 0.30, & A(c_1) \neq A(c_2),\ d_{\text{nuc}}(c_1, c_2) = 1, \\ 0.00, & d_{\text{nuc}}(c_1, c_2) \geq 2,\ A(c_1) \neq A(c_2), \end{cases} \qquad \Gamma(g) = -\min(\gamma_{\text{open}} + \gamma_{\text{ext}}\, g,\ \gamma_{\text{cap}}), \quad \text{(S1)}$$

with gap-opening penalty $\gamma_{\text{open}} = 1.10$, extension penalty $\gamma_{\text{ext}} = 0.30$, and exon-skip cap $\gamma_{\text{cap}} = 3.80$ (reached at $g \geq 9$ codons). Capping internal gaps at $\gamma_{\text{cap}}$ bridges across large internal domain insertions or alternatively spliced exons ($A_1BA_2$ versus $A_1A_2$) at a fixed cost equivalent to four identical codons, while the steep affine penalty on short gaps ($1 \leq g \leq 8$) suppresses opportunistic gap-hunting in unrelated sequences. At the optimal semi-global terminal column $c^* \in [\max(1, L_{\text{eff}} - B_0), L_{\max}]$, both the alignment score and exact amino-acid match count are normalized by $L_{\text{eff}}$, yielding the per-codon overlap score $\tilde{S}_{ij} = S_{ij}(L_{\text{eff}}, c^*)/L_{\text{eff}}$ and fractional overlap amino-acid identity $\text{ID}_{ij} = \text{matches}_{ij}(L_{\text{eff}}, c^*)/L_{\text{eff}}$.

To standardize $(\text{ID}_{ij}, \tilde{S}_{ij})$ against the pair's exact codon composition and overhang geometry, we generate $K = 16$ deterministic Fisher–Yates codon permutations $(S_i^{(k)}, S_j^{(k)})_{k=1}^{K}$ and align each decoy pair with the same semi-global dynamic program to obtain empirical null means $(\hat{\mu}_{\text{ID}}, \hat{\mu}_{\text{DP}})$ and sample variances $(s^2_{\text{ID}}, s^2_{\text{DP}})$. Shrinking the sample variances toward the analytical within-alignment variance bounds $\sigma^2_{\text{ID,plug}} = \hat{\mu}_{\text{ID}}(1 - \hat{\mu}_{\text{ID}})/L_{\text{eff}}$ and $\sigma^2_{\text{DP,plug}} = 0.35^2/L_{\text{eff}}$ with prior weight $\nu_0 = 16$ yields the regularized homology $Z$-score:

$$\hat{\sigma}^2_{\text{ID}} = \left(1 + \frac{1}{K}\right) \max\left(\sigma^2_{\text{ID,plug}},\ \frac{\nu_0\, \sigma^2_{\text{ID,plug}} + (K-1)s^2_{\text{ID}}}{\nu_0 + K - 1}\right), \qquad Z_{ij} = 0.65\, \frac{\text{ID}_{ij} - \hat{\mu}_{\text{ID}}}{\hat{\sigma}_{\text{ID}}} + 0.35\, \frac{\tilde{S}_{ij} - \hat{\mu}_{\text{DP}}}{\hat{\sigma}_{\text{DP}}}. \quad \text{(S2)}$$

Because each decoy averages over $L_{\text{eff}}$ codon steps ($16\, L_{\text{eff}}$ aligned steps in total), $K = 16$ bounds mean-estimation standard-error inflation to $\sqrt{1 + 1/16} = 1.031\times$ while the analytical floor $\sigma^2_{\text{ID,plug}}$ suppresses finite-$K$ Student-$t_{15}$ tail inflation at short sequence lengths. Taxa $i$ and $j$ are linked in an undirected homology graph whenever $Z_{ij} \geq 3.5$ and $\text{ID}_{ij} \geq \max(0.18, \hat{\mu}_{\text{ID}} + 0.04)$, and the largest connected component is retained via breadth-first search (halting if $< 3$ sequences remain or if the cohort median $Z_{ij} < 4.5$; Table S1 and Fig. S1). In addition, before progressive alignment, terminal non-homologous overhangs (e.g., unannotated $5'$ leader peptides or $3'$ read-through extensions that fail local homology conservation against the cohort consensus) are trimmed, and redundant identical or near-identical splice isoforms from the same species header are collapsed when requested.

**Gate 4 (Post-alignment expansion and core-occupancy verification).** Following progressive alignment in Stage 1, Gate 4 verifies the assembled alignment length $C$ against the median input length $L_{\text{med}}$. Capping the expansion ratio at $E = C/L_{\text{med}} \leq 2.0$ bounds the overall matrix gap fraction below 50%, while requiring at least 25% of columns to maintain core occupancy ($o_c \geq 0.40$, guaranteeing $\geq 3$ non-gap leaves in small cohorts) rejects non-transitive domain chaining ($A$–$B$, $B$–$C$, $C$–$D$) and ensures a shared orthologous backbone.

##### 1.2 Stage 1: Ohmic Guide Scaffold, Progressive Codon Profile Alignment, and Inside–Outside Window Realignment

Stage 1 transforms the admitted unaligned coding sequences into a reading-frame-preserving multiple codon alignment, an initial maximum-likelihood phylogeny, a per-column homology confidence profile ($PP_c$), and a Partial Order Alignment (POA) graph in three phases.

**Table S1.** *Operating characteristics of Stage 0 Gate 3 homology admission control ($K = 16$ codon-shuffled decoys, $\nu_0 = 16$, threshold $Z_{ij} \geq 3.5$).* Null regimes evaluate false-positive edge rates across unrelated sequence pairs. Homologous regimes evaluate true-positive admission recall under terminal truncation (50% overhang) and internal cassette-exon indels ($A_1BA_2$ versus $A_1A_2$, $|A_1| = |A_2| = 80$ codons).

| Evaluation Regime | Sequence Geometry ($L_i$ vs. $L_j$) | Observed $\mathrm{ID}_{ij}$ | Null $\hat{\mu}_{\mathrm{ID}}$ | Mean $Z_{ij}$ (Range / SD) | Rate ($Z_{ij} \geq 3.5$) |
|---|---|---|---|---|---|
| *Null Specificity (Unrelated Sequence Pairs — False-Positive Rate)* | | | | | |
| Uniform codon usage | 100 vs. 100 codons | 0.198 | 0.198 | $-0.01$ (SD $= 0.59$, max $= +1.87$) | 0.0% |
| Uniform codon usage | 300 vs. 300 codons | 0.210 | 0.210 | $-0.02$ (SD $= 0.58$, max $= +1.67$) | 0.0% |
| Uniform codon usage | 1000 vs. 1000 codons | 0.211 | 0.211 | $+0.00$ (SD $= 0.56$, max $= +1.78$) | 0.0% |
| Extreme GC bias (80% GC) | 300 vs. 300 codons | 0.291 | 0.290 | $+0.03$ (SD $= 0.55$, max $= +1.35$) | 0.0% |
| Asymmetric length null | 160 vs. 320 codons ($\Delta L = 160$) | 0.210 | 0.209 | $+0.02$ (SD $= 0.57$, max $= +1.38$) | 0.0% |
| *Homologous Sensitivity: $5'/3'$-Truncated Exons ($L_i = 160$ vs. $L_j = 320$ codons — True-Positive Recall)* | | | | | |
| Moderate divergence (45% ID) | 50% terminal truncation ($\Delta L = 160$) | 0.495 | 0.208 | $+17.12$ (min $= +12.88$) | 100.0% |
| Twilight Zone (30% ID) | 50% terminal truncation ($\Delta L = 160$) | 0.352 | 0.208 | $+9.17$ (min $= +4.26$) | 100.0% |
| Deep divergence (25% ID) | 50% terminal truncation ($\Delta L = 160$) | 0.304 | 0.208 | $+5.93$ (min $= +1.82$) | 90.0% |
| *Homologous Sensitivity: Internal Exon / Domain Indels ($A_1BA_2$ vs. $A_1A_2$ — True-Positive Recall)* | | | | | |
| Internal exon $|B| = 80$ (45% ID) | 160 vs. 240 codons (internal skip) | 0.490 | 0.200 | $+9.56$ (min $= +6.84$) | 100.0% |
| Internal exon $|B| = 80$ (35% ID) | 160 vs. 240 codons (internal skip) | 0.398 | 0.200 | $+6.57$ (min $= +4.12$) | 100.0% |
| Internal exon $|B| = 160$ (35% ID) | 160 vs. 320 codons (internal skip) | 0.392 | 0.208 | $+6.11$ (min $= +3.79$) | 100.0% |
| Internal exon $|B| = 300$ (35% ID) | 160 vs. 460 codons (internal skip) | 0.385 | 0.223 | $+5.47$ (min $= +3.41$) | 96.7% |

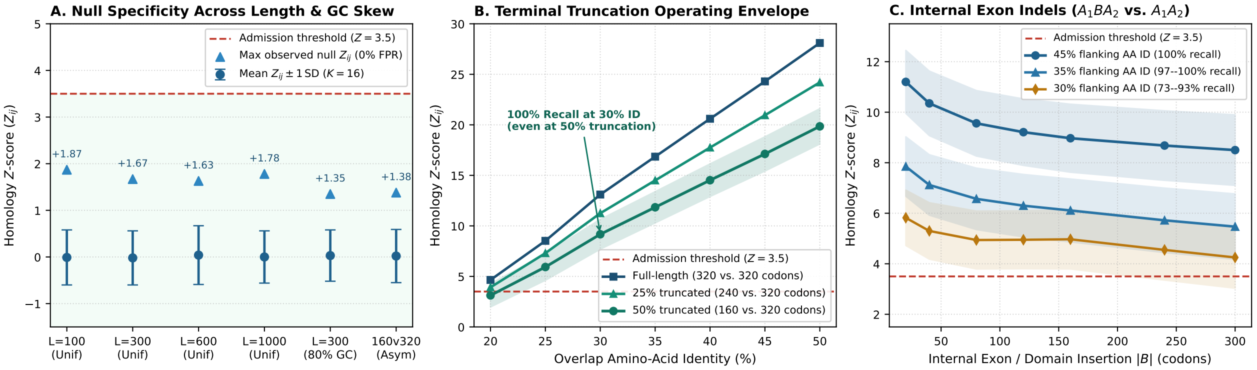

**Figure S1.** *Empirical operating characteristics of Stage 0 Gate 3 homology admission control ($K = 16$ codon-shuffled decoys, $\nu_0 = 16$, admission threshold $Z_{ij} \geq 3.5$).* Panel A: null specificity across sequence lengths ($L \in [100, 1000]$ codons), 80% GC skew, and asymmetric lengths (160 vs. 320 codons), yielding zero false positives ($\max Z_{ij} \leq 1.87$). Panel B: homology $Z_{ij}$ operating envelope across overlap amino-acid identity (20%–50%) for full-length, 25%-truncated, and 50%-truncated coding sequences. Panel C: homology $Z_{ij}$ across internal exon insertion lengths $|B| \in [20, 300]$ codons ($A_1BA_2$ vs. $A_1A_2$) under piecewise-affine capped gap scoring $\Gamma(g) = -\min(1.10 + 0.30\,g, 3.80)$.

**Phase 1: Multi-track alignment-free distances, screened Ohmic effective resistance, and rate-corrected guide tree $T_0$.** Along the optimal semi-global dynamic programming path between each sequence pair $(i, j)$ from Eq. (S1), we accumulate three evolutionary distance channels normalized by $L_{\mathrm{eff}} = \min(L_i, L_j)$: a synonymous codon track $D_{ij}^{(\mathrm{syn})}$ (resolving shallow splits), a nonsynonymous amino-acid track $D_{ij}^{(\mathrm{ns})}$ (unsaturated across deep splits), and a Tamura–Nei nucleotide distance $D_{ij}^{(\mathrm{TN93\text{-}DP})}$ over DP-aligned codon pairs. Rescaling $D^{(\mathrm{syn})}$ and $D^{(\mathrm{ns})}$ to the nucleotide substitution scale on pairs with $d_0 = D_{ij}^{(\mathrm{TN93\text{-}DP})} < 0.35$ gives the depth-adaptive distance:

$$D_{ij} = \alpha_{\mathrm{syn}}(d_0)\, D_{ij}^{(\mathrm{syn})} + \big(1 - \alpha_{\mathrm{syn}}(d_0)\big)\Big(0.85\, D_{ij}^{(\mathrm{TN93\text{-}DP})} + 0.15\, D_{ij}^{(\mathrm{ns})}\Big), \quad \alpha_{\mathrm{syn}}(d_0) = 0.20\, e^{-(d_0/0.35)^2}. \tag{S3}$$

To combine direct pairwise distances $D_{ij}$ with multi-step stepping-stone paths $(i \to k \to j)$ through intermediate relatives without assuming a topology in advance, we model the taxon network as a screened electrical resistor circuit. Each taxon pair $(i, j)$ is linked by a conductor with screened conductance

$$w_{ij} = \frac{\exp(-D_{ij}/\sigma)}{D_{ij} + \epsilon_0}, \tag{S4}$$

where $\sigma = \text{med}(D)$ and $\epsilon_0 = 10^{-3}$. For close relatives $(D_{ij} \ll \sigma)$, $w_{ij} \approx (D_{ij} + \epsilon_0)^{-1}$ provides Fitch–Margoliash-like inverse-variance precision weighting; at deep divergences $(D_{ij} \gg \sigma)$, the exponential screening factor $\exp(-D_{ij}/\sigma)$ suppresses saturated long-range shunts. From the graph Laplacian $L = \text{diag}(W\mathbf{1}) - W$ and its Moore–Penrose pseudoinverse $L^+$, we obtain the Klein–Randić effective resistance $\Omega_{ij} = L^+_{ii} + L^+_{jj} - 2L^+_{ij}$. On an additive tree $T$ with edge conductances $1/t_e$, $\Omega_{ij} = d_T(i, j)$ identically; on a noisy complete distance graph, $\Omega_{ij}$ reduces the four-point tree-additivity defect across deep quartet splits while $O(1/N_C)$ parallel paths inside dense clades compress terminal branches (Table S2 and Fig. S2A). Blending $D$ and scale-matched $\Omega$,

$$D_{\text{ohmic}} = (1 - \alpha_\Omega)\, D + \alpha_\Omega \frac{\sigma}{\text{med}(\Omega)}\, \Omega, \qquad \alpha_\Omega = 0.30, \tag{S5}$$

preserves terminal branch lengths (70% weight) while stabilizing deep internal splits (30% weight).

To extract a rooted binary guide tree $T_0$ from $D_{\text{ohmic}}$ without assuming a molecular clock, we apply the outgroup-referenced rate-subtraction transform. At the root split ($U = V$, $|U| = m$), we set $r_i^{(V)} = \frac{1}{m-2} \sum_{k \in V} D_{\text{ohmic}}(i, k)$; at subsequent splits of a sub-clade $U \subset V$ with external outgroup $A \subset V \setminus U$, we set $r_i^{(A)} = \frac{1}{|A|} \sum_{a \in A} D_{\text{ohmic}}(i, a)$. The outgroup-referenced shared-ancestry matrix,

$$G_{ij}^{(A)} = \frac{1}{2}\left(r_i^{(A)} + r_j^{(A)} - D_{\text{ohmic}}(i, j)\right), \tag{S6}$$

cancels root-to-tip path lengths $L_i$ and $L_j$ on any additive tree ($r_i^{(A)} = c_A + L_i$ and $D_{\text{ohmic}}(i, j) = L_i + L_j - 2S_{ij} \implies G_{ij}^{(A)} = c_A + S_{ij}$, where $S_{ij} \geq 0$ is the shared internal stem below the root of $U$). Subtracting each taxon's cross-root floor $g_i^{\min} = \min_{k \neq i} G_{ik}^{(A)} = c_A$ isolates $\Delta G_{ij}^{(A)} = \max(0, G_{ij}^{(A)} - \frac{1}{2}(g_i^{\min} + g_j^{\min})) = S_{ij}$ (Fig. S2B). Guide-tree splits are extracted via Neighbor-Joining agglomeration on $Q^{\text{NJ}}(D_{\text{ohmic}}) = -2G^{(V)}$ or recursive Fiedler bipartitioning of $\Delta G^{(A)}$ (Table S2 and Figs. S2C, S3).

Figure S3 illustrates the resulting Phase 1 alignment-free guide trees $T_0$ alongside the final Maximum Likelihood phylogenies $T_{\text{ML}}$ across four empirical datasets spanning distinct evolutionary regimes. On mammalian *CD2* ($N = 10$, 179–185 codons; Fig. S3A), $T_0$ recovers all 7/7 internal splits (100.0%, Robinson–Foulds distance RF = 0, cophenetic correlation $r_{\text{coph}} = 0.851$) from unaligned coding sequences, separating Primates, Rodentia, and Laurasiatheria (*Cat*, (*Horse*, (*Cow*, *Pig*))). On raw unaligned GenBank coding sequences of mammalian *IFITM3* ($N = 13$, 127–149 codons with variable-length 5′ N-terminal extensions; Fig. S3B), $T_0$ recovers 9/10 internal splits (90.0%, $r_{\text{coph}} = 0.916$). In this dataset, Euarchontoglires (*Colugo* basal to Primates and Rodentia), basal Chiroptera (*Rhinolophus hipposideros*), basal Suina (*Pig*), and Cetacea (*Bottlenose whale*, (*Baiji dolphin*, *Narwhal*)) match $T_{\text{ML}}$ identically; only the short internal stem (0.0025 substitutions/site) inside the three-taxon Pecoran ruminant radiation (*Giraffe*, *Sheep*, *Fallow deer*) differs prior to Stage 2 refinement. On *Drosophila adh* ($N = 23$, 254 codons; Fig. S3C), despite strong evolutionary rate and third-position GC heterogeneity across subgenera, $T_0$ recovers 17/20 splits (85.0%, $r_{\text{coph}} = 0.958$), cleanly isolating the *melanogaster*, *obscura*, and Hawaiian/*repleta* radiations and confining discrepancies to three short intraclade stems

**Table S2.** *Empirical operating characteristics of Stage 1 Phase 1 Ohmic conductance regularization and rate-corrected guide-tree construction.* Part I compares three conductance kernels at $\alpha_\Omega = 0.30$ and sweeps the effective-resistance blending weight $\alpha_\Omega \in [0, 1]$ under the screened Ohmic kernel Eq. (S4). Entries report the mean four-point tree-additivity defect ($\Delta_{4pt}/\sigma$), quartet concordance with the Maximum Likelihood (ML) phylogeny, and the terminal sister-branch ratio $P_{10}/P_{50}$ (which inflates as $\alpha_\Omega \to 1$ due to parallel intra-clade averaging). Part II compares initial guide-tree split concordance (%) with the ML phylogeny across uncorrected spectral cuts ($W_{\mathrm{RBF}}$), standard Neighbor-Joining on single-track DP distances ($D_{\mathrm{tot}}$), and top-down spectral vs. bottom-up Neighbor-Joining on the multi-track Ohmic distance matrix ($D_{\mathrm{ohmic}}$, Eqs. S3–S6).

| Part I: Circuit Kernel & $\alpha_\Omega$ Sweep | Parameter / Kernel Formula | CD2 ($N$=10) | IFITM3 ($N$=13) | *adh* ($N$=23) | VHH ($N$=212) | OPN1SW ($N$=33) |
|---|---|---|---|---|---|---|
| Pure Ohm's Law ($\alpha_\Omega = 0.30$) | $w_{ij} = (D_{ij} + \epsilon_0)^{-1}$ | 0.1875 / 67.6% | 0.2037 / 94.0% | 0.1245 / 95.8% | 0.3227 / 82.2% | 0.3747 / 68.4% |
| Pure Gaussian RBF ($\alpha_\Omega = 0.30$) | $w_{ij} = \exp(-D_{ij}/\sigma)$ | 0.1886 / 67.6% | 0.2071 / 94.0% | 0.1340 / 95.4% | 0.3230 / 82.1% | 0.3769 / 68.5% |
| *Screened Ohmic ($\alpha_\Omega = 0.30$)* | $w_{ij} = \exp(-D_{ij}/\sigma)/(D_{ij} + \epsilon_0)$ | 0.1871 / 67.6% | 0.2008 / 94.0% | 0.1212 / 95.7% | 0.3224 / 82.2% | 0.3735 / 68.5% |
| No Circuit Regularization | $\alpha_\Omega = 0.00$ ($\Delta_{4pt}$ / $P_{10}/P_{50}$) | 0.1891 / 0.113 | 0.2081 / 0.356 | 0.1345 / 0.180 | 0.3230 / 0.547 | 0.3774 / 0.477 |
| *Default Operating Blend* | $\alpha_\Omega = 0.30$ ($\Delta_{4pt}$ / $P_{10}/P_{50}$) | 0.1871 / 0.105 | 0.2008 / 0.513 | 0.1212 / 0.321 | 0.3224 / 0.542 | 0.3735 / 0.474 |
| Half-Resistance Blend | $\alpha_\Omega = 0.50$ ($\Delta_{4pt}$ / $P_{10}/P_{50}$) | 0.1867 / 0.103 | 0.1910 / 0.542 | 0.1103 / 0.413 | 0.3217 / 0.536 | 0.3687 / 0.461 |
| Pure Effective Resistance | $\alpha_\Omega = 1.00$ ($\Delta_{4pt}$ / $P_{10}/P_{50}$) | 0.1764 / 0.098 | 0.1291 / 0.556 | 0.0686 / 0.636 | 0.2490 / 0.510 | 0.2822 / 0.437 |
| Part II: Guide-Tree Extraction ($T_0$) | Criterion & Distance Input | CD2 Splits | IFITM3 Splits | *adh* Splits | OPN1SW Splits | Mean Recovery |
| Uncorrected Spectral Cut | Clock Affinity $W_{\mathrm{RBF}}(D_{\mathrm{tot}})$ | 100.0% | 70.0% | 40.0% | 36.7% | 61.7% |
| Standard Neighbor-Joining | Single-Track $Q^{\mathrm{NJ}}(D_{\mathrm{tot}})$ | 71.4% | 90.0% | 80.0% | 60.0% | 75.4% |
| *Spectral Shared-Ancestry Cut* | Top-Down Fiedler on $\Delta G^{(A)}(D_{\mathrm{ohmic}})$ | 100.0% | 90.0% | 85.0% | 56.7% | 82.9% |
| *Ohmic Neighbor-Joining* | Bottom-Up NJ on $Q^{\mathrm{NJ}}(D_{\mathrm{ohmic}})$ | 100.0% | 80.0% | 90.0% | 66.7% | 84.2% |

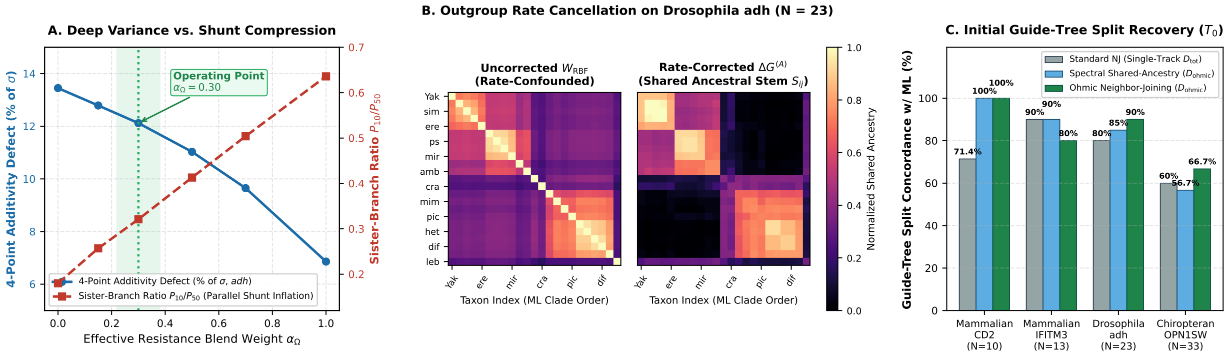

**Figure S2.** *Physical intuition and empirical operating characteristics of Stage 1 Phase 1 Ohmic circuit regularization and rate-corrected guide-tree extraction.* Panel A sweeps the effective-resistance weight $\alpha_\Omega \in [0, 1]$ on *Drosophila adh* ($N = 23$). Increasing $\alpha_\Omega$ monotonically reduces the deep four-point tree-additivity defect ($\Delta_{4pt}/\sigma$, blue circles) via stepping-stone path averaging, whereas values above $\alpha_\Omega = 0.30$ inflate the terminal sister-branch ratio $P_{10}/P_{50}$ (red squares) as $O(1/N_C)$ intra-clade parallel paths compress short terminal branches. Panel B displays the empirical $23 \times 23$ similarity matrices on *Drosophila adh* ordered by Maximum Likelihood clades. Uncorrected Gaussian affinity $W_{\mathrm{RBF}} = \exp(-D_{\mathrm{ohmic}}/\sigma)$ (left) exhibits diffuse off-diagonal cross-clade coupling driven by lineage rate heterogeneity, whereas the outgroup-referenced shared-ancestry matrix $\Delta G^{(A)}$ (right, Eq. S6) cancels root-to-tip branch lengths ($+L_i - L_i = 0$) identically and isolates crisp block-diagonal sub-clades. Panel C reports initial guide-tree ($T_0$) internal split concordance with the ML phylogeny across empirical benchmarks prior to Stage 2 coupled SPR refinement.

(< 0.009 substitutions/site). Finally, chiropteran short-wavelength opsin *OPN1SW* ($N = 33$, 106–286 codons; Fig. S3D) combines intact functional opsins with truncated pseudogenes ($\Psi$) undergoing relaxed purifying selection across a 2.7-fold length range. Even under these severe truncations, $T_0$ preserves monophyly of every major bat family (Yangochiroptera, Megadermatidae, Rhinolophidae, Hipposideridae, and Pteropodidae; 17/30 splits, $r_{\mathrm{coph}} = 0.897$) and restricts topological uncertainty to rapid intrafamilial radiations ($b < 0.005$ substitutions/site) that Stage 2 resolves via coupled

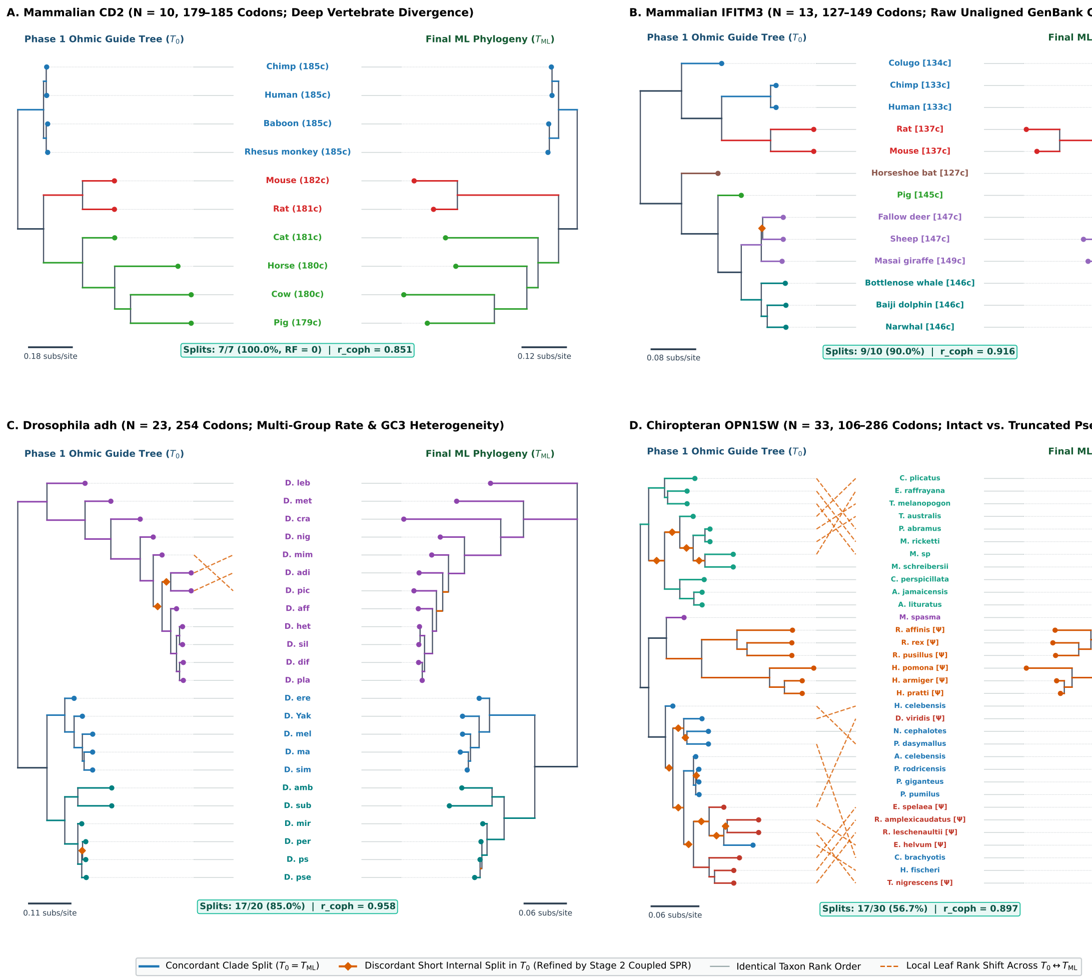

**Figure S3.** *Representative Stage 1 Phase 1 alignment-free Ohmic guide trees ($T_0$, left in each panel) compared against final Maximum Likelihood phylogenies ($T_{ML}$, right in each panel) across four empirical coding datasets.* Solid clade-colored branches indicate internal bipartitions shared identically between $T_0$ and $T_{\mathrm{ML}}$; dashed orange internal branches with diamond markers indicate short internal splits in $T_0$ that are subsequently refined by Stage 2 coupled SPR search. Horizontal gray connectors link leaves with identical vertical rank order across both trees, while dashed orange connectors mark local leaf rank shifts. Panel A: Mammalian *CD2* ($N = 10$, 179–185 codons; 7/7 splits, 100.0%, RF = 0, $r_{\mathrm{coph}} = 0.851$). Panel B: Uncurated GenBank mammalian *IFITM3* transcripts with variable 5′ N-terminal extensions ($N = 13$, 127–149 codons shown in brackets; 9/10 splits, 90.0%, $r_{\mathrm{coph}} = 0.916$). Panel C: *Drosophila adh* across the *melanogaster*, *obscura*, and Hawaiian/*repleta* radiations ($N = 23$, 254 codons; 17/20 splits, 85.0%, $r_{\mathrm{coph}} = 0.958$). Panel D: Chiropteran short-wavelength opsin *OPN1SW* combining functional genes and truncated pseudogenes ($\Psi$) across 33 bat species (106–286 codons; 17/30 splits, 56.7%, $r_{\mathrm{coph}} = 0.897$).

alignment-topology search.

In the second phase, we traverse $T_0$ in bottom-up post-order to progressively align ancestral codon profiles by Felsenstein pruning. Protein-coding genes comprise both a conserved structural core ($\mathcal{C}_{\mathrm{core}}$) and faster-evolving surface loops ($\mathcal{C}_{\mathrm{loop}}$) where indels, radical amino-acid replacements, and multi-nucleotide changes concentrate. We therefore calibrate an empirical two-stratum codon substitution model from the homologous sequence pairs $(u, v)$ retained in Stage 0 ($\geq 35\%$ amino-acid identity). To ensure that internal insertions or deletions never shift downstream codons out of register during calibration, each retained pair $(u, v)$ is aligned via banded affine-gap codon dynamic programming ($B_0 = \max(24, |L_u - L_v| + 8)$, $\gamma^{\text{cal}}_{\text{open}} = 1.10$, $\gamma^{\text{cal}}_{\text{ext}} = 0.30$). Across all aligned non-gap codon pairs $(c_1, c_2)$ at site $k$, we measure the normalized nucleotide divergence $d_{\text{nt}}(k) \in \{0, \frac{1}{3}, \frac{2}{3}, 1\}$ and the Grantham–Miyata amino-acid physicochemical distance $d_{\text{AA}}(k) \in [0, 1]$, and partition sites into $\mathcal{C}_{\text{loop}}$ versus $\mathcal{C}_{\text{core}}$ using a five-codon sliding window $W_k = [k-2, k+2]$. Table S3 summarizes all site quantities, genetic-code neutral reference constants, and closed-form moment estimators evaluated across each stratum $m \in \{\text{core}, \text{loop}\}$ without numerical optimization.

Each stratum $m \in \{\text{core}, \text{loop}\}$ defines a reversible $S \times S$ codon rate generator $\hat{Q}_m$:

$$\hat{Q}_{m,xy} = \begin{cases} \mu_{xy}\, \Phi_{xy}, & \text{if } x \neq y \text{ and } A(x) = A(y), \\ \hat{\omega}_m \, \dfrac{\exp\!\big(-\hat{\lambda}_{\text{AA},m}\, d_{\text{AA}}(A(x), A(y))\big)}{\bar{w}_{\text{AA},m}} \, \delta_{m,|\Delta(x,y)|}\, \mu_{xy}\, \Phi_{xy}, & \text{if } A(x) \neq A(y). \end{cases} \tag{S7}$$

Here $\Phi_{xy} = \prod_{p \in \Delta(x,y)} \pi^{(p)}_{y_p}$ is the target F3×4 positional nucleotide frequency product, and $\mu_{xy} = \prod_{p \in \Delta(x,y)} r_{x_p, y_p}$ is the Tamura–Nei nucleotide exchangeability product ($r_{a,b} = 1$ for purine transitions A $\leftrightarrow$ G, $\hat{\theta}_{CT}$ for pyrimidine transitions C $\leftrightarrow$ T, and $\hat{\theta}_{\text{TV}}$ for transversions). The constant $\bar{w}_{\text{AA},m}$ normalizes the mean physicochemical weight across single-nucleotide nonsynonymous pairs to unity, and $\delta_{m,1} = 1, \delta_{m,2} = \hat{\delta}_{2\text{nt},m}, \delta_{m,3} = \hat{\psi}_{3\text{nt},m}$. Equation (S7) assigns full synonymous exchangeability across the 20 multi-nucleotide synonymous codon pairs in the universal genetic code (Leu `CTN` $\leftrightarrow$ `TTR`, Arg `CGN` $\leftrightarrow$ `AGR`, and Ser `TCN` $\leftrightarrow$ `AGY`), preventing six-fold degenerate synonymous switches from being misclassified as indels on short branches. Because Phase 1 guide-tree branch lengths $t_b$ measure divergence along the conserved synonymous/structural backbone, we evaluate branch transition probabilities at $t_b$ under the empirical two-stratum mixture $P(t_b) = (1 - \hat{w}_{\text{loop}}) \exp(t_b \hat{Q}_{\text{core}}) + \hat{w}_{\text{loop}} \exp(\hat{\rho}_{\text{loop/core}} t_b \hat{Q}_{\text{loop}})$. Each entry $P_{s,x}(t_b) = \mathbb{P}(X_v = x \mid X_u = s, t_b)$ gives the forward probability that an ancestral codon $s$ at parent node $u$ evolves into codon $x$ at child node $v$.

Panel D of Table S3 illustrates how these Stage 0 estimates adapt $\hat{Q}_{\text{core}}$, $\hat{Q}_{\text{loop}}$, and $P(t_b)$ to the selective regime of each gene. In a strongly conserved housekeeping enzyme such as *Drosophila adh* ($\hat{w}_{\text{loop}} = 0.33, \hat{\rho}_{\text{loop/core}} = 4.90$), observed nonsynonymous changes in the core have mean Grantham–Miyata distance $\bar{d}^{\text{obs}}_{\text{AA,core}} = 0.12 \ll \mu_0 = 0.302$, yielding $\hat{\omega}_{\text{core}} = 0.06$ and $\hat{\lambda}_{\text{AA,core}} = 3.50$. Even in the loop stratum of *adh*, purifying and physicochemical constraints remain strict ($\hat{\omega}_{\text{loop}} = 0.18, \hat{\lambda}_{\text{AA,loop}} = 2.33$), so $\hat{Q}_{\text{core}}$ and $\hat{Q}_{\text{loop}}$ suppress polar or charge-reversing replacements such as `CTG` (Leu) $\rightarrow$ `CAG` (Gln) ($0.02 \rightarrow 0.06$) and `GAG` (Glu) $\rightarrow$ `AAG` (Lys) ($0.03 \rightarrow 0.09$) relative to synonymous `CTA` $\rightarrow$ `CTG` ($1.00 \rightarrow 0.77$). In contrast, on immune and cell-surface genes such as mammalian *IL10* ($\hat{\omega} : 0.11 \rightarrow 0.79, \hat{\lambda}_{\text{AA}} : 3.50 \rightarrow 0.66$), *CD2* ($\hat{\omega} : 0.32 \rightarrow 1.16, \hat{\lambda}_{\text{AA}} : 2.70 \rightarrow 0.00$), and primate *APOBEC3H* ($\hat{\omega} : 0.40 \rightarrow 1.50, \hat{\lambda}_{\text{AA}} : 3.50 \rightarrow 0.00$), core structural domains remain conserved while surface loops exhibit unrestrained physicochemical turnover ($\bar{d}^{\text{obs}}_{\text{AA,loop}} = 0.28$–$0.40$) and elevated multi-hit rates ($\hat{\delta}_{2\text{nt,loop}} = 0.09$–$0.26$). In *CD2*, the scaled instantaneous rate for `CTG` $\rightarrow$ `CAG` rises from 0.17 in $\hat{Q}_{\text{core}}$ to 0.48 in $\hat{Q}_{\text{loop}}$ (exceeding synonymous `CTG` $\rightarrow$ `CTA` $= 0.33$), and `GAG` $\rightarrow$ `AAG` rises from 0.24 to 0.63. Across a typical sister-branch pair ($t_L = t_R = 0.10$), the resulting mixture $P(t_b)$ assigns negative Felsenstein match log-Bayes factors (Eq. S11) to polar and charged replacements in *adh* ($\mathcal{E}(\texttt{CTG}, \texttt{CAG}) = -0.29$, $\mathcal{E}(\texttt{GAG}, \texttt{AAG}) = -0.03$), but positive homology support in *IL10* (+0.17 and +0.42), *APOBEC3H* (+0.44 and +0.52), and *CD2* (+1.09 and +0.55). Meanwhile, non-homologous multi-nucleotide mismatches remain negative across all datasets ($\mathcal{E}(\texttt{CTG}, \texttt{GAG}) = -1.34$ in *adh*, $-1.07$ in *IFITM3*, and $-0.54$ in *CD2*).

At any node $v \in T_0$ subtending $N_v$ descendant leaves, we represent the partial in-frame alignment of its clade as a codon profile $\mathcal{P}_v$ of length $M_v$ columns. Let $D_{v,c}$ denote the observed leaf codons at column $c \in \{1, \dots, M_v\}$ across the clade of $v$, and let $\mathbb{P}(D_{v,c} \mid X_v = x)$ be the standard Felsenstein conditional probability of observing $D_{v,c}$ given ancestral sense codon $X_v = x \in \{1, \dots, S\}$ at node $v$. Under the stationary codon distribution $\boldsymbol{\pi}$, the marginal probability of observing column $D_{v,c}$ is $\mathbb{P}(D_{v,c}) = \sum_{y=1}^{S} \pi_y \, \mathbb{P}(D_{v,c} \mid X_v = y)$. Each column $c$ of $\mathcal{P}_v$ stores a leaf coordinate map and a fractional taxon occupancy $o_c^{(v)} \in (0, 1]$, defined as the proportion of $v$'s leaves carrying a non-gap codon at column $c$. It also stores a rescaled Felsenstein likelihood vector $\boldsymbol{\ell}_c^{(v)} \in \mathbb{R}_{\geq 0}^{S}$, defined as the conditional clade likelihood divided by the marginal clade probability:

$$\ell_{c,x}^{(v)} = \frac{\mathbb{P}(D_{v,c} \mid X_v = x)}{\mathbb{P}(D_{v,c})} = \frac{\mathbb{P}(D_{v,c} \mid X_v = x)}{\sum_{y=1}^{S} \pi_y \, \mathbb{P}(D_{v,c} \mid X_v = y)}, \qquad \text{so that } \sum_{x=1}^{S} \pi_x \, \ell_{c,x}^{(v)} = 1. \tag{S8}$$

By Bayes' rule, $\pi_x \ell_{c,x}^{(v)} = \mathbb{P}(X_v = x \mid D_{v,c})$ is the marginal posterior probability of ancestral codon $x$ at node $v$ given $D_{v,c}$, and $\ell_{c,x}^{(v)}$ is the posterior-to-prior likelihood ratio for state $x$. Rescaling by $\mathbb{P}(D_{v,c})$ keeps $\boldsymbol{\ell}_c^{(v)}$ unit-mean at every height of the tree, preventing numerical underflow and factoring out each column's null marginal probability in advance. At a leaf taxon $t$ with observed sense codon $c_{t,k}$ at position $k$ (before traversing its parent branch), $\mathbb{P}(D_{t,k} \mid X_t = x) = \mathbb{I}[x = c_{t,k}]$ and $\mathbb{P}(D_{t,k}) = \pi_{c_{t,k}}$. Thus $\mathcal{P}_t$ is initialized with unit occupancy $o_k^{(t)} = 1$ and $\ell_{k,x}^{(t)} = \mathbb{I}[x = c_{t,k}]/\pi_{c_{t,k}}$ (or $\boldsymbol{\ell}_k^{(t)} = \mathbf{1}$ for ambiguous triplets). At an internal node $u$ with left and right children $(v_L, v_R)$ and branch lengths $(t_L, t_R)$, we express each child column's conditional likelihood as a function of the parent state $X_u = s$ by marginalizing over the intermediate child state $x \in \{1, \dots, S\}$ along the top-down branch transition $u \to v$:

$$\tilde{\boldsymbol{\ell}}_i^{(L)} = P(t_L) \, \boldsymbol{\ell}_i^{(L)}, \qquad \tilde{\boldsymbol{\ell}}_j^{(R)} = P(t_R) \, \boldsymbol{\ell}_j^{(R)}. \tag{S9}$$

For each candidate ancestral codon $s \in \{1, \dots, S\}$ at parent $u$, the $s$-th component $\tilde{\ell}_{i,s}^{(L)} = \sum_{x=1}^{S} P_{s,x}(t_L) \, \ell_{i,x}^{(L)} = \mathbb{P}(D_{L,i} \mid X_u = s)/\mathbb{P}(D_{L,i})$ is the conditional probability of generating left-clade column $D_{L,i}$ down branch $t_L$ from state $X_u = s$, divided by $\mathbb{P}(D_{L,i})$. Because $\hat{Q}_{\text{core}}$ and $\hat{Q}_{\text{loop}}$ share the same Laplace-smoothed F3×4 equilibrium distribution $\boldsymbol{\pi}$ while relaxing nonsynonymous exchangeabilities in loops, the mixture matrix $P(t_b)$ satisfies $\boldsymbol{\pi}^\top P(t_b) = \boldsymbol{\pi}^\top$ for every $t_b \geq 0$. Consequently, $\tilde{\boldsymbol{\ell}}_i^{(L)}$ and $\tilde{\boldsymbol{\ell}}_j^{(R)}$ retain unit stationary expectation ($\sum_{s=1}^{S} \pi_s \tilde{\ell}_{i,s}^{(L)} = 1$) with strictly positive entries across all $S$ codons whenever $t_L, t_R > 0$. Using $\boldsymbol{\pi}$ as the common prior for both homologous and independent-origins clades is required for this invariance: under any non-stationary prior $\mathbf{q} \neq \boldsymbol{\pi}$, merely lengthening a branch $t_L$ above an unaligned clade would alter its null marginal probability. We then align $\mathcal{P}_L$ (length $M_L$) and $\mathcal{P}_R$ (length $M_R$) via three-move affine-gap dynamic programming (Gotoh's algorithm) over $(i, j) \in \{0, \dots, M_L\} \times \{0, \dots, M_R\}$:

1. *Homologous match (M, $(i-1, j-1) \to (i, j)$):* Aligns column $i$ of $\mathcal{P}_L$ with column $j$ of $\mathcal{P}_R$ as descendants of a single shared ancestral codon $X_u = s \sim \boldsymbol{\pi}$ at parent $u$. Under shared ancestry at $u$, conditional independence of the two child branches gives joint probability $\mathbb{P}(D_{L,i}, D_{R,j} \mid \text{homologous}) = \sum_{s=1}^{S} \pi_s \, \mathbb{P}(D_{L,i} \mid X_u = s) \, \mathbb{P}(D_{R,j} \mid X_u = s)$, whereas under independent origins (separate indels) the joint probability is $\mathbb{P}(D_{L,i}) \, \mathbb{P}(D_{R,j})$. Because $\tilde{\boldsymbol{\ell}}_i^{(L)}$ and $\tilde{\boldsymbol{\ell}}_j^{(R)}$ are already divided by $\mathbb{P}(D_{L,i})$ and $\mathbb{P}(D_{R,j})$, their $\boldsymbol{\pi}$-weighted inner product equals the exact homology Bayes factor:

$$\text{BF}(i, j) = \frac{\mathbb{P}(D_{L,i}, D_{R,j} \mid \text{homologous})}{\mathbb{P}(D_{L,i}) \, \mathbb{P}(D_{R,j})} = \sum_{s=1}^{S} \pi_s \, \tilde{\ell}_{i,s}^{(L)} \, \tilde{\ell}_{j,s}^{(R)}. \tag{S10}$$

Equation (S10) generalizes classic pairwise PAM/BLOSUM log-odds ratios ($\mathrm{BF}(x,y) = P_{x,y}(t_L + t_R)/\pi_y$ when $v_L, v_R$ are leaves) to arbitrary phylogenetic subtrees. Across successive merges from the tips to the root, intermediate clade marginals $\mathbb{P}(D_{v,c})$ telescope out so that the product of node Bayes factors equals the full-tree likelihood ratio $\mathbb{P}(D_{\mathrm{root},c})/\prod_t \pi_{c_t}$. We score the match by its regularized log-Bayes factor and update parent $u$'s rescaled likelihood vector for the merged column $D_{u,c} = (D_{L,i}, D_{R,j})$ using $\mathrm{BF}(i,j)$ as the normalizer:

$$\mathcal{E}(i,j) = \ln\big(\mathrm{BF}(i,j) + \epsilon_0\big), \qquad \ell^{(u)}_{c,s} = \frac{\mathbb{P}(D_{u,c} \mid X_u = s)}{\mathbb{P}(D_{u,c})} = \frac{\tilde{\ell}^{(L)}_{i,s}\,\tilde{\ell}^{(R)}_{j,s}}{\mathrm{BF}(i,j)}. \tag{S11}$$

Here $\epsilon_0 = 0.22$ is a Laplace floor that bounds the worst-case mismatch penalty at $\ln(\epsilon_0) \approx -1.51$ while keeping non-homologous codon pairs strictly negative, and logarithmic scoring ensures that rare-codon matches ($1/\pi_x$) increase $\mathcal{E}(i,j)$ only additively ($\ln(1/\pi_x)$). The emitted parent column sets its occupancy to $o^{(u)}_c = w_L o^{(L)}_i + w_R o^{(R)}_j$, where $w_L = N_L/(N_L + N_R)$ and $w_R = 1 - w_L$. It merges both children's leaf coordinate maps and induces the marginal ancestral codon posterior $d^{(u)}_{c,s} = \pi_s \ell^{(u)}_{c,s}$.

2. *Left-clade insertion / right-clade deletion ($I$, $(i-1,j) \to (i,j)$):* Advances column $i$ of $\mathcal{P}_L$ while placing a codon gap (---) across all $N_R$ leaves of $v_R$. Because a gap in clade $R$ imposes no character constraint ($\tilde{\boldsymbol{\ell}}^{(R)} = \mathbf{1}$ and $\sum_s \pi_s \tilde{\ell}^{(L)}_{i,s} = 1$, giving $\mathrm{BF} = 1$), Felsenstein's rule emits $\boldsymbol{\ell}^{(u)}_c = \tilde{\boldsymbol{\ell}}^{(L)}_i$ with diluted occupancy $o^{(u)}_c = w_L o^{(L)}_i$.

3. *Right-clade insertion / left-clade deletion ($D$, $(i,j-1) \to (i,j)$):* Advances column $j$ of $\mathcal{P}_R$ while placing a codon gap (---) across all $N_L$ leaves of $v_L$, emitting $\boldsymbol{\ell}^{(u)}_c = \tilde{\boldsymbol{\ell}}^{(R)}_j$ with diluted occupancy $o^{(u)}_c = w_R o^{(R)}_j$.

Standard progressive aligners penalize gaps identically regardless of whether an indel is ancestral or clade-specific (the "once a gap, always a gap" bias analyzed by Loytynoja and Goldman). We instead scale the affine gap-opening ($M \to I/D$, $\gamma_{\mathrm{open}} = 2.85$) and gap-extension ($I \to I$, $D \to D$, $\gamma_{\mathrm{ext}} = 0.65$) penalties linearly by the child column's clade occupancy (including along the $i = 0$ and $j = 0$ DP boundaries). When pairing partial columns in state $M$, we also include a symmetric missing-occupancy indel term:

$$\begin{aligned} &\gamma^{(I)}_{\mathrm{open}}(i) = \gamma_{\mathrm{open}}\, o^{(L)}_i, \qquad \gamma^{(I)}_{\mathrm{ext}}(i) = \gamma_{\mathrm{ext}}\, o^{(L)}_i, \qquad \gamma^{(D)}_{\mathrm{open}}(j) = \gamma_{\mathrm{open}}\, o^{(R)}_j, \qquad \gamma^{(D)}_{\mathrm{ext}}(j) = \gamma_{\mathrm{ext}}\, o^{(R)}_j, \\ &\mathcal{S}_M(i,j) = \mathcal{E}(i,j) - \gamma_{\mathrm{open}}\big(1 - \min(o^{(L)}_i, o^{(R)}_j)\big). \end{aligned} \tag{S12}$$

Here $o^{(L)}_i \in (0,1]$ serves as the empirical prior probability that column $i$ was already present at ancestor $v_L$. A full-occupancy column ($o^{(L)}_i = 1$) is present in every leaf of the left clade, so aligning it to a gap in the right clade implies a genuine indel event along branch $t_L$ or $t_R$ and incurs the full cost ($\gamma_{\mathrm{open}}, \gamma_{\mathrm{ext}}$), whereas a partial column ($o^{(L)}_i < 1$) arose from an indel inside the left subtree and propagates across node $u$ with proportionally reduced cost. Conversely, the parameter-free missing-occupancy term $-\gamma_{\mathrm{open}}(1 - \min(o^{(L)}_i, o^{(R)}_j))$ in $\mathcal{S}_M(i,j)$ vanishes identically when full-occupancy columns merge ($o^{(L)}_i = o^{(R)}_j = 1$) and prevents independent low-occupancy insertions ($o^{(L)}_i, o^{(R)}_j \ll 1$) from collapsing into the same column unless their homology Bayes factor $\mathrm{BF}(i,j)$ genuinely supports shared ancestry. When the post-order pass reaches the root of $T_0$, the leaf coordinate map of $\mathcal{P}_{\mathrm{root}}$ assembles the initial $N \times C_0$ in-frame codon alignment $\mathcal{A}_0$ (see Fig. S4 for a complete five-taxon, eight-codon numerical walkthrough).

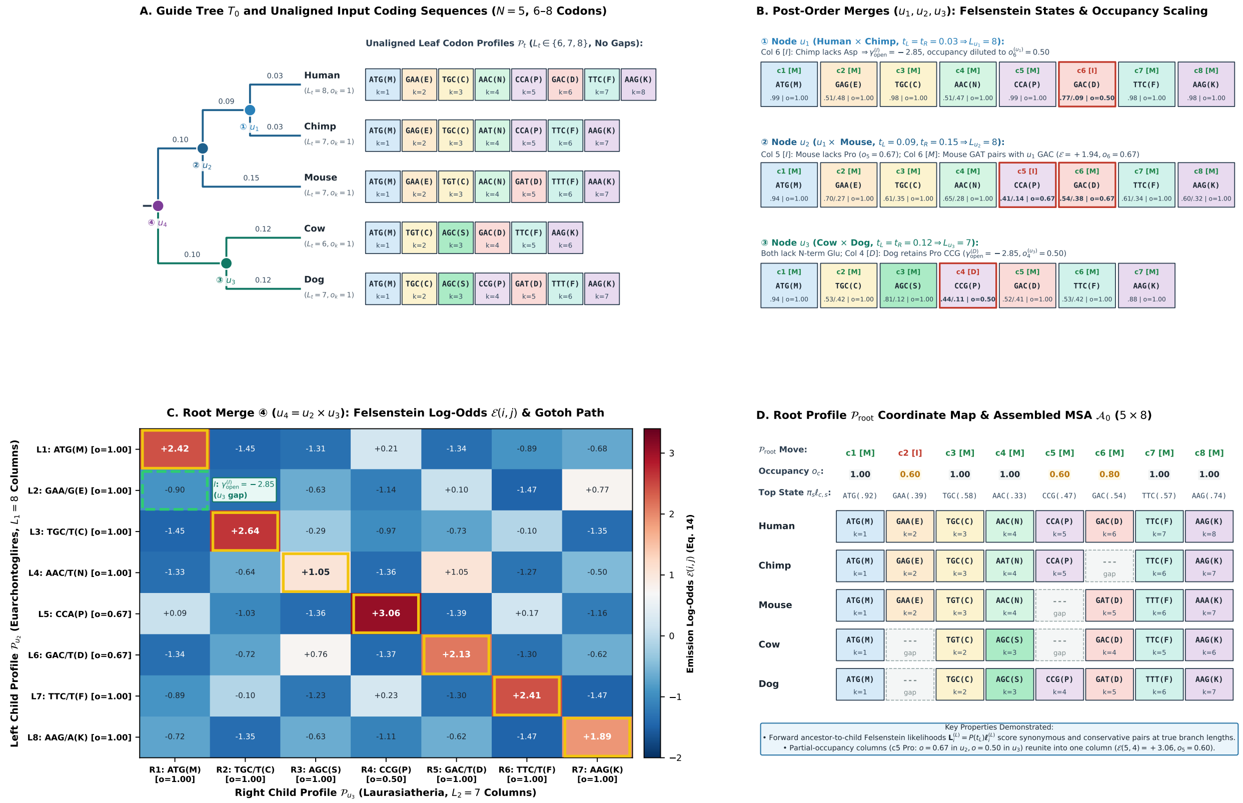

**Figure S4.** *Worked five-taxon, eight-codon walkthrough of Stage 1 Phase 2 progressive codon profile alignment along guide tree* $T_0$. Panel A: rooted guide tree $T_0$ with post-order merge schedule (① $u_1 \to$ ② $u_2 \to$ ③ $u_3 \to$ ④ $u_4$) and unaligned input coding sequences ($M_t \in \{6, 7, 8\}$ codons). Panel B: ancestral codon profiles $\mathcal{P}_{u_1}$, $\mathcal{P}_{u_2}$, and $\mathcal{P}_{u_3}$, showing Gotoh transducer moves $(M, I, D)$, top ancestral sense-codon marginal probabilities $\pi_s \ell_{c,s}^{(u)}$, and fractional clade occupancy $o_c^{(u)} \in (0, 1]$. Panel C: $8 \times 7$ Felsenstein emission log-odds matrix $\mathcal{E}(i, j)$ (Eq. S11) at the root merge $u_4 = u_2 \times u_3$, where gold boxes trace homologous match steps $(M)$ and the dashed green box marks the single insertion step $(I, \gamma_{\text{open}}^{(I)} = -2.85)$ for the Euarchontoglires-specific Glu codon $(L_2)$. Panel D: resulting root profile $\mathcal{P}_{\text{root}}$ and $5 \times 8$ in-frame codon alignment $\mathcal{A}_0$.

**Phase 3: Leave-one-out Inside–Outside cavity Pair-HMM refinement and POA graph construction.** To break the conditioning of $\mathcal{A}_0$ on the bottom-up merge order, Phase 3 refines all gapped regions via leave-one-out Felsenstein Inside–Outside cavities and a two-state Pair-HMM. We first optimize branch lengths $t_b$ on $(T_0, \mathcal{A}_0)$ and evaluate branch transition matrices $P_b = P(t_b)$ under the empirical two-stratum generator $(1 - \hat{w}_{\text{loop}}) \exp(t_b \hat{Q}_{\text{core}}) + \hat{w}_{\text{loop}} \exp(\hat{\rho}_{\text{loop/core}} t_b \hat{Q}_{\text{loop}})$ recalibrated on $\mathcal{A}_0$. Window extraction is seeded strictly by columns containing at least one internal gap: gapped columns separated by at most 2 ungapped columns are coalesced into a contiguous run, padded by 2 ungapped flanking anchor codons on each side, and covered by 50%-overlapping half-stride tiles (stride = 9 codons, maximum tile width $|W|_{\max} = 18$ codons). For each window $W = [c_{\text{start}}, c_{\text{end}})$, a two-pass Felsenstein Inside–Outside traversal under $P_b$ computes the leave-one-taxon-out tree cavity distribution $\mathbf{u}_{t,w}^{\text{tree}} = \mathbb{P}(X_{t,w} = \cdot \mid \mathcal{A}_{-t,w}, T_0) \in \Delta^{S-1}$ at every leaf $t$ and window column $w \in \{1, \ldots, |W|\}$ in $O(N \cdot |W|)$ time. To prevent a locally misresolved binary split in $T_0$ from biasing the cavity under reticulation or incomplete lineage sorting, we query $\Omega$ for taxon $t$'s lowest-resistance non-sister partner $u^*(t) = \arg\min_{u \notin \mathrm{Sib}(t)} \Omega_{tu}$ and form the Ohmic-augmented cavity distribution:

$$\mathbf{u}_{t,w} = \big(1 - w_{\mathrm{shunt}}(t)\big)\, \mathbf{u}^{\mathrm{tree}}_{t,w} + w_{\mathrm{shunt}}(t)\, P(d_{\mathrm{shunt}})^\top \mathbf{e}_{c_{u^*(t),w}}, \tag{S13}$$

where $w_{\mathrm{shunt}}(t) = \mathrm{clamp}(0.25\, g_{\mathrm{shunt}}/(g_{\mathrm{sib}} + g_{\mathrm{shunt}}), 0.02, 0.20)$.

For each taxon $t$ carrying $M_t \in (0, |W|)$ non-gap codons $(y_{t,1}, \ldots, y_{t,M_t})$ inside $W$, we strip its existing gaps and model its placement across the $|W|$ columns as a hidden Markov state sequence $\mathbf{z}_t \in \{M, G\}^{|W|}$. Emitting codon $y_{t,j}$ in state $z_{t,w} = M$ receives the column-mean-centered log-odds score:

$$E_t(j, w) = \ln u_{t,w}(y_{t,j}) - \frac{1}{S} \sum_{s=1}^{S} \ln u_{t,w}(s) + 2.4\, I_w - 1.5\,(1 - I_w), \tag{S14}$$

where $I_w = \mathrm{clamp}((\max_s u_{t,w}(s) - 1/S)/(0.20 - 1/S), 0, 1)$ measures column concentration, and entering/extending $z_{t,w} = G$ incurs $-\gamma_{\mathrm{open}}$ / $-\gamma_{\mathrm{ext}}$ plus the column-concentration gap prior $g_{\mathrm{col}}(w) = -3.0\,(\max_s u_{t,w}(s))\, I_w$. Running Viterbi (max) and Forward–Backward (log-sum-exp) recurrences over the $|W| \times M_t$ lattice in $O(|W| \cdot M_t)$ time yields the MAP placement $\hat{\mathbf{z}}_t$, the marginal column homology posterior $PP_c \in [0, 1]$, and the posterior weights of alternative placements recorded in the POA graph. Candidate taxon moves are applied via sequential Gauss–Seidel updates (refreshing $\mathbf{u}_{t,w}$ after each move across up to three bidirectional sweeps) and are accepted if and only if they strictly increase the joint two-stratum substitution and 2-state Felsenstein affine-indel log-likelihood:

$$\ln \mathbb{P}(\mathcal{A}, T \mid \hat{Q}_{\mathrm{core}}, \hat{Q}_{\mathrm{loop}}, \hat{\lambda}_{\mathrm{open}}, \hat{\varepsilon}_{\mathrm{ext}}) = \sum_{c=1}^{C} \ln L_{2\text{-stratum}}\Big(\mathcal{A}_{\cdot,c} \;\Big|\; T, \hat{Q}_{\mathrm{core}}, \hat{Q}_{\mathrm{loop}}\Big) + \sum_{c=1}^{C} \ln L_{\mathrm{indel}}\Big(\mathbf{I}_{\cdot,c} \;\Big|\; \mathbf{I}_{\cdot,c-1}, T, \hat{\lambda}_{\mathrm{open}}, \hat{\varepsilon}_{\mathrm{ext}}\Big) \tag{S15}$$

where $L_{\mathrm{indel}}(\mathbf{I}_{\cdot,c} \mid \mathbf{I}_{\cdot,c-1}, T)$ marginalizes binary codon presence/absence $I_{u,c} \in \{0, 1\}$ across all internal nodes via $2 \times 2$ Felsenstein sum–product pruning with branch transition probability $p^{(b,c)}_{1\to 0} = (1 - q^{(1\to 0)}_{b,c-1})\, \frac{1}{2}(1 - e^{-\hat{\lambda}_{\mathrm{open}} t_b}) + q^{(1\to 0)}_{b,c-1} \hat{\varepsilon}_{\mathrm{ext}}$. When invoked on a pre-existing alignment (`-a`), WALRUS bypasses Phase 1–2 progressive assembly and directly executes Phase 3 local indel-boundary audit, POA homology posterior estimation ($PP_c$), and terminal non-homologous overhang trimming.

#### 1.3 Stage 2: Coupled Ohmic SPR Search and Kadane Segment-Specific Routing

Stage 2 addresses two distinct topological failure modes: (A) global guide-tree indel lock-in, where an initial guide-tree error freezes gap placements against the wrong sister clade, and (B) local intragenic recombination or gene conversion, where no single tree topology holds across the entire coding sequence.

**Part A (Global Coupled SPR and Boundary Indel Realignment).** Candidate subtree-prune-and-regraft (SPR) moves $(u \to v)$ are ranked by their Ohmic stretch ratio $\Omega_{uv}/D^{(\mathrm{syn})}_{uv}$ and Wheatstone bridge conductance on the resistor network, augmented by nearby NNI/SPR target edges within path radius 3. Because evaluating $T_{u\to v}$ on the fixed alignment $\mathcal{A}^{(1)}$ penalizes valid topological moves whenever gap columns in clade $\mathcal{C}_u$ remain aligned to its old neighbor instead of its new sister clade $\mathcal{C}_v$, every candidate $T_{u\to v}$ undergoes a coupled three-step update: (i) 1D Brent coordinate ascent on the four local branch lengths $(t_u, t_v, t_w, t_s)$ around the regraft junction (accelerated via `ScalableWoodbury` low-rank updates on large trees); (ii) re-running the Stage 1 Phase 3 tree-cavity transducer (Eq. S14) across all indel windows for leaves $t \in \mathcal{C}_u$ under the updated Outside cavities $\mathbf{u}_{t,w}(T_{u\to v})$ to produce $\mathcal{A}_{u\to v}$; and (iii) a final local branch-length polish whenever any gap placement changes. Candidate pairs $(T_{u\to v}, \mathcal{A}_{u\to v})$ are accepted if and only if they strictly improve the joint substitution and affine-indel log-likelihood of Eq. (S15).

**Part B (Localized Recombinant Segment Detection, Permutation/BIC Gating, and Cavity Routing).** We next scan the converged backbone tree $T_{\text{backbone}}$ for localized horizontal transfers or recombinant tracts $(u \to v)$ affecting a contiguous sub-segment of the gene. Candidate moves are proposed where leaves $i \in \mathcal{C}_u$ exhibit excess sliding-window codon identity to $\mathcal{C}_v$ relative to their backbone sibling clade $\mathcal{C}_{\text{sib}(u)}$, weighted by Ohmic stretch chords. To prevent donor–recipient branch-length inflation on $T_{\text{backbone}}$ from suppressing localized transfer likelihoods and to maintain symmetric degrees of freedom under site-to-site rate heterogeneity, we evaluate the site-wise log-likelihood differential at each codon $c \in \{1, \ldots, C\}$ using a symmetric two-point local branch-scale envelope on both topologies:

$$\Delta\ell_c = \max_{\tau \in \{\text{std},\text{short}\}} \ell_c\left(T^{(\tau)}_{u\to v}\right) - \max_{\tau \in \{\text{std},\text{short}\}} \ell_c\left(T^{(\tau)}_{\text{backbone}}\right), \tag{S16}$$

where $\tau$ = std retains baseline branch lengths (splitting target edge $t_v$ equally at regraft node $w$) and $\tau$ = short evaluates a near-tip transfer scale ($0.20\, t_u$ with an 85/15 split of the recipient stem edge on both $T_{\text{backbone}}$ and $T_{u\to v}$). Applying Kadane's $O(C)$ dynamic programming recurrence $S_c = \max(0, S_{c-1} + \Delta\ell_c)$ identifies the maximum contiguous subarray $[c^*_{\text{start}}, c^*_{\text{end}}] = \arg\max_{1\le a\le b\le C} \sum_{c=a}^{b} \Delta\ell_c$ with peak cumulative log-likelihood gain $S_{\max}$. To control look-elsewhere inflation over the $O(C^2)$ sub-intervals, $S_{\max}$ is calibrated via $B_{\text{perm}} = 2{,}000$ random permutations of the site order of $\{\Delta\ell_1, \ldots, \Delta\ell_C\}$, preserving the exact marginal distribution of per-site log-likelihood differences while breaking spatial contiguity. Acceptance requires three simultaneous gates: (1) Bonferroni-corrected permutation $p_{\text{perm}} \le 0.01$, (2) Schwarz BIC threshold $2S_{\max} > 3\ln C$, and (3) minimum span $|c^*_{\text{end}} - c^*_{\text{start}} + 1| \ge 8$ codons. For each accepted segment, $(t_u, t_v, t_w)$ are polished on $[c^*_{\text{start}}, c^*_{\text{end}}]$ and assigned a dedicated regraft node index $w = |V(T_{\text{backbone}})|$. All downstream Stage 4 cavity computations and Stage 5 selection tests for $c \in [c^*_{\text{start}}, c^*_{\text{end}}]$ route through the local topology $T_{u\to v}$.

#### 1.4 Stage 3: Phylogenetic Directional Drift Audit, Automatic Root Placement, and 3-Regime Softmax Kernel

A classical $N \times 4$ Pearson $\chi^2$ contingency table on third-codon-position ($\text{CP}_3$) base counts treats tip sequences as independent draws and is therefore vulnerable to Felsenstein clade pseudoreplication, while assigning branch regimes from static tip offsets $(\text{GC}_v - \overline{\text{GC}})$ misclassifies lineages undergoing an AT-ward reversal inside a GC-rich clade. Stage 3 eliminates both artifacts in three steps:

1. *Phylogenetic GLS ancestral reconstruction, Inside–Outside substitution flux, and automatic directional rooting.* Internal node compositions $\widehat{\text{GC}}_u$ are estimated via continuous-trait generalized least squares (GLS), weighting child messages by inverse branch length $1/\max(10^{-3}, t_b)$ in a post-order upward pass followed by pre-order smoothing. Simultaneously, an Inside–Outside cavity pass under the stationary MG94 baseline computes the exact posterior expected AT $\to$ GC ($N^{(b)}_{W\to S}$) and GC $\to$ AT ($N^{(b)}_{S\to W}$) codon substitution counts and standardized directional flux $Z_b = (N^{(b)}_{W\to S} - N^{(b)}_{S\to W})/\sqrt{N^{(b)}_{W\to S} + N^{(b)}_{S\to W} + 2}$ along every branch $b = (p \to v)$. When an unrooted tree is supplied, WALRUS automatically places the directional root along the internal branch $b^*_{\text{root}}$ that maximizes the bipartition compositional separation $|\widehat{\text{GC}}_{C_b} - \widehat{\text{GC}}_{\bar{C}_b}|\sqrt{\min(|C_b|, N - |C_b|)}$ concordant with the net directional substitution flux $Z_b$.

2. *Reversal-aware branch drift trajectory and 3-regime Softmax kernel.* Each branch's directional trajectory $d_b$ combines its parent-to-child step $\Delta_{\text{step}}(b) = \widehat{\text{GC}}_v - \widehat{\text{GC}}_p$, background offset $\Delta_{\text{bg}}(b) = \widehat{\text{GC}}_v - \widehat{\text{GC}}_{\text{root}}$, and codon flux asymmetry $A_b = (N^{(b)}_{W\to S} - N^{(b)}_{S\to W})/(N^{(b)}_{W\to S} + N^{(b)}_{S\to W} +$

4). Whenever a lineage reverses against its clade background ($\Delta_{\text{step}}(b)\,\Delta_{\text{bg}}(b) < 0$), $d_b$ is anchored to the parent-to-child step and substitution flux ($d_b = 0.65\,\Delta_{\text{step}}(b) + 0.14\,A_b$) so reversals are assigned to their true evolutionary direction. Branch regime weights $w_{b,k} = \exp(\beta_{\text{gate}} s_{b,k}) / \sum_{k'=0}^{2} \exp(\beta_{\text{gate}} s_{b,k'})$ over $K_{\text{reg}} = 3$ directional MG94×REV rate generators $Q^{(k)}(1,\omega)$ ($\psi^{(0)} < 0$ AT-enriching, $\psi^{(1)} = 0$ stationary, $\psi^{(2)} > 0$ GC-enriching, where target nucleotide frequencies are tilted by $\pi_j^{(k)} \propto \pi_j \exp(\psi^{(k)} \mathbb{I}[j \in \{G,C\}])$) are mapped onto the backbone and routed SPR topologies by exact leaf-bipartition matching:

$$P_b(t_b \mid \alpha, \omega) = \sum_{k=0}^{2} w_{b,k} \exp\Big(\alpha t_b Q^{(k)}(1,\omega)\Big)\,. \tag{S17}$$

3. *Phylogenetic Directional LRT/BIC stationarity gate.* The 3-regime directional kernel is activated ($K_{\text{reg}} = 3$, activating directional `dFEL` in Stage 5) if and only if $\text{CP}_3$ heterogeneity is significant and the phylogenetic directional test statistic $\Lambda_{\text{dir}} = \max\big(2(\ln L_{\text{dir}}(T) - \ln L_{\text{stat}}(T)),\, \max_b Z_b^2 - 2.5\big)$ satisfies $p_{\chi^2_2}(\Lambda_{\text{dir}}) \le 0.05$ and $\Lambda_{\text{dir}} \ge \min(6.5,\, 1.5 \ln C)$ (with extreme-value threshold $2 \ln B$ for large trees); otherwise $K_{\text{reg}} = 1$ ($\psi^{(1)} = 0$) is retained.

#### 1.5 Stage 4: Shared 16-Point Spectral Grid, $K = 3$ POA Ensemble Cavities, Sufficiency Tensor, and Erlang-3 Error-Sink HMM

To eliminate redundant tree traversals across selection tests while preserving exact site-specific synonymous rate scaling $\tilde{\alpha}_c$ and segment-routed branch lengths $t_{b,c}$, `WALRUS` pre-diagonalizes each active directional rate matrix $Q^{(k)}(1,\omega_g) = R_{k,g} \Lambda_{k,g} L_{k,g}$ across a 16-point $\omega$ grid $\mathcal{G} = \{0.0, 0.02, 0.05, 0.10, 0.20, 0.35, 0.50, 0.75, 1.$[illegible]
Stage 4 constructs the shared sufficiency tensor in three steps:

1. *Bayes-updated $K = 3$ POA ensemble and 3-way Wheatstone bridge cavities.* At each site $c$ with column homology posterior $PP_c < 0.99$, up to $K = 3$ candidate column realizations (the MAP column and adjacent indel-boundary shift states) are evaluated on the active topology $T_c$ (backbone or routed SPR tree) at $(\hat{\alpha}_c, \hat{\omega}_c)$. At any internal quartet hub $u$ carrying a local Ohmic Wheatstone bridge over subtrees $(A, B, C)$ with conductance weights $(w_0, w_1, w_2)$ and resolution branch transition matrices $(P_{\text{br},0}, P_{\text{br},1}, P_{\text{br},2})$, the upward Inside vector is $\mathbf{L}_u = \sum_{r=0}^{2} w_r (P_{\text{br},r}(\mathbf{M}_A \odot \mathbf{M}_B)) \odot \mathbf{M}_C$, permuting $(A, B, C)$ across the three NNI resolutions alongside the downward Outside adjoint messages $(\mathbf{U}_A, \mathbf{U}_B, \mathbf{U}_C)$. Each POA realization's Pair-HMM prior weight $w_{c,m}^{\text{prior}}$ is Bayes-updated by its phylogenetic column log-likelihood $\ell_c^{(m)}$,

$$\tilde{w}_{c,m} = \frac{w_{c,m}^{\text{prior}} \exp\big(\ell_c^{(m)} - \ell_c^{\max}\big)}{\sum_{m'=1}^{K} w_{c,m'}^{\text{prior}} \exp\big(\ell_c^{(m')} - \ell_c^{\max}\big)}, \tag{S18}$$

suppressing shift realizations that introduce phylogenetically incompatible codon clashes before forming the posterior-weighted Inside ($\bar{\mathbf{L}}_{b,c}$) and Outside ($\bar{\mathbf{U}}_{b,c}$) cavity vectors.

2. *Conjugate Empirical Bayes $\tilde{\alpha}_c$ and eigenbasis Hadamard projection.* Plugging unregularized per-site maximum-likelihood estimates $\hat{\alpha}_c = 0$ into branch-site mixtures artificially collapses branch lengths at zero-synonymous-change sites on shallow trees. `WALRUS` therefore forms the parameter-free conjugate Gamma$(1,1)$–Poisson Empirical Bayes posterior mean synonymous rate

$$\tilde{\alpha}_c = \frac{1 + \hat{\alpha}_c T_{\text{syn},c}}{1 + T_{\text{syn},c}}, \tag{S19}$$

where $T_{\mathrm{syn},c}$ is the expected synonymous tree length of the non-gap taxa at site $c$, and evaluates the background tree cavity $(\bar{\mathbf{U}}_{b,c}, \bar{\mathbf{L}}_{b,c})$ at the purifying-to-neutral baseline $\hat{\omega}_{\mathrm{cav},c} = \min(1, \hat{\omega}_c)$ so single-branch episodic substitutions or errors do not diffuse across internal branches. Stacking $\bar{\mathbf{U}}_{b,c}$ and $\bar{\mathbf{L}}_{b,c}$ across all $C$ sites into $61 \times C$ matrices $U_b$ and $L_b$, we project both into the $(k, g)$ eigenbasis via two $61 \times 61 \times C$ BLAS matrix–matrix multiplications ('DGEMM') and form their Hadamard product $H_b^{(k,g)} = (R_{k,g}^{\top} U_b) \odot (L_{k,g} L_b) \in \mathbb{R}^{61 \times C}$. Because $R_{k,g}$ and $L_{k,g}$ are independent of branch length, the exact bilinear likelihood at site $c$'s scaled branch length $\tau_{b,c} = \tilde{\alpha}_c t_{b,c}$ reduces to a 61-element vector dot product, yielding the dimensionless $C \times B \times 16$ sufficiency tensor:

$$\bar{\mathbf{U}}_{b,c}^{\top} P_b(\tau_{b,c} \mid \omega_g) \bar{\mathbf{L}}_{b,c} = \sum_{k=0}^{K_{\mathrm{reg}}-1} w_{b,k} \big(H_{b,:,c}^{(k,g)}\big)^{\top} \exp\big(\tau_{b,c} \boldsymbol{\lambda}_{k,g}\big), \qquad W[c][b][g] = \frac{\bar{\mathbf{U}}_{b,c}^{\top} P_b(\tau_{b,c} \mid \omega_g) \bar{\mathbf{L}}_{b,c}}{\bar{\mathbf{U}}_{b,c}^{\top} P_b(\tau_{b,c} \mid \hat{\omega}_{\mathrm{cav},c}) \bar{\mathbf{L}}_{b,c}}. \tag{S20}$$

3. *Erlang-3 minimum-duration Baum–Welch error-sink HMM and masked cavity recalculation.* To prevent local sequencing errors, unmasked frameshifts, or non-homologous overhangs from masquerading as episodic diversifying selection, each branch $b$ is scanned along $c = 1, \ldots, C$ by an Erlang-3 minimum-duration hidden Markov model ($0 \xrightarrow{\epsilon_b} 1_1 \xrightarrow{1} 1_2 \xrightarrow{1} 1_3 \looparrowright \rho_b$) with transition parameters $(\hat{\epsilon}_b, \hat{\rho}_b)$ estimated via Baum–Welch expectation–maximization. The error-tract states $(1_1, 1_2, 1_3)$ emit from the off-diagonal random-jump kernel $P_{ij}^{\mathrm{sink}} = \mathbb{I}(i \neq j)\pi_j/(1 - \pi_i)$ along the 1D cavity maximum-likelihood homologous branch transition $P_b^{\mathrm{sink}}(\hat{t}_b^{\mathrm{hom}}) = (1 - e^{-\hat{t}_b^{\mathrm{hom}}}) P^{\mathrm{sink}}$. Crucially, both the error-sink numerator and the 16-point Empirical Bayes biological codon denominator are normalized on the identical $W[c][b][g]$ baseline scale $(\bar{\mathbf{U}}_{b,c}^{\top} P_b(\tau_{b,c} \mid \hat{\omega}_{\mathrm{cav},c}) \bar{\mathbf{L}}_{b,c})$:

$$W_{\mathrm{sink}}[c][b] = \frac{\bar{\mathbf{U}}_{b,c}^{\top} P_b^{\mathrm{sink}}(\hat{t}_b^{\mathrm{hom}}) \bar{\mathbf{L}}_{b,c} \;/\; \bar{\mathbf{U}}_{b,c}^{\top} P_b(\tau_{b,c} \mid \hat{\omega}_{\mathrm{cav},c}) \bar{\mathbf{L}}_{b,c}}{\max_{g \in \mathcal{G}} \; \hat{\pi}_{\mathrm{grid}}(g)\, W[c][b][g]}, \tag{S21}$$

where $\hat{\pi}_{\mathrm{grid}}(g)$ is the Laplace-smoothed empirical histogram of site MLEs $\hat{\omega}_c$ across the gene (with $\hat{\pi}_{\mathrm{grid}}(g) = 1$ for $\omega_g \leq 1$ and for site $c$'s baseline $\hat{\omega}_c$). Because entering the error sink requires traversing at least three consecutive mismatched codons ($1_1 \to 1_2 \to 1_3$) with $P_{ii}^{\mathrm{sink}} = 0$, isolated 1- or 2-codon substitutions cannot enter the error sink, and genuine 1-nucleotide adaptive epitopes are explained by $\omega_g > 1$ ($W_{\mathrm{sink}}[c][b] < 1$). Error-sink posteriors $\gamma_{c,b}$ on branch $b$ are retained if and only if the converged HMM log-likelihood $\ln Z_b(\hat{\epsilon}_b, \hat{\rho}_b)$ passes the Schwarz BIC gate $2 \ln Z_b(\hat{\epsilon}_b, \hat{\rho}_b) > 2 \ln(B \cdot C)$. When triggered, $t_b$ is updated via 1D golden-section maximization of the $(1 - \gamma_{c,b})$-weighted cavity log-likelihood, and the Inside–Outside cavities at affected sites are recomputed with the corrupted branch's upward message attenuated toward the missing-data vector $\mathbf{1}$ as $(1 - \gamma_{c,b}) P_b \mathbf{L}_{b,c} + \gamma_{c,b} \mathbf{1}$, preventing error leakage into sibling and ancestor Outside vectors while preserving valid evolutionary signal on sister lineages.

##### 1.6 Stage 5: Multi-Scale Active-Set Polish, Structural Segmentation, and Simes Omnibus Synthesis

From the cached cavity vectors $(\bar{\mathbf{U}}_{b,c}, \bar{\mathbf{L}}_{b,c})$ and sufficiency tensor $W[c][b][g]$, Stage 5 executes a unified screen-and-polish across all evolutionary scales without repeating global tree traversals:

1. *Gene-wide episodic diversifying selection with error sink and boundary-consistency guards (`BUSTED-E`).* `WALRUS` fits a four-component branch-site mixture over $\omega$: three biological regimes ($\omega_1 \leq \omega_2 \leq 1 \leq \omega_3$ with weights $w_1, w_2, w_3$) plus a dedicated alignment/multi-hit error-sink

regime fixed at $\omega_{\text{err}} = 100$ with weight $w_{\text{err}} \leq 2.5\%$ ($\sum_{m=1}^{3} w_m + w_{\text{err}} = 1$). Using the cavity rank-1 approximation on $W[c][b][g]$, the conditional likelihood ratio at site $c$ and branch $b$ under mixture component $m$ is $W_{c,b}(\omega_m)$, and the gene-wide log-likelihood is optimized via L-BFGS-B and Nelder–Mead simplex polish against the constrained null $\omega_3 = 1$. Residual single-branch multi-nucleotide alignment artifacts are absorbed into $\omega_{\text{err}} = 100$ rather than inflating $\omega_3$. To prevent boundary-splitting optimization pathologies on degenerate or flat neutral likelihood surfaces (Davies, 1977), Stage 5 enforces three boundary-consistency conditions plus a false-negative rescue seed inside the integrated pipeline:

- *Minimum effective codon support:* The unconstrained positive-selection class must represent at least $w_3 C \geq 0.50$ effective codons and $w_3 \geq 0.15\%$ weight, preventing sub-codon numerical dust from generating positive LRTs.
- *Flat-neutral boundary-split guard:* When the inferred purifying weight vanishes ($w_1 < 0.52$, diagnostic of the degenerate $\omega \equiv 1$ boundary where splitting a point mass at $\omega = 1$ into $\omega_1 < 1$ and $\omega_3 > 1$ fits Poisson branch variance) and zero individual codon sites reach Simes omnibus $q \leq 0.10$, the gene-wide LRT is conservatively damped (LRT $\leftarrow 0, p \leftarrow 1.0$).
- *Moderate-tail curvature guard:* When the unconstrained tail sits at moderate intensity ($1 < \omega_3 \leq 2.5$) with zero zero-synonymous diversifying sites ($\tilde{\alpha}_c < 0.25, \hat{\omega}_c > 1.5$) and at most three Simes $q \leq 0.10$ sites (diagnostic of U-shaped Beta$(0.5, 0.5)$ density curvature near $\omega = 1$ in PAML `M7`), the gene-wide LRT is likewise damped (LRT $\leftarrow 0, p \leftarrow 1.0$). Across 11,190 empirical avian proteins, these boundary guards trigger on only 0.12% (13/11,190) and 0.00% (0/11,190) of orthogroups, whereas on synthetic `all0is1` and `M7` null simulations they fire on 12.3% and 21.8% of replicates (vs. 1.3% and 5.0% of positive controls).
- *Boundary-absorption false-negative rescue:* Conversely, when the initial 3-class grid initialization traps $w_3 < 1.2\%$ near zero on a conserved gene ($w_1 + w_2 > 0.25$) with zero Simes diversifying sites, `WALRUS` re-optimizes the unconstrained mixture from a seeded positive-tail initialization ($w_3^{(0)} = 0.04, \omega_3^{(0)} = \max(3.5, \hat{\omega}_3)$) and retains the rescued fit if and only if it strictly improves $\ln L$ and yields $p \leq 0.05$.

2. *BIC-penalized 1D change-point protein domain segmentation and Synonymous Constraint Islands (SCIs).* Rather than relying on external structural annotations, `WALRUS` partitions the coding sequence into contiguous selective domains via exact $O(C^2)$ BIC-penalized 1D change-point dynamic programming. At each codon $c$, we form the standardized 4-vector $\mathbf{z}_c = (\ln(\hat{\omega}_c + 0.05),\ \tilde{\alpha}_c,\ 1 - o_c,\ N_{\text{eff}}(c))$ and minimize the penalized within-segment Gaussian sum of squares across partitions $0 = \tau_0 < \tau_1 < \cdots < \tau_D = C$ subject to minimum domain length $L_{\min} = 10$ codons and change-point penalty $\lambda_{\text{BIC}} = 2.5 \ln C$:

$$\mathcal{J}(\tau_0, \ldots, \tau_D) = \sum_{d=1}^{D} \sum_{c=\tau_{d-1}+1}^{\tau_d} \left\| \mathbf{z}_c - \bar{\mathbf{z}}_{(\tau_{d-1}, \tau_d]} \right\|_2^2 + (D-1)\, \lambda_{\text{BIC}}. \tag{S22}$$

Each resulting segment $d$ is classified by its mean selective profile ($\bar{\omega}_d, \overline{\text{gap}}_d$) as *Purifying Core* ($\bar{\omega}_d < 0.25$ and $\overline{\text{gap}}_d < 0.15$), *Hypervariable / Intrinsically Disordered Region (IDR)* ($\bar{\omega}_d > 0.75$ or $\overline{\text{gap}}_d \geq 0.25$), or *Permissive Surface* (otherwise). Simultaneously, a sliding-window scan ($W_{\text{SCI}} = 15$ codons) over the Empirical Bayes synonymous rates $\tilde{\alpha}_c$ flags *Synonymous Constraint Islands* (overlapping regulatory elements, splicing enhancers, or RNA secondary structures) wherever the window mean $\bar{\alpha}_w \leq 0.35\, \bar{\alpha}_{\text{gene}}$ and the standardized suppression score $z_w = (\bar{\alpha}_{\text{gene}} - \bar{\alpha}_w)/(\text{SD}(\tilde{\alpha})/\sqrt{W_{\text{SCI}}}) \geq 3.0$.

3. *Autonomous BIC clade scan with tree-wide Holm multiplicity control (`RELAX` and `Contrast-FEL`).* When user-defined foreground branch tags are supplied, `WALRUS` evaluates the designated foreground versus background partitions directly. In fully unguided mode, `WALRUS` scans all $B_{\text{int}}$ internal clade splits $b$ satisfying $\min(|\mathcal{C}_b|, N - |\mathcal{C}_b|) \geq \max(3, \lceil 0.10\, N \rceil)$. For each candidate clade $\mathcal{C}_b$, it fits a two-partition selection-intensity parameter $K_b$ ($\omega_{\text{test}} = \omega_{\text{ref}}^{K_b}$) on $W[c][b][g]$ and computes $\Delta\text{BIC}_b = 2(\ln L(K_b) - \ln L(K_b = 1)) - \ln(B \cdot C)$. Splits achieving $\Delta\text{BIC}_b \geq 10$ undergo full `RELAX` ($K_b > 1$ intensification vs. $K_b < 1$ relaxation) and site-level `Contrast-FEL` ($\beta_{\text{test},c} \neq \beta_{\text{ref},c}$) continuous optimization. Because scanning $B_{\text{int}}$ internal splits constitutes multiple hypothesis testing across the phylogeny, all unguided clade-scan $p$-values are adjusted via the step-down Holm–Bonferroni procedure across all $B_{\text{int}}$ tested splits:

$$p_{\text{Holm},(m)} = \min\Big(1, \max_{1 \leq j \leq m} (B_{\text{int}} - j + 1)\, p_{(j)}\Big), \tag{S23}$$

controlling the family-wise error rate (FWER) of autonomous regime discovery at $\alpha = 0.05$.

4. *Branch-level adaptive bursts (`aBSREL`) versus Lineage Loss-of-Constraint (LoC).* On each branch $b$, `WALRUS` screens for episodic diversifying bursts (`aBSREL` 2-rate mixture $\omega_{b,1} \leq 1 < \omega_{b,2}$ on $W[c][b][g]$, followed by continuous golden-section polish on branches with $p_{\text{screen}} < 0.25$ and Holm–Bonferroni correction across all $B$ branches). To distinguish genuine adaptive bursts from lineage-specific pseudogenization or relaxed purifying selection, `WALRUS` defines the gene's conserved structural core $\mathcal{S}_{\text{core}} = \{c : \hat{\omega}_c < 0.15 \text{ and } N_{\text{eff}}(c) \leq 1.5\}$. For each branch $b$, let $p_{b,c}^{\text{NS}}$ denote the null posterior probability of a nonsynonymous substitution at core site $c \in \mathcal{S}_{\text{core}}$ under the gene-wide purifying baseline, $\lambda_{b,\text{core}} = \sum_{c \in \mathcal{S}_{\text{core}}} p_{b,c}^{\text{NS}}$ be the expected core hit count, and $K_{b,\text{core}}$ be the observed posterior core nonsynonymous count. `WALRUS` computes the Poisson–Binomial tail probability $\mathbb{P}(X \geq K_{b,\text{core}} \mid \lambda_{b,\text{core}})$ and the Lineage Loss-of-Constraint Bayes factor $\text{BF}_{\text{LoC}}(b)$ contrasting invasion of $\mathcal{S}_{\text{core}}$ ($\omega_{\text{core},b} = 1$) against core conservation ($\omega_{\text{core},b} = \hat{\omega}_{\text{core}}$).

5. *Codon-level active-set polish (`FEL`/`dFEL`, `MEME`, `PRIME`), amino-acid mode classification, and Simes omnibus FDR.* At every codon site $c \in \{1, \ldots, C\}$, initial grid screening on $W[c][b][g]$ identifies candidate sites with $p_{\text{screen}} < 0.25$, which then undergo exact continuous active-set optimization using the cached cavity vectors $(\bar{\mathbf{U}}_{b,c}, \bar{\mathbf{L}}_{b,c})$:

   - *Pervasive selection (`FEL`/ `dFEL`):* 1D Brent optimization of $\beta_c = \omega_c \tilde{\alpha}_c$ against the neutral null $\beta_c = \tilde{\alpha}_c$ under the stationary ($K_{\text{reg}} = 1$) or Stage 3 directional ($K_{\text{reg}} = 3$) rate kernel (1-df $\chi_1^2$ LRT).
   - *Episodic diversifying selection (`MEME`):* Continuous optimization of the two-class branch mixture $\omega_c^- \in [0, 1]$ (weight $p_c^-$) and $\omega_c^+ \geq 1$ (weight $1 - p_c^-$) against the constrained null $\omega_c^+ = 1$, yielding both the site LRT ($0.33\chi_0^2 + 0.30\chi_1^2 + 0.37\chi_2^2$) and empirical Bayes branch-specific posterior probabilities $\text{PP}_{b,c}(\omega^+ > 1)$ and Bayes factors $\text{BF}_{b,c}$.
   - *Biochemical property-informed selection (`PRIME`):* Damped Newton–Raphson optimization of the $P = 5$ Atchley/Conant–Stadler amino-acid property slopes $\boldsymbol{\lambda}_c = (\lambda_{c,1}, \ldots, \lambda_{c,5})^\top \in \mathbb{R}^5$ governing nonsynonymous exchangeabilities $\omega_c(x, y) = \omega_{0,c} \exp(-\sum_{p=1}^5 \lambda_{c,p} |v_p(A(x)) - v_p(A(y))|)$. From the analytical Fréchet score vector $\mathbf{U}_c = \nabla_{\boldsymbol{\lambda}} \ell_c(\mathbf{0}) \in \mathbb{R}^5$ and observed Fisher information $\mathcal{I}_c \in \mathbb{R}^{5 \times 5}$ evaluated on the cavity sufficient statistics, `WALRUS` computes both the omnibus 5-df statistic $T_c^{\text{PRIME}} = \mathbf{U}_c^\top (\mathcal{I}_c + \tau I_5)^{-1} \mathbf{U}_c$ and individual property $z$-scores ($\lambda_{c,p} > 0$ property conservation vs. $\lambda_{c,p} < 0$ radical property diversification).

For every site with expected nonsynonymous substitutions $M_c = \sum_b \mathbb{E}[N_{b,c}^{\text{NS}} \mid D] \geq 2$, `WALRUS` computes the effective amino-acid alphabet size $N_{\text{eff}}(c) = \exp(H(\hat{\boldsymbol{\pi}}_c^{\text{AA}}))$ relative to the neutral codon-branching expectation $\mathbb{E}_0[N_{\text{eff}} \mid M_c]$ and classifies the substitution trajectory into five mechanistic modes: *Toggle* ($A \leftrightarrow B$, top-2 residues accounting for $\geq 85\%$ of nonsynonymous changes with $\geq 1$ bidirectional reversion), *Convergent* ($\rightarrow A$, $\geq 3$ independent origins of target residue $A$ across non-sister branches), *Sweep* (single-branch fixation onto a derived clade of $\geq 4$ leaves), *Focused Exchange* ($N_{\text{eff}}(c) < 0.75\,\mathbb{E}_0[N_{\text{eff}} \mid M_c]$), or *Broadly Diversifying* (unconstrained turnover across $\geq 4$ amino acids), alongside 'CpG' (`CGN/NCG`) and 'APOBEC3' (`TCN` → `TTN/TGN`) mutational context fractions. Finally, site-level $p$-values across the $K_{\text{tests}} = 3$ active diversifying test families ($p_{c,1}, \ldots, p_{c,K_{\text{tests}}}$ ordered as $p_{c,(1)} \leq \cdots \leq p_{c,(K_{\text{tests}})}$) are combined via Simes' (1986) omnibus procedure (Simes, 1986),

$$p_{c,\text{Simes}} = \min_{1 \leq j \leq K_{\text{tests}}} \left\{ \frac{K_{\text{tests}}\, p_{c,(j)}}{j} \right\}, \tag{S24}$$

which remains valid under positive regression dependency among test statistics evaluated on the same site, followed by Benjamini–Hochberg step-up control ($q_c \leq 0.10$) across all $C$ codons.

### 2 Supplementary Empirical Analyses: Avian Exome Realignment Forensics and Covariate Stratification

To complement Section 4.2 of the main text, we examined how `WALRUS`'s Stage 0 overhang trimming and Stage 1–2 phylogeny-aware codon realignment depend on gene-level structural and evolutionary covariates across the 11,189 paired orthogroups from Shultz and Sackton (2019), and visualized the codon-level alignment restorations for the three conserved housekeeping false-positive case studies (*AP3B2*, *NOC2L*, and *FBN2*; Table 5 of the main text).

#### 2.1 Data Governance and Biosafety Compliance

All empirical sequence datasets analyzed in this study were retrieved from public, de-identified nucleotide repositories (GenBank and the HyPhy benchmark archive) or generated via in silico Markov sequence simulation. No human subjects, clinical patient records, or personally identifiable health data were accessed. This computational work involved no wet-laboratory experiments, pathogen culturing, or gain-of-function modifications.

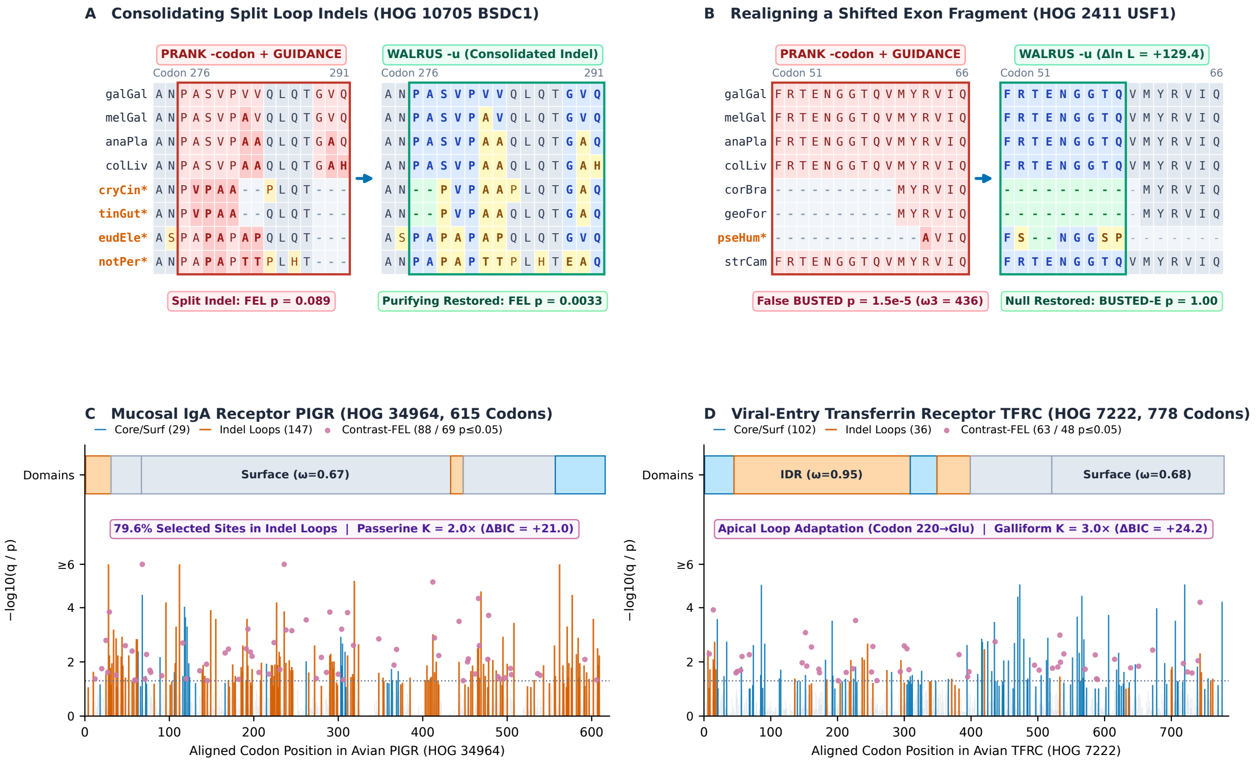

**Figure S5.** *Codon-level alignment forensics comparing published progressive alignments against* WALRUS -u *co-estimated alignments for three conserved avian housekeeping genes from* Shultz and Sackton *(2019) (Table 5 of the main text).* Each panel displays the published alignment window (left, where staggered indel placement or unmasked frameshifts create localized clusters of apparent nonsynonymous substitutions and drive standalone BUSTED $p < 10^{-7}$) alongside the corresponding WALRUS -u restored alignment window (right, where Stage 0–2 phase repair, overhang trimming, and cavity Pair-HMM realignment restore homologous codon columns and yield $p = 1.00$). Panel A: *AP3B2* (Orthogroup 6798, clathrin adaptor complex $\beta$2 subunit). Panel B: *NOC2L* (Orthogroup 34964, nucleolar complex-associated 2 homolog). Panel C: *FBN2* (Orthogroup 7222, fibrillin-2 extracellular matrix structural glycoprotein).

**Table S3.** *Stage 0 empirical summary quantities, genetic-code neutral reference constants, closed-form moment estimators, and representative empirical estimates for Stage 1 Phase 2 codon profile alignment.* Counts $C_\bullet$ are tallied across banded gapped-DP alignments of Stage 0 homologous pairs ($\geq 35\%$ amino-acid identity) with unit Laplace pseudocounts. Here $\Delta(x, y) \subseteq \{1, 2, 3\}$ denotes the differing nucleotide positions between codons $x$ and $y$, and $A(x)$ is the encoded amino acid.

| Symbol | Definition / Estimator | Biological Role & Universal-Code / Empirical Values |
|---|---|---|
| *A. Site-level divergences and five-codon sliding-window stratification ($W_k = [k-2, k+2]$).* | | |
| $d_{\mathrm{nt}}(k)$, $d_{\mathrm{AA}}(k)$ | $\dfrac{\lvert\Delta(c_1, c_2)\rvert}{3} \in \{0, \frac{1}{3}, \frac{2}{3}, 1\}, \quad d_{\mathrm{AA}}\big(A(c_1), A(c_2)\big) \in [0, 1]$ | Per-site nucleotide divergence and Grantham–Miyata amino-acid distance (0 if synonymous; e.g. Leu ↔ Ile 0.04, Glu ↔ Lys 0.50). |
| $\mathcal{C}_{\mathrm{loop}}$ vs. $\mathcal{C}_{\mathrm{core}}$ | $k \in \mathcal{C}_{\mathrm{loop}} \iff \bar{d}_{\mathrm{AA}}^{W_k} > \bar{d}_{\mathrm{AA}} \quad (\mathcal{C}_{\mathrm{core}}$ otherwise) | Assigns site $k$ to surface loop stratum if its 5-codon window amino-acid divergence exceeds the pair mean $\bar{d}_{\mathrm{AA}}$. |
| $r(k)$ | $\dfrac{1}{2}\left(\dfrac{d_{\mathrm{nt}}(k)}{\bar{d}_{\mathrm{nt}}} + \dfrac{d_{\mathrm{AA}}(k)}{\bar{d}_{\mathrm{AA}}}\right), \quad \dfrac{1}{M}\sum_{k=1}^{M} r(k) = 1$ | Unit-mean relative evolutionary rate at site $k$ ($\bar{r}_{\mathrm{core}} \in [0.18, 0.53]$ vs. $\bar{r}_{\mathrm{loop}} \in [1.65, 2.58]$ across benchmarks). |
| $\hat{w}_{\mathrm{loop}}$, $\hat{\rho}_{\mathrm{loop/core}}$ | $\hat{w}_{\mathrm{loop}} = \dfrac{\lvert\mathcal{C}_{\mathrm{loop}}\rvert}{\lvert\mathcal{C}_{\mathrm{core}}\rvert + \lvert\mathcal{C}_{\mathrm{loop}}\rvert}, \quad \hat{\rho}_{\mathrm{loop/core}} = \dfrac{\bar{r}_{\mathrm{loop}}}{\bar{r}_{\mathrm{core}}}$ | Loop mixture weight ($\hat{w}_{\mathrm{loop}} \in [0.28, 0.42]$) and loop-to-core branch rate multiplier ($\hat{\rho}_{\mathrm{loop/core}} \in [3.10, 8.42]$). |
| *B. Genetic-code neutral reference constants (unordered sense-codon pairs $\{x, y\}$, shown for Universal code $S = 61$).* | | |
| $N_1$, $N_2$, $N_3$ | $\big\lvert\{\{x, y\} : \lvert\Delta(x, y)\rvert = h\}\big\rvert \quad (h \in \{1, 2, 3\})$ | Sense-codon pairs at $h = 1, 2, 3$ diffs: $N_1 = 263$, $N_2 = 784$ ($N_1/N_2 = 0.336$), $N_3 = 783$ ($N_1/N_3 = 0.336$). |
| $N_{\mathrm{syn}}^{(1)}$, $N_{\mathrm{ns}}^{(1)}$ | $\big\lvert\{\{x, y\} : \lvert\Delta\rvert = 1,\ A(x) = A(y)\}\big\rvert,\ N_1 - N_{\mathrm{ns}}^{(1)}$ | 1-nt synonymous and nonsynonymous pair counts: $N_{\mathrm{syn}}^{(1)} = 67$, $N_{\mathrm{ns}}^{(1)} = 196$ ($N_{\mathrm{syn}}^{(1)}/N_{\mathrm{ns}}^{(1)} = 0.342$). |
| $\mu_0$, $\sigma_0^2$ | $\mathbb{E}_0[d_{\mathrm{AA}}], \quad \mathrm{Var}_0(d_{\mathrm{AA}})$ over $N_{\mathrm{ns}}^{(1)}$ pairs | Neutral mean $\mu_0 = 0.302$ and variance $\sigma_0^2 = 0.0301$ ($\sigma_0 = 0.174$) of $d_{\mathrm{AA}}$ across the 196 1-nt nonsynonymous pairs. |
| *C. Stratum-specific substitution moment estimators ($m \in \{core, loop\}$) and nucleotide exchangeabilities.* | | |
| $\hat{\theta}_{CT}$, $\hat{\theta}_{\mathrm{TV}}$ | $\hat{\theta}_{CT} = \dfrac{C_{CT}}{C_{AG}}, \quad \hat{\theta}_{\mathrm{TV}} = \dfrac{C_{\mathrm{TV}}}{C_{AG} + C_{CT}}$ | Pyrimidine-to-purine transition ratio ($\hat{\theta}_{CT} \in [0.55, 2.50]$) and transversion ratio ($\hat{\theta}_{\mathrm{TV}} \in [0.37, 0.85]$). |
| $\hat{\omega}_m$ | $\dfrac{C_{\mathrm{ns},m}^{(1)}}{C_{\mathrm{syn},m}^{(1)}} \cdot \dfrac{N_{\mathrm{syn}}^{(1)}}{N_{\mathrm{ns}}^{(1)}}$ | Nonsynonymous-to-synonymous rate ratio ($\hat{\omega}_{\mathrm{core}} \in [0.06, 0.40]$ vs. $\hat{\omega}_{\mathrm{loop}} \in [0.18, 1.50]$). |
| $\hat{\lambda}_{\mathrm{AA},m}$ | $\max\left(0, \dfrac{\mu_0 - \bar{d}_{\mathrm{AA},m}^{\mathrm{obs}}}{\sigma_0^2}\right)$ | Physicochemical selectivity slope ($\hat{\lambda}_{\mathrm{AA,core}} \in [2.70, 3.50]$ vs. $\hat{\lambda}_{\mathrm{AA,loop}} \in [0.00, 3.03]$). |
| $\hat{\delta}_{2\mathrm{nt},m}$, $\hat{\psi}_{3\mathrm{nt},m}$ | $\dfrac{C_{2\mathrm{nt},m}}{C_{1\mathrm{nt},m}} \cdot \dfrac{N_1}{N_2}, \quad \dfrac{C_{3\mathrm{nt},m}}{C_{1\mathrm{nt},m}} \cdot \dfrac{N_1}{N_3}$ | Opportunity-normalized 2-nt and 3-nt multi-hit rates ($\hat{\delta}_{2\mathrm{nt}}$ : 0.06–0.13 → 0.06–0.26; $\hat{\psi}_{3\mathrm{nt}}$ : 0.01–0.03 → 0.01–0.09). |

*D. Concrete Stage 0 parameter estimates and scaled instantaneous rates ($\hat{Q}_{core} \to \hat{Q}_{loop}$) across six empirical coding datasets.*

| Dataset ($N$) | $\hat{w}_{\mathrm{loop}}$ | $\hat{\rho}_{\mathrm{loop/core}}$ | $\hat{\theta}_{CT}/\hat{\theta}_{\mathrm{TV}}$ | $\hat{\omega}_{\mathrm{core}} \to \hat{\omega}_{\mathrm{loop}}$ | $\bar{d}_{\mathrm{AA,c}}^{\mathrm{obs}} \to \bar{d}_{\mathrm{AA,l}}^{\mathrm{obs}}$ | $\hat{\lambda}_{\mathrm{AA,c}} \to \hat{\lambda}_{\mathrm{AA,l}}$ | $\hat{\delta}_{2\mathrm{nt,c}} \to \hat{\delta}_{2\mathrm{nt,l}}$ | $\hat{Q}$(`CTG`→`CAG`) | $\hat{Q}$ |
|---|---|---|---|---|---|---|---|---|---|
| *D. adh* (23) | 0.33 | 4.90× | 2.50/0.81 | 0.06 → 0.18 | 0.12 → 0.23 | 3.50 → 2.33 | 0.06 → 0.15 | 0.02 → 0.06 | 0 |
| *OPN1SW* (33) | 0.28 | 7.03× | 1.56/0.37 | 0.06 → 0.58 | 0.13 → 0.28 | 3.50 → 0.85 | 0.06 → 0.06 | 0.01 → 0.09 | 0 |
| *Mamm. IL10* (16) | 0.38 | 5.07× | 0.71/0.60 | 0.11 → 0.79 | 0.15 → 0.28 | 3.50 → 0.66 | 0.06 → 0.13 | 0.06 → 0.33 | 0 |
| *Mamm. IFITM3* (13) | 0.38 | 4.90× | 1.72/0.85 | 0.11 → 0.51 | 0.11 → 0.21 | 3.50 → 3.03 | 0.07 → 0.18 | 0.03 → 0.08 | 0 |
| *Mamm. CD2* (10) | 0.42 | 3.10× | 0.76/0.79 | 0.32 → 1.16 | 0.22 → 0.32 | 2.70 → 0.00 | 0.13 → 0.26 | 0.17 → 0.48 | 0 |
| *APOBEC3H* (17) | 0.38 | 8.42× | 0.55/0.62 | 0.40 → 1.50 | 0.15 → 0.40 | 3.50 → 0.00 | 0.06 → 0.09 | 0.12 → 0.45 | 0 |

**Table S4.** *Stratification of* `WALRUS` *Stage 0–4 alignment restructuring, quarantine, POA boundary masking, error-sink absorption, and false-positive rescue across evolutionary and structural covariates in* 11,183 *avian orthogroups (Shultz and Sackton, 2019).* Columns report the number of orthogroups ($n$), the percentage of genes with altered codon length ($\Delta C \neq 0$), mean codon length change ($\overline{\Delta C}$), percentage of genes with $\geq 1$ sequence quarantined by Stage 0 (Quar.%), mean percentage of columns inside Stage 1 Phase 3 indel realignment windows (IndelWin%), percentage of genes with $\geq 1$ false-positive site eliminated by Stage 4 POA column marginalization (POA FP%), percentage of genes with active Stage 5 error-sink absorption ($w_{\mathrm{err}} > 0.01\%$), published standalone `BUSTED` positive rate ($q \leq 0.05$), `WALRUS -u` positive rate ($q \leq 0.05$), relative fold reduction in positive calls, and median single-node runtime.

| Covariate Axis | Stratum | $n$ | $\Delta C \neq 0$ | Mean $\Delta C$ | Quar.% | IndelWin% | POA FP% | $w_{\mathrm{err}} > 0\%$ | Publ. `BUSTED` | `WALRUS -u` | Fold Red. | Med. Time |
|---|---|---|---|---|---|---|---|---|---|---|---|---|
| Coding Length ($C_{\mathrm{pub}}$) | Q1: 34–204 c | 2,232 | 12.0% | +0.18 | 4.5% | 34.4% | 3.6% | 18.1% | 32.6% | 5.6% | 5.87× | 43.9 s |
| | Q2: 205–321 c | 2,220 | 17.4% | +0.31 | 3.2% | 38.0% | 9.4% | 20.6% | 46.4% | 9.0% | 5.14× | 88.5 s |
| | Q3: 322–459 c | 2,247 | 22.0% | +0.52 | 2.5% | 37.5% | 12.7% | 21.1% | 53.7% | 9.8% | 5.46× | 143.8 s |
| | Q4: 460–710 c | 2,246 | 30.5% | +0.69 | 1.9% | 37.7% | 18.0% | 21.5% | 65.0% | 14.2% | 4.56× | 218.0 s |
| | Q5: 711–5,117 c | 2,238 | 43.8% | +1.36 | 2.4% | 37.3% | 34.0% | 21.4% | 79.2% | 21.7% | 3.66× | 467.7 s |
| Tree Length ($T_{\mathrm{M0}}$) | Q1: 0.25–2.87 | 2,796 | 12.7% | +0.31 | 1.6% | 23.6% | 4.4% | 12.0% | 44.1% | 2.3% | 19.27× | 79.8 s |
| | Q2: 2.87–3.87 | 2,795 | 21.1% | +0.49 | 1.6% | 31.0% | 11.7% | 17.9% | 53.3% | 7.5% | 7.12× | 139.2 s |
| | Q3: 3.87–5.20 | 2,796 | 29.4% | +0.67 | 1.9% | 39.5% | 19.2% | 21.4% | 59.4% | 14.7% | 4.04× | 180.9 s |
| | Q4: 5.20–129.10 | 2,796 | 37.4% | +0.98 | 6.6% | 53.8% | 26.9% | 30.9% | 64.9% | 23.8% | 2.72× | 200.4 s |
| Selective Rate ($\omega_{\mathrm{M0}}$) | Q1: 0.000–0.045 | 2,796 | 17.8% | +0.54 | 3.6% | 30.7% | 3.5% | 13.0% | 41.6% | 1.3% | 32.31× | 94.4 s |
| | Q2: 0.045–0.102 | 2,795 | 23.8% | +0.67 | 2.8% | 35.3% | 9.4% | 19.6% | 49.4% | 5.0% | 9.93× | 141.7 s |
| | Q3: 0.102–0.205 | 2,796 | 26.9% | +0.57 | 2.4% | 36.9% | 15.2% | 22.2% | 55.9% | 7.7% | 7.24× | 157.4 s |
| | Q4: 0.205–2.106 | 2,796 | 32.2% | +0.67 | 2.9% | 45.1% | 34.2% | 27.4% | 74.7% | 34.3% | 2.18× | 197.0 s |
| Indel Rate (events/100 c) | Q1: 0.0–4.0/100 c | 2,795 | 7.4% | +0.10 | 1.3% | 16.7% | 3.9% | 7.9% | 39.2% | 5.6% | 7.03× | 115.0 s |
| | Q2: 4.0–6.4/100 c | 2,795 | 19.1% | +0.38 | 2.5% | 29.3% | 11.3% | 14.1% | 53.1% | 9.7% | 5.48× | 147.0 s |
| | Q3: 6.4–10.0/100 c | 2,796 | 29.0% | +0.73 | 3.5% | 41.1% | 19.1% | 24.6% | 59.3% | 14.6% | 4.07× | 152.1 s |
| | Q4: 10.0–73.1/100 c | 2,797 | 45.1% | +1.24 | 4.4% | 60.8% | 28.0% | 35.5% | 70.0% | 18.4% | 3.79× | 160.4 s |
